# Atom level enzyme active site scaffolding using RFdiffusion2

**DOI:** 10.1101/2025.04.09.648075

**Authors:** Woody Ahern, Jason Yim, Doug Tischer, Saman Salike, Seth M. Woodbury, Donghyo Kim, Indrek Kalvet, Yakov Kipnis, Brian Coventry, Han Raut Altae-Tran, Magnus Bauer, Regina Barzilay, Tommi S. Jaakkola, Rohith Krishna, David Baker

## Abstract

*De novo* enzyme design starts from ideal active site descriptions consisting of constellations of catalytic residue functional groups around reaction transition state(s), and seeks to generate protein structures that can accurately hold the site in place. Highly active enzymes have been designed starting from such descriptions using the generative AI method RFdiffusion [1–3], but there are two current methodological limitations. First, the geometry of the active site can only be specified at the residue level, so for each catalytic residue functional group placed around the reaction transition state, the possible locations of the residue backbone must be enumerated by building side chain rotamers back from the functional group. Second, the location of the catalytic residues along the sequence must be specified in advance, which considerably limits the space of solutions which can be sampled. Here we describe a new deep generative method, Rosetta Fold diffusion 2 (RFdiffusion2), that solves both problems, enabling enzymes to be designed from sequence agnostic descriptions of functional group locations without inverse rotamer generation. We first evaluate RFdiffusion2 on an *in silico* enzyme design benchmark of 41 diverse active sites and find that it is able to successfully build proteins scaffolding all 41 sites, compared to 16/41 with prior state-of-the-art deep learning methods. Next, we design enzymes around three diverse catalytic sites and characterize the designs experimentally; in each case we identify active catalysts in testing less than 96 sequences. RFdiffusion2 demonstrates the potential of atomic resolution generative models for the design of *de novo* enzymes directly from their reaction mechanisms.

## 1 Main

A grand challenge in *de novo* protein design is the generation of enzymes that catalyze novel reactions. *De novo* enzyme design starts from a detailed description of an active site composition and geometry predicted to catalyze the reaction of interest. This active site description, called a theozyme, describes placement of protein functional groups around the reaction transition state(s) and any reaction cofactors [4, 5]. The *de novo* enzyme design task is to generate protein scaffolds that accommodate such theozymes. Pre-deep learning methods such as RosettaMatch searched through sets of already existing native or designed [6] scaffolds for possible placements of the catalytic residues. While many enzymes were designed using this approach, it was restricted to theozyme geometries that could be matched to the input scaffold set [7]. Advances in deep learning with diffusion models [8–11] have removed the need for scaffold libraries for many substructure scaffolding tasks by directly sampling diverse proteins containing the desired substructure (motif) through a technique known as motif scaffolding [12–15]. However, thus far these methods all operate on a backbone level representation of proteins as a series of amino acid residues, and consequently, can only scaffold motifs represented at the backbone level.

Current approaches attempt to overcome this limitation for atom level active site descriptions by enumerating possible conformations and sequence indices for the catalytic residues and then in a separate step using motif scaffolding to generate proteins which scaffold these backbone positions [1, 3]. While active enzymes have been generated using this approach, it is computationally inefficient and does not scale to more complex active sites as the number of combinations of backbone coordinates to scaffold grows exponentially with the number of catalytic residues. A method capable of scaffolding complex theozymes described at the atom level would have wide-spread applications for enzyme design and beyond [16–18].

We reasoned that substantially improved performance on more complex enzyme active site scaffolding challenges could be achieved by a generative model capable of selecting the conformations and sequence indices of the catalytic residues by modeling the full joint distribution of rotamers, sequence indices, and scaffolds conditioned directly on atom level active site descriptions. With RFdiffusion2, we set out to extend RosettaFold diffusion All-Atom (RFdiffusionAA) [19] to generate structures conditioned on these minimal active site descriptions (Figure 1).

**Figure 1:**
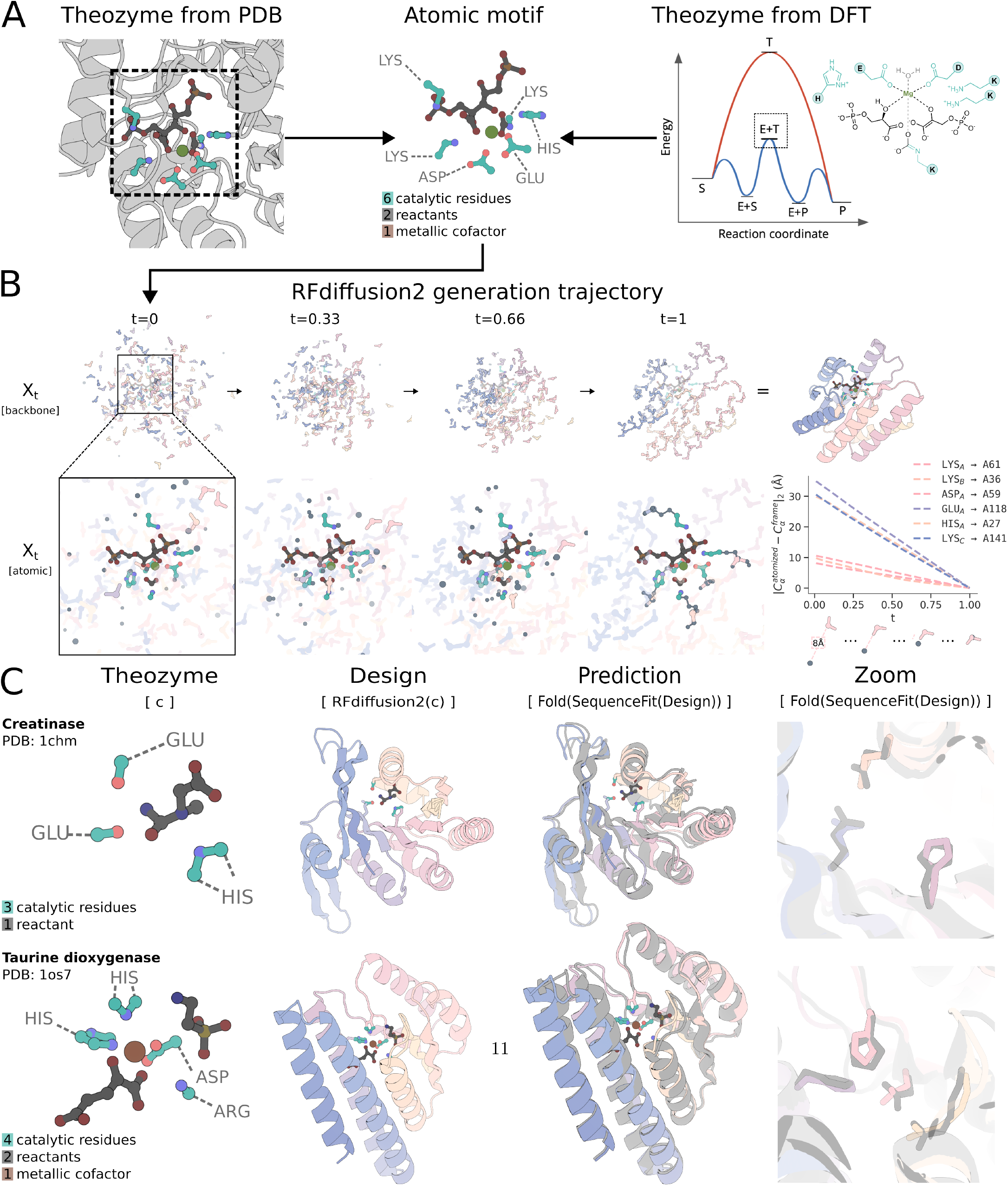
RFdiffusion2 overview.**A**.*De novo* enzyme design starts from a configuration of catalytic groups around the reaction transition state(s) (a theozyme) generated using quantum chemistry, protein structural analysis and/or chemical reasoning.**B**.RFdiffusion2 generates protein structures that support the theozyme. Row 1: The backbone trajectory shows the amino acid residue frames (pas-tel) as they transform from a sample drawn from the noise distribution into a protein backbone. Row 2: Zoom in of Row 1, showing the non-motif side chain atoms (slate grey) connecting the atomic motif (teal) with the protein backbone. At *t* = 1 the intra-residue bonds are shown for the atomized residues. (right) The distances between the C *α* coordinates of the unindexed, atomized residues and the backbone residues they superimpose at *t*= 1. Over the course of the trajectory, the model matches these unindexed residues to indexed residues of the protein backbone, such that by the end of the trajectory the unindexed residue’s C *α* occupies the same location as the C *α* of the protein backbone in Euclidean space.**C**.The design pipeline starts from the input theozyme, followed by RFdiffusion2 to generate the structure, and LigandMPNN to generate amino acid sequences that encode the structure and stabilize the transition state. Designs are evaluated by all atom structure prediction (using Chai-1, AF3, etc) and are considered successful if the design (pastel) and prediction (light gray) align to a sufficient degree. Two representative examples of consistency between design model and predicted structure at the level we take to constitute a success are shown in the right panels. The two cases pictured are the Creatinase and Taurine dioxygenase motifs from the AME benchmark described in section 3 (AME IDs: M0096_1chm, M0129_1os7).

## 2 Atomic motif conditioning

To address this challenge, we sought to generalize motif scaffolding beyond sequence indexed, backbone level motifs. Prior motif scaffolding methods represent motifs as “backbone frames”, in which each amino acid’s N, C *α*, C backbone atoms are parameterized as an element in SE(3). Each motif frame requires a pre-specified sequence index that indicates the frame’s location along the backbone chain. In contrast, the protein component of an atom level active site description includes only the side chain functional group atoms, not the backbone, of the participating amino acid residues; these residues can be anywhere along the sequence, i.e. of unknown index. While the indexed frame representation is sufficient for scaffolding large contiguous domains that can be accurately described at the residue level, it is insufficient to express the task of scaffolding disconnected groups of atoms belonging to residues of unknown indices. Our approach is to create an extended representation with differing levels of resolution and index information that is capable of expressing more complex motifs. In RFdiffusion2, we use the RosettaFold All-Atom neural network architecture in which each residue in the input and output can be represented as a frame or as heavy atom coordinates, i.e. an atomized residue [19]. During training, we represent some residues with frames and atomize others. For each atomized residue, the network learns to model the distribution of side chain poses. By providing known coordinates for some side chain atoms, which we term the atomic motif, the network learns to model the distribution of proteins conditioned on the inclusion of such atomic substructures. At inference time, we can then condition on the coordinates of individual protein atoms such as the side chain (or backbone) functional groups present in a theozyme. This ability to condition on individual atoms rather than entire residues allows us to forgo inverse rotamer sampling [7] used in previous approaches and instead allow the model to simultaneously infer an appropriate rotamer and scaffold.

We can further extend the representation to remove the need to know the sequence indices of a motif in order to scaffold it (Appendix B.1). During training, we select subsets of residues, duplicate them, and remove their index features to create *unindexed residues*. Without any auxiliary losses, the network learns that unindexed residues are always superimposed on indexed residues in unnoised structures. By providing coordinates for unindexed residues, which we term an *unindexed motif*, the network learns to model the distribution of proteins conditioned on the inclusion of a known substructure at unknown indices. At inference time, we then have the flexibility to condition on a motif consisting of residues with known or unknown indices because theozymes do not prescribe the sequence indices of their constituent residues. The ability to condition on motifs without specifying their indices enables us to forgo naive index sampling used in previous approaches and instead allow the model to simultaneously infer indices for the motif while scaffolding it (Figure 2B).

**Figure 2:**
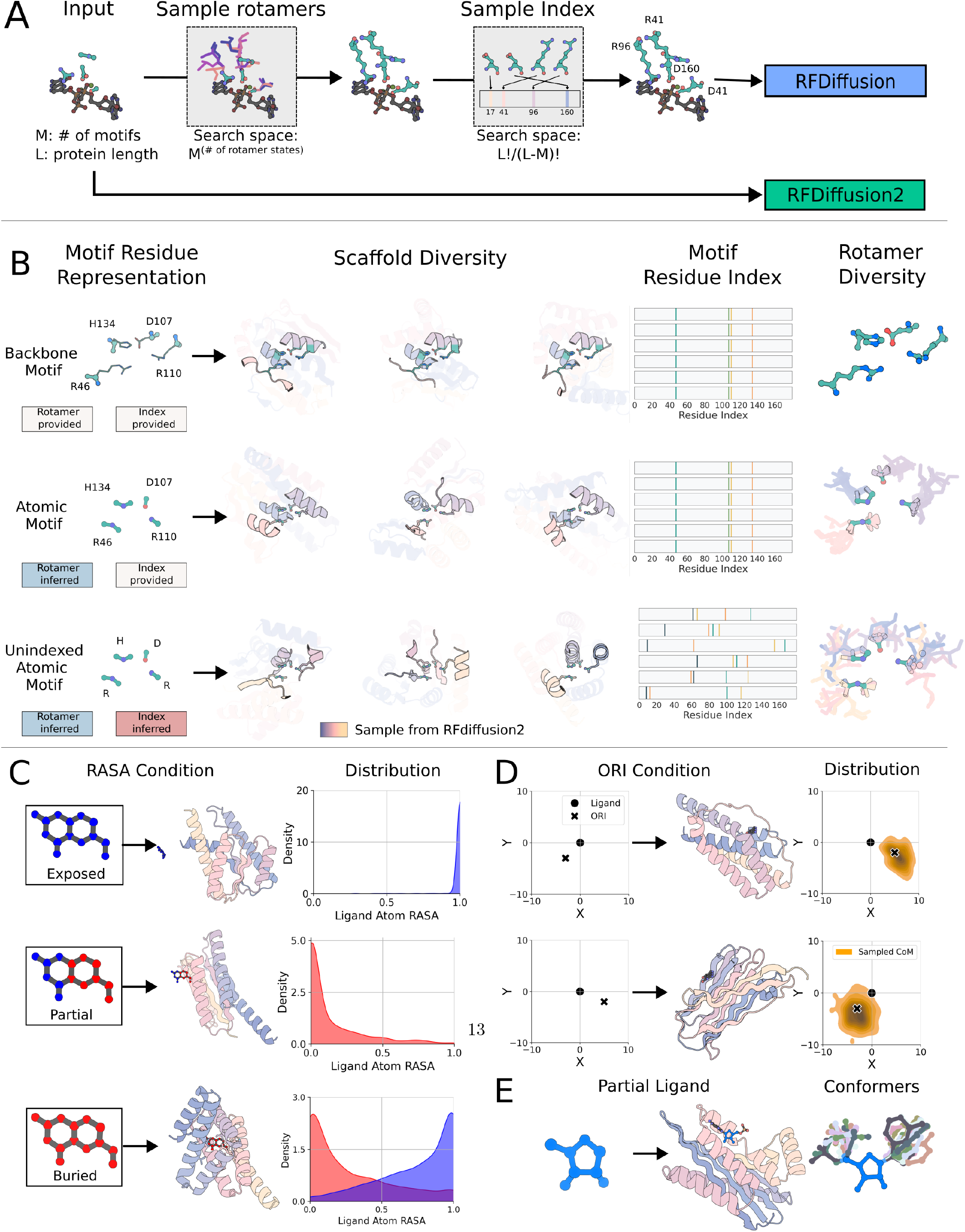
Motif scaffolding with RFdiffusion2. **A**. In the original RFdiffusion, two preprocessing steps are required to transform an unindexed atomic motif into a suitable input. These steps – inverse rotamer sampling and sequence index sampling – both require selecting from an exponentially large search spaces of *L*!*/*(*L* − *M*)! and *M*^(# of possible rotamer states)^ respectively, where *L* is the number of residues while *M* is the number of residues that are needed for the active site. RFdiffusion2 does not require such preprocessing steps and can scaffold unindexed atomic motifs directly. **B**. RFdiffusion2 can be conditioned on motifs in different representations. Three versions of the same motif (M0904, PDB 1qgx) from AME) are shown on the left most column, different backbone samples in the middle columns, and the resulting diversity of sequence indices and rotamers on the right most columns. (i) The backbone motif includes a pre-specified rotamer and index as required by RFdiffusion. (ii) The atomic motif has pre-specified sequence indices but unspecified side chain conformation. (iii) Only unindexed atom positions are provided, not the residue indices or side chain rotamer conformations. The rotamer and sequence indices are sampled during the RFdiffusion2 trajectories, increasing the diversity of possible solutions to the motif scaffolding problem. **C**. Each ligand atom can be labeled with a RASA category to control how solvent exposed the ligand is. The example RASA conditions are in the left column, a backbone sample with the ligand in the middle, and the distribution of ligand atom RASA from 100 designs with the RASA condition. When all atoms are labeled as exposed, the ligand RASA is concentrated around 1.0 and the backbone does not come into contact with the ligand. Conversely, when all atoms are labeled as buried, the ligand RASA is concentrated around 0.0; the sample shows the backbone almost completely covering the ligand. Labeling half the ligand as exposed and the other half as buried leads to RFdiffusion2 generating backbones that only bind to the buried side of the ligand. **D**. RFdiffusion2 can be provided with an `ORI` token that specifies the desired center of mass (CoM) of the scaffold with respect to the ligand. Two different `ORI` positions are shown in the left column. The middle column shows samples with scaffolds centered at the indicated `ORI` token positions. The distribution of CoMs from 100 sampled designs with the `ORI` token show that the scaffolds generally follow the `ORI` condition of where to place the scaffold. **E**. RFdiffusion2 can be provided with partial ligand input in which case it must sample the remaining ligand degrees of freedom while generating the protein. The left column shows the partial ligand input. The middle column shows in gray a conformer along with the protein generated by RFdiffusion2. Finally, the right column shows the distribution of 10 generated conformers. In fig. 7, we analyze the physical plausibility of the generated conformers.

The resulting model, RFdiffusion2, can generate proteins conditioned on a broad range of motifs including side chain motifs, motifs without known sequence indices, and ligand motifs. When training RFdiffusion and RFDiffusionAA, we found performance started to worsen over extended periods of training (Figure 5). Through a combination of auxiliary losses and self-conditioning [20], these methods were able to achieve high success rates on their respective tasks in short training sessions. Due to the complexity of the unindexed atomic motif scaffolding task, we expected that a stable objective would be required. To this end, we train RFdiffusion2 with flow matching [10, 11], a simpler framework for diffusion models [21, 22] shown empirically to have improved training and generation efficiency in other domains [23]. Briefly, this framework interpolates a training example towards a noise sample and trains a neural network to denoise by predicting the original, uncorrupted example (Appendix B.2). If trained to denoise with sufficient accuracy, the model can sample from the data distribution by iteratively denoising a sample drawn from the noise prior. Our representation of the data distribution contains both atoms and backbone frames which are elements of R^3^ and SE(3), respectively. Flow matching on R^3^ follows its original derivation using Gaussian probability paths while for SE(3) we follow the formulation in Frame-Flow [24, 25] that utilizes Riemannian flow matching [26] and removes approximations for rotational losses present in the RFdiffusion [12]. With these improvements, RFd-iffusion2 trained stably from randomly initialized neural network weights, does not require auxiliary losses, or use self-conditioning. Decoupling RFdiffusion2 from structure prediction is an important step to remove constraints around the neural network architecture as well as enable new generative modeling tasks.

Our training dataset consists of biomolecular structures from the Protein Data Bank (PDB) [27] that include proteins, protein–small molecule complexes, protein-metal complexes, and covalently modified proteins,filtering out common solvents and crystallization additives [19]. Each structure undergoes a motif extraction procedure to construct a motif-scaffolding training example (Appendix B.4). First, we select a random subset of residues to be the motif. Motif residues are chosen uniformly at random to ensure RFdiffusion2 sees a diverse range of interatomic geometries in the motif. Second, each motif residue is represented either as a frame or atomized residue. If a motif residue is atomized, only a random subgraph of the amino acid heavy atoms are provided as the motif. Third, we decide whether the motif will be featurized as an indexed or unindexed. Lastly, for any ligand present, we sample a random connected subgraph of its heavy atoms to be provided as a motif with Relative Accessible Surface Area (RASA) [28] labels for each atom in the ligand intermittently provided. We resample the motif on each training step as a form of data augmentation to ensure that the same noised structure is not always shown alongside the same motif. We train the final model for 17 days using 24 A100 Nvidia GPUs.

In our early experiments with flow matching, we found that the choice of how the ground truth structures were centered relative to the origin in training significantly affected the quality of inference outputs. A natural strategy is to globally center each ground truth structure; however, this leaks the offset between the scaffold center of mass and the motif. At inference time, this strategy requires the exact specification of the desired offset which is usually not known. A common fix for this pathology in motif-scaffolding diffusion models is to center ground truth structures on the motif, allowing the model to determine the offset between the scaffold center of mass and the motif [15, 25]. However even when the motif is centered, due to the interpolation scheme used in flow matching, the model is able to exactly determine this offset from any partially noised structure. The resultant behavior is that in the first denoising step, in which the network receives pure noise, it predicts an offset that it does not learn to refine in subsequent denoising steps. We instead introduce *stochastic centering* which first center the ground truth structure and add a small global translation sampled from a 3D Gaussian such that the noised input structures only encode an approximate offset between the motif and scaffold (Appendix B.3). This enables the model to refine the placement of the motif within the overall structure over the course of the inference trajectory. At inference time, users supply a prior belief about the placement of the motif through a special `ORI` pseudo-atom that specifies the approximate center of mass of the generated structure. This enables enzyme designers to control the active site and transition state orientation relative to the protein core (Figure 2D). For example, given an elongated small molecule or transition state with one end quite polar or charged, placement of the `ORI` token adjacent to the opposite end (or displaced from this end along a vector running through the long axis of the molecule) results in a designed binder or enzyme with a binding pocket extending radially from the center of the protein with the polar/charged end of the small molecule exposed to solvent.

RFdiffusion2 provides two additional conditioning capabilities of the ligand that are useful in *de novo* enzyme design. First, to provide finer control over the depth at which each reactant and/or cofactor is buried within the protein, we enable users to specify the RASA of each atom. By providing the RASA of each ligand atom 50% of the time during training, RFdiffusion2 learns to generate structures that respect those atomwise conditions at inference time when provided (Figure 2C). Second, the user may know the ligand atoms of a transition state but may not know the full ligand conformer. We allow the user to specify “partial ligands” where only the known ligand atoms are provided while RFdiffusion2 infers the rest of the ligand conformer. A brief analysis on the physical plausibility of the generated conformers shows they match closely with RDkit generated conformers (Figure 7). We find RFdiffusion2 can respect partial ligands by sampling physically realistic conformers (Figure 2D), removing the need to naively resolve the ligand conformer with an external tool *a priori*. Together, the conditioning capabilities opens up greater control over the geometric properties of the protein-ligand complex during inference (Appendix B.7).

## 3 Atomic Motif Enzyme Benchmark

An in silico enzyme motif scaffolding benchmark was introduced in RFdiffusion [12, 29] which contained one active site from each of the five most represented Enzyme Commission (EC) classes in the Mechanism and Catalytic Site Atlas (M-CSA) [30]. We find that this benchmark does not accurately reflect the challenges of *de novo* enzyme design due to its use of indexed backbone motifs, lack of ligands, and lack of active site diversity with only five enzyme cases. To evaluate RFdiffusion2, we developed a new benchmark which better reflects the theozyme scaffolding problem of *de novo* enzyme design (Appendix C).

We cross referenced the 958 hand-curated catalytic active sites downloaded from the M-CSA with the Proportion of Atoms Residing In Identical Topology (PARITY) [31] dataset and selected reactions with PDB crystal structures where all reactants and cofactors were present. After curation, we found 41 active sites spanning a diverse set of reactions in Enzyme Commission (EC) [29] classes 1-5 (Figure 3A; EC classes 1-5 account for 96% of examples in M-CSA). The annotations in M-CSA are at the residue level, so we extracted random connected sets of atoms from each catalytic residue to treat as the protein component of the theozyme for each benchmark case (Figure 3B). We evaluate RFdiffusion and RFdiffusion2 by sampling 100 structures for each benchmark case with the atomic motif as conditioning information. We assign 8 sequences to each structure with LigandMPNN [32], conditioned on the backbone, catalytic residue sidechain, and ligand coordinates. To evaluate whether the sequences fold to the intended structures, we use Chai-1 [33], an open-source implementation of AlphaFold3 [34], for structure prediction rather than AlphaFold2 [35] because we find that Chai-1’s side chain interaction distances more closely align with the reference distribution of native side chains (Figure 6). We call a design a success if (1) the Root Mean Square Deviation (RMSD) of all the heavy atoms in the catalytic residues is *<*1.5Å when aligned on the backbone N, C *α*, C of those catalytic residues in a Chai-1 prediction for at least one of the LigandMPNN sequences, and (2) the design contains no clashes with the ligand where a clash is defined as two atoms being within 1.5Å. We release this benchmark that we call Atomic Motif Enzyme (AME) for the scientific community.

**Figure 3:**
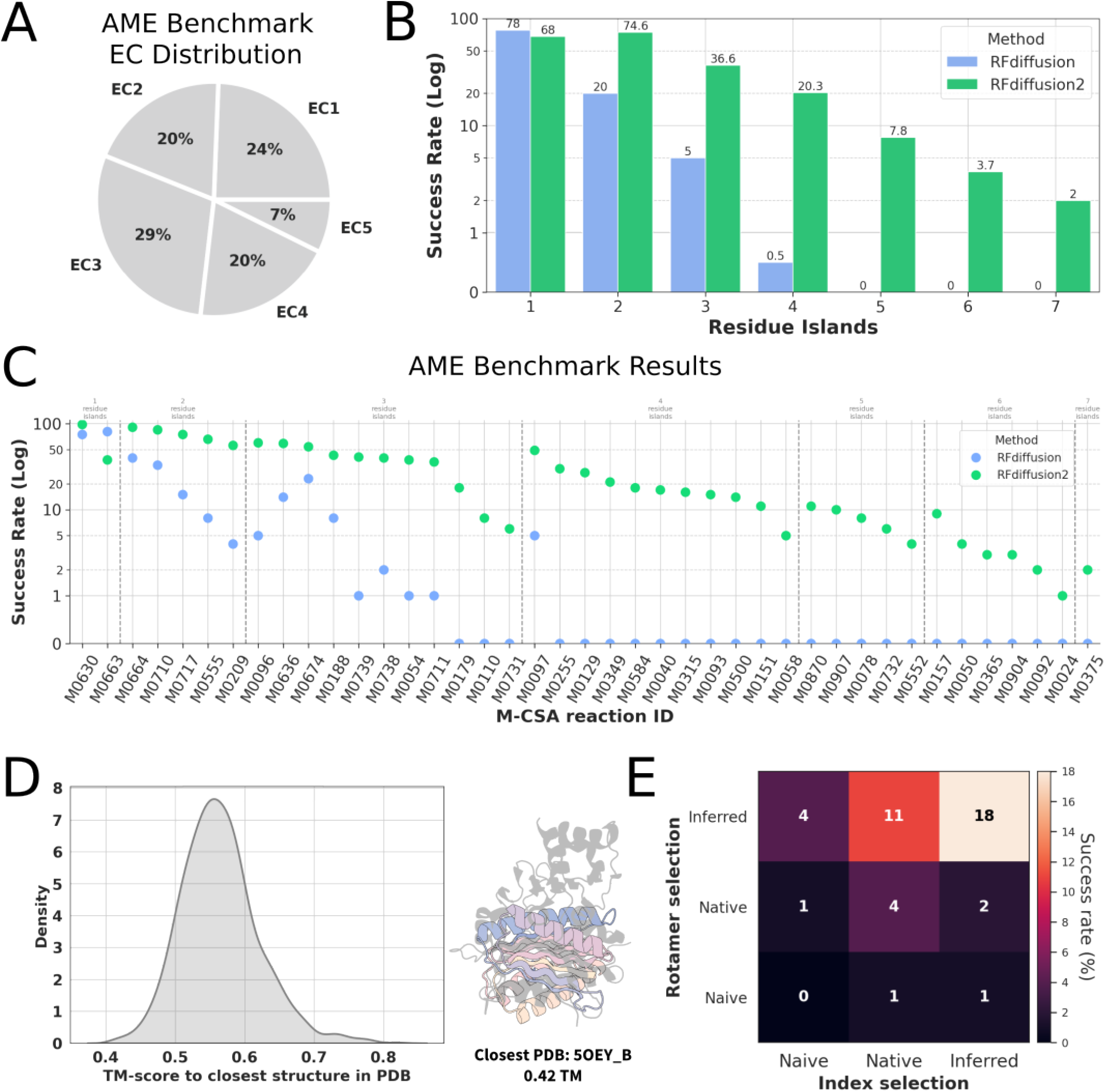
Atomic Motif Enzyme (AME) benchmark results.**A**.EC number distribution of the AME benchmark.**B**.Success rate of RFdiffusion2 and RFdiffusion as a function of the discontiguous chain segments containing motif atoms (the number of residue islands). While RFdiffusion performs marginally better on atomic motifs with one residue island, RFdiffusion2 is able to solve complex scaffolding problems with up to 7 residue islands.**C**.Success rate across each case for RFdiffusion2 and RFdiffusion. RFdiffusion2 samples at least one scaffold passing the successfilters for 41/41 cases while RFdiffusion achieves 16/41, and RFdiffusion2 has a higher success rate than RFdiffusion on all but one case. Difficulty correlates with more residue islands.**D**.Left: Distribution of TM-scores to the closest structure in the PDB for successful designs. Most designs have low closest TM-scores between 0.5 and 0.6. Right: a representative superposition of an RFdiffusion2 design (colored) on the closest PDB structure (in grey).**E**.Comparison of the three approaches for rotamer and index selection for representative AME benchmark cases from each of the residue island categories – 3, 4, 5, and 6. 100 designs were generated for each motif using each approach. Here we display the total success rate over all residue island categories while in Table 1 we show the success rate per category.*Inferred*: RFdiffusion2 generates the rotamer or index during the flow matching trajectory.*Native*: RFdiffusion2 is given the native rotamer or index at input.*Naive*: RFdiffusion2 as with RFdiffusion is provided with randomly selected side chain rotamers and residue indices. We find that “Naive” results in the worst overall success rate – as expected since RFdiffusion fails in most cases. Using native results in more successes, as expected, since the native rotamers and residue spacings are the result of evolutionary optimization. The best performance is achieved by allowing RFdiffusion2 to infer both rotamers and indices, which provides a far larger space to sample over to find optimal solutions than the other two approaches.

To our knowledge, RFdiffusion is the only deep learning method that has been shown to successfully design *de novo* enzymes [1, 2]. We compare RFdiffusion2 to RFdiffusion by establishing a pipeline that processes each atomic motif into a suitable input for RFdiffusion. The pipeline samples inverse rotamers and residue indices for the atomic motif to transform it into an indexed, backbone motif, and replaces the ligand with an attractive-repulsive potential. RFdiffusion2 generates enzymes conditioned directly on the theozyme without additional processing (Figure 2A).

We find that RFdiffusion2 finds solutions to all 41 benchmark cases while RFdiffusion finds solutions for only 16/41 cases. In 40/41 cases, we find that RFdiffusion2 significantly outperforms RFdiffusion, setting a new state-of-the-art for theozyme scaffolding (Figure 3C). We find the difficulty of a benchmark case correlates with the complexity of the motif which we quantify with the number of “residue islands”, the number of contiguous segments of catalytic residues in the original PDB structure. The successful designs from RFdiffusion2 are quite different from any protein in the training set as measured by FoldSeek [36] and TM-score [37](Figure 3D). Although the motif examples in AME come from the PDB, RFdiffusion2 is able to find completely novel scaffolds that house these motifs.

We next perform a study to understand the relative contribution of atomic motif and unindexed motif scaffolding to the improved success rates of RFdiffusion2. To resolve the rotamers of the atomic motif, we compare three approaches: *naive* inverse rotamer sampling as done with RFdiffusion, *inferring* the rotamer with RFdiffusion2, and the reference case of using the rotamer present in the *native* structure. For specifying the index of the atomic motif, we compare *naive* index sampling as done in RFdiffusion, RFdiffusion2 inference of the index, and the reference case of using the index present in the native structure. We evaluated every combination of the settings over four cases in the AME benchmark where each case is randomly chosen from 3, 4, 5, and 6 residue islands. We find the best strategy is to infer both the rotamer and sequence indices — even surpassing using the native rotamer and sequence indices, which would not be available when designing enzymes for novel reactions (Figure 3E). At 4 residue islands, we find the naive sampling strategy in RFdiffusion fails to achieve any successes (Table 1). We analyze the diversity of the residues surrounding the motif and find the greatest structural diversity with the unidexed atomic motif strategy followed by atomic motif and lastly backbone motifs with the lowest diversity (Figure 8). Our results show a deep learning approach to resolve the additional degrees-of-freedom represented by the rotamers and sequence indices is more effective than fixing these to specific values or naive enumeration.

## 4 *In vitro* results

Upon observation that RFdiffusion2 is capable of scaffolding theozymes according to our in silico success metrics, we experimentally tested whether the model was capable of generating functional enzymes from theozymes. For two reactions, we used theozymes from enzyme crystal structures in order to decouple the problem of theozyme design from theozyme scaffolding and directly assess the model’s capability on the latter. For another three reactions, we assessed whether it was possible to design a functional enzyme starting from only a desired catalytic mechanism, that is, without a priori knowledge of a functional theozyme geometry. For these cases, we perform optimization with Density Functional Theory (DFT) to find the saddle points of the energy landscape corresponding to transition state geometries of each case (Appendix D.3.1). In all four cases, we generate structures with RFdiffusion2 from the input theozyme,fit sequences to those structures, and filter them with structure prediction models to select designs for experimental validation (full details in appendix D). In all cases we find functional enzymes when testing less than 96 designed proteins. The specifics of theozyme preparation and experimental characterization for these four reactions are described below.

The aldol reaction forms carbon-carbon bonds between two carbonyl reactants. Enzymatic catalysis of this reaction allows for regio-and stereochemical control that would be impossible with non biological catalysts. Directed evolution campaigns have demonstrated that a catalytic tetrad composed of a nucleophilic lysine enmeshed in a hydrogen-bonding network with two tyrosines and an asparagine can stabilize the transition states of this reaction [38]. We construct a minimal theozyme, comprised of the hydrogen bond donors and acceptors of this network and the terminal CE and NZ atoms of lysine required to position the NZ for nucleophilic attack on the reactant, from the crystal structure of one such evolved retroaldolase: RA95.5-8F (PDB: 5AN7 [38]). We generated designs scaffolding this theozyme,filtered them, and expressed 96 in an in-vitro transcription/translation (IVTT) system. We tested activity of the IVTT produced proteins using racemic Methodol as a substrate and found four variants that show detectable levels of retro-aldolase activity in our semi-quantitative assay [39] (Figure 4A; Appendix D.1.2).

**Figure 4:**
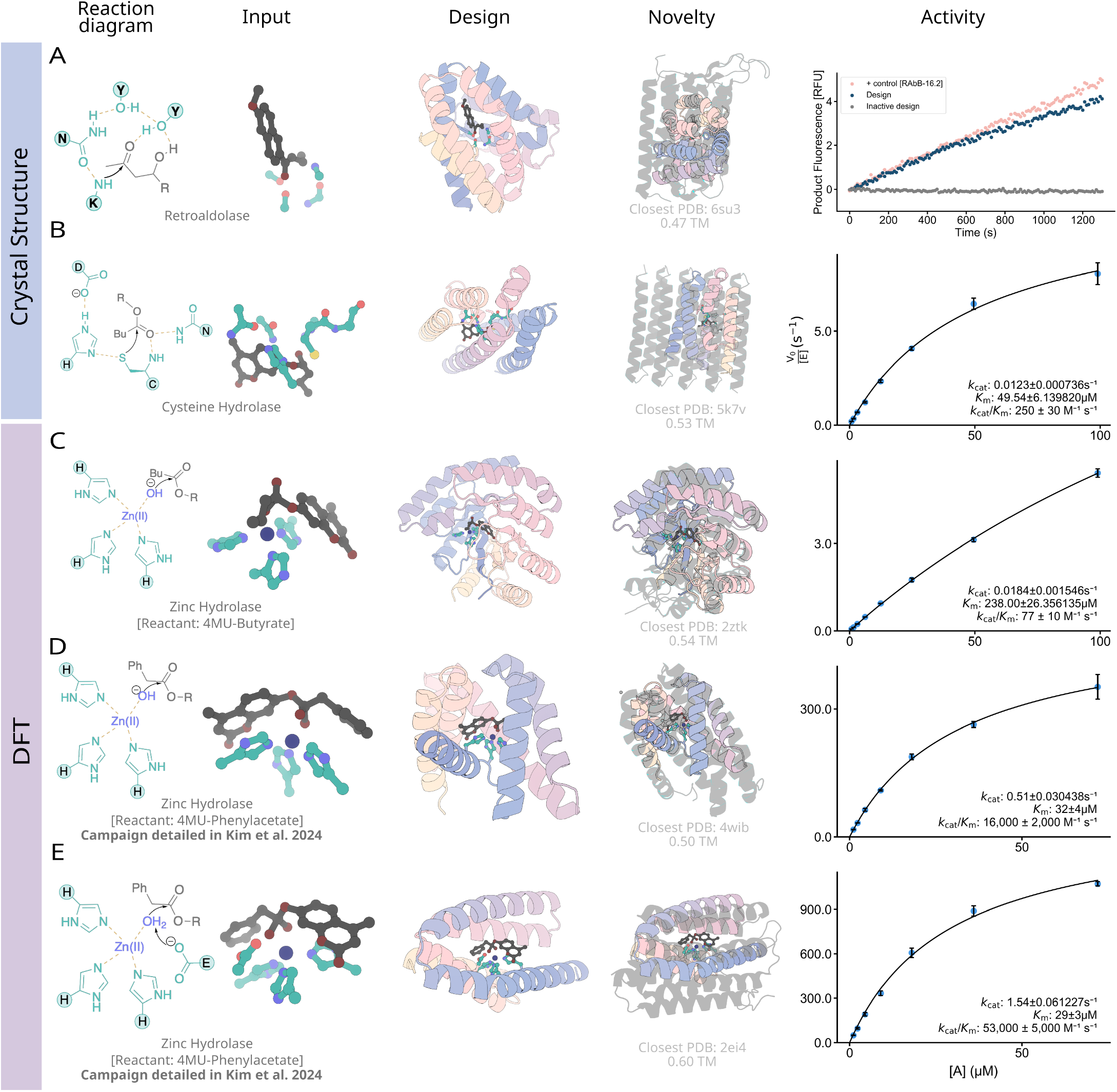
RFdiffusion2 generates active enzymes from minimal chemical constraints. Columns from left to right are reaction mechanism hypothesis, a theozyme based on this hypothesis input into RFdiffusion2, a resulting design, the closest protein in the training set (by Foldseek), and experimental activity.**A**.*Designed retroaldolase*. (From left to right) Retroaldolase reaction mechanism uses an activated lysine as a nucleophile; input to RFdiffusion2 (taken from PDB: 5an7); design from RFdiffusion2; superimposition with closest structure in the PDB (6su3; TM score=0.47), in vitro transcription translation assay measuring retroaldolase activity (pink is a enzyme optimized by directed evolution as a positive control, blue is design from RFdiffusion2 and grey is an inactive design).**B**.*Designed hydrolase using nucleophilic cysteine*. (From left to right) reaction mechanism involving a catalytic triad with a cysteine and an oxyanion hole to stabilize the transition state; inputs to the model derived from PDB: 1PPN; design from RFdiffusion2; superimposition with closest structure in the PDB (5k7v; TM score=0.53); kinetics assay measuring afluorescence of the product once hydrolyzed (*k*_cat_*/K*_*M*_ = 250*M*^−1^*s*^−1^).**C**.*Designed hydrolase using zinc as a Lewis acid as 4MU-Butyrate as a reactant*. (From left to right) reaction mechanism involving a activated water molecule as a nucleophile; DFT computed sidechain coordinates that position the zinc and substrate in the optimal geometry for the reaction; design from RFdiffusion2; superimposition with closest structure in the PDB (2ztk; TM score=0.54); kinetics assay performed in the same manner as in panel B (*k*_cat_*/K*_*M*_ = 77*M*^−1^*s*^−1^).**D**.*Designed hydrolase using zinc as a Lewis acid as 4MU-Phenylacetate as a reactant*. (From left to right) reaction mechanism involving a activated water molecule as a nucleophile; DFT computed sidechain coordinates that position the zinc and substrate in the optimal geometry for the reaction; design from RFdiffusion2; superimposition with closest structure in the PDB (4wiB; TM score=0.5); kinetics assay performed in the same manner as in panel B, C (*k*_cat_*/K*_*M*_ =16,000*M*^−1^*s*^−1^).**E**.*Designed hydrolase using zinc as a Lewis acid as 4MU-Phenylacetate as a reactant, and glutatmate as a general base*(From left to right) reaction mechanism involving a glutamate activating a water molecule which will serve as a nucleophile; DFT computed sidechain coordinates that position the zinc and substrate in the optimal geometry for the reaction; design from RFdiffusion2; superimposition with closest structure in the PDB (2ei4; TM score=0.60); kinetics assay performed in the same manner as in panel B, C (*k*_cat_*/K*_*M*_ = 53,000*M*^−1^*s*^−1^).

Ester hydrolysis is the cleavage of an ester bond by water. Enzymes which catalyze ester hydrolysis perform many cellular functions [40] and have numerous industrial applications [41, 42]. In cysteine hydrolases, catalysis of this reaction can be understood in terms of two components: a Cys-His-Asn catalytic triad in which the cysteine performs nucleophilic attack with the histidine acting as a general acid/base activated by the asparagine, and a helix-dipole-stabilized oxyanion hole created by the same cysteine’s backbone nitrogen together with a glutamine that stabilizes the negative charge of the tetrahedral intermediate formed during nucleophilic addition. We take as the minimal catalytic components for this reaction the active functional groups of the cysteine, histidine, asparagine, and glutamine, and 3-4 backbone atoms of the residues abutting the cysteine to force the local backbone into a region of Ramachandran space corresponding to the termination of a helix oriented in such a way that its dipole stabilizes the local oxyanion hole. We took the relative positions of these atoms from the crystal structure of a Papaya cysteine hydrolase (PDB: 1PPN [43]), and oriented a conformer of our reactant (4MU-Butyrate) according to the known optimal geometries for nucleophilic attack by the cysteine, proton donation by the histidine, and hydrogen bond stabilization of the oxyanion [44]. Among the 48 designs that were screened experimentally, several displayed detectable activity (Figure 16A); the best design shown exhibited multiple turnover activity with a *k*_cat_*/K*_*M*_ of 248 *±* 34*M*^−1^*s*^−1^ for the acylation step, exceeding previous results [45] for the same leaving group (Figure 4B).

Metallohydrolases coordinate metal ions and leverage their lewis acidity to activate water to form a potent nucleophile capable of hydrolyzing some of the most stable molecules in biological systems. Harnessing this mechanism for arbitrary substrates would enable the design of *de novo* enzymes capable of hydrolyzing environmental pollutants with long half-lives [46, 47]. In contrast to the previous case studies in which the theozyme geometry was extracted from native enzymes, here we perform both the theozyme design and theozyme scaffolding. To design the theozyme, we perform optimization in DFT to find the geometry of the transition state in which the hydroxide ion forms a bond with the carbonyl carbon, simulating the Zn(II) metal, metal-coordinating functional groups (imidazole), a chosen reactant, and a hydroxide ion. We performed this DFT optimization to create theozymes for two reactants – 4MU-Butyrate and 4MU-Phenylacetate – and generated enzymes with RFdiffusion2. We obtained 96 designs for each and identified 3 functional enzymes for the 4MU-B reaction and 5 for the 4MU-PA reaction. We found the best enzyme for the 4MUB reactant had a *k*_cat_*/K*_*M*_ of 77 ± 10 *M*^−1^*s*^−1^, higher than previously designed zinc hydrolases [48, 49]. The best enzyme for 4MU-PA, detailed in our companion paper [50], had a *k*_cat_*/K*_*M*_ of 16,000 *±* 2000*M*^−1^*s*^−1^, several orders of magnitude higher than previously designed zinc hydrolases. Typically in native zinc hydrolases, a general base (Glu/Asp/His) is present within hydrogen-bonding distance of the Zn(II)-bound water. In the aforementioned zinc-hydrolase designs we relied on the chance inclusion of such a general base during sequence fitting; however, knockout experiments revealed that mutating it to alanine in the most active design for 4MU-PA had little-to-no effect on catalytic efficiency. To rectify this, we included a general base (GLU) in the theozyme at the optimal geometry according to DFT calculations and generated another set of designs with RFdiffusion2. Of the 96 tested, we identified 11 functional enzymes, the best of which had a *k*_cat_*/K*_*M*_ of 53,000 *±* 5000*M*^−1^*s*^−1^ (Figure 4C-E).

The experimental validation of designs from RFdiffusion2 demonstrates its ability to generate functional enzymes when screening less than 96 designs when the optimal catalytic geometry is known, as demonstrated by scaffolding theozymes from native enzymes, as well as when starting from only the reaction mechanism in the cases where the theozyme was generated by DFT. The most active design for each reaction is structurally distinct from all structures in the PDB (Figure 4; Novelty column).

## 5 Discussion

RFdiffusion2 outperforms the prior state of the art methods on *in silico* benchmarks, removes expert intuition necessary with prior backbone motif scaffolding and scaffold library methods, and can design enzymes with *in vitro* catalytic activity. RFdiffusion2 enables direct scaffolding of ideal active sites described at the atom level without pre-specifying sequence indices or enumerating side chain rotamers. We show on our newly curated AME benchmark that RFdiffusion2 substantially improves on RFdiffusion over a range of atom level active site descriptions. Our design campaigns for retroaldolases, cysteine hydrolases and zinc hydrolases found active and novel enzymes for each reaction. Our *in silico* success in the AME benchmark suggests RFdiffusion2 should be applicable to designing enzymes across many more reactions at higher success rates than the prior state-of-the-art.

There are several avenues for improvement. Despite RFdiffusion2’s success in obtaining active enzymes across four reactions, the enzymes designed by RFdiffusion2 are not as active as native enzymes. Our theozymes might not be capturing all the necessary interactions for high activity and RFdiffusion2 might be able to sample higher activity enzymes by expanding our theozyme definition to include more interactions that are necessary for catalysis [1]. Automating the design of enzymes from theozymes opens up for the first time the possibility of large-scale testing of varied theozymes to broaden our understanding of enzymes and validate mechanistic hypotheses of much wider scope than those testable with catalytic residue knockout experiments or directed evolution. Alternative neural network architectures such as Diffusion Transformers [51] and modules from AlphaFold3 [34] for all-atom tasks could improve RFdiffusion2 which uses the neural network architecture of RFAA. Finally, we expect that co-designing the protein sequence [13] and side chains [52] outside the active site could lead to more favorable pocket interactions with the substrate and potentially enable sequence-based guidance based on experimental kinetics data.

RFdiffusion2 should be immediately useful to protein designers working on design problems requiring atomic resolution modeling such as small molecule binding and enzyme design. We expect that the introduction of RFdiffusion2 and the AME benchmark will open up new research efforts in the machine learning community exploring designing new modeling approaches for atomic resolution protein design. To this end, we are making the RFdiffusion2 inference and training code freely available to the research community.

## Acknowledgments

We thank L. Goldschmidt and K. VanWormer for maintaining the computational and wet lab resources at the Institute for Protein Design, S. Mathis for insightful conversations about conditional generative modeling, J. Watson for valuable discussions on objective stability in generative models, S. Pellock and A. Lauko for their insight on the challenges faced by enzyme designers. This work was funded by the National Science Foundation Graduate Research Fellowship Program, Grant No. DGE-2140004 (S.M.W), The Howard Hughes Medical Institute (D.B., I.K.), the Audacious Project at the Institute for Protein Design (A.L., N.H.), the Open Philanthropy Project Improving Protein Design Fund (I.K., S.J.P.), the ETH Zurich (D.H.), The Bill and Melinda Gates Foundation INV-010680 (D.K., W.A.), and the Schmidt Futures Foundation (A.L., S.J.P., S.S.), NSF-GRFP (J.Y.), NSF Expeditions grant (award 1918839: Collaborative Research: Understanding the World Through Code) (J.Y., R.B., and T.S.J.), Machine Learning for Pharmaceutical Discovery and Synthesis (MLPDS) consortium (J.Y., R.B., and T.S.J.), the Abdul Latif Jameel Clinic for Machine Learning in Health (J.Y., R.B., and T.S.J.), the DTRA Discovery of Medical Countermeasures Against New and Emerging (DOMANE) threats program (J.Y., R.B., and T.S.J.), the DARPA Accelerated Molecular Discovery program (J.Y., R.B., and T.S.J.) and the Sanofi Computational Antibody Design grant (J.Y., R.B., and T.S.J.).

## Author Contributions

W.A., D.T. and D.B. conceived the study. W.A., J.Y., D.T., S.S., and H.R.A.T contributed to development of RFdiffusion2. W.A. trained all versions of RFdiffusion2 described in the study. W.A., S.S., and R.K. performed *in silico* benchmarking. W.A., S.S., J.Y., and R.K. analyzed RFdiffusion2 results. S.S., S.M.W., D.K., I.K., Y.K., and B.C. experimentally characterized designs. W.A, J.Y., R.K., and D.B were responsible for the manuscript writing. S.S., S.M.W., D.K., and I.K. were responsible for writing *in vitro* details. D.B., R.K., T.S.J. and R.B. offered supervision throughout the project.

