## Supplemental Information for "Atom level enzyme active site scaffolding using RFdiffusion2"

### Appendices

**A Additional Figures** **26**

**B RFdiffusion2 details** **32**

### A Additional Figures

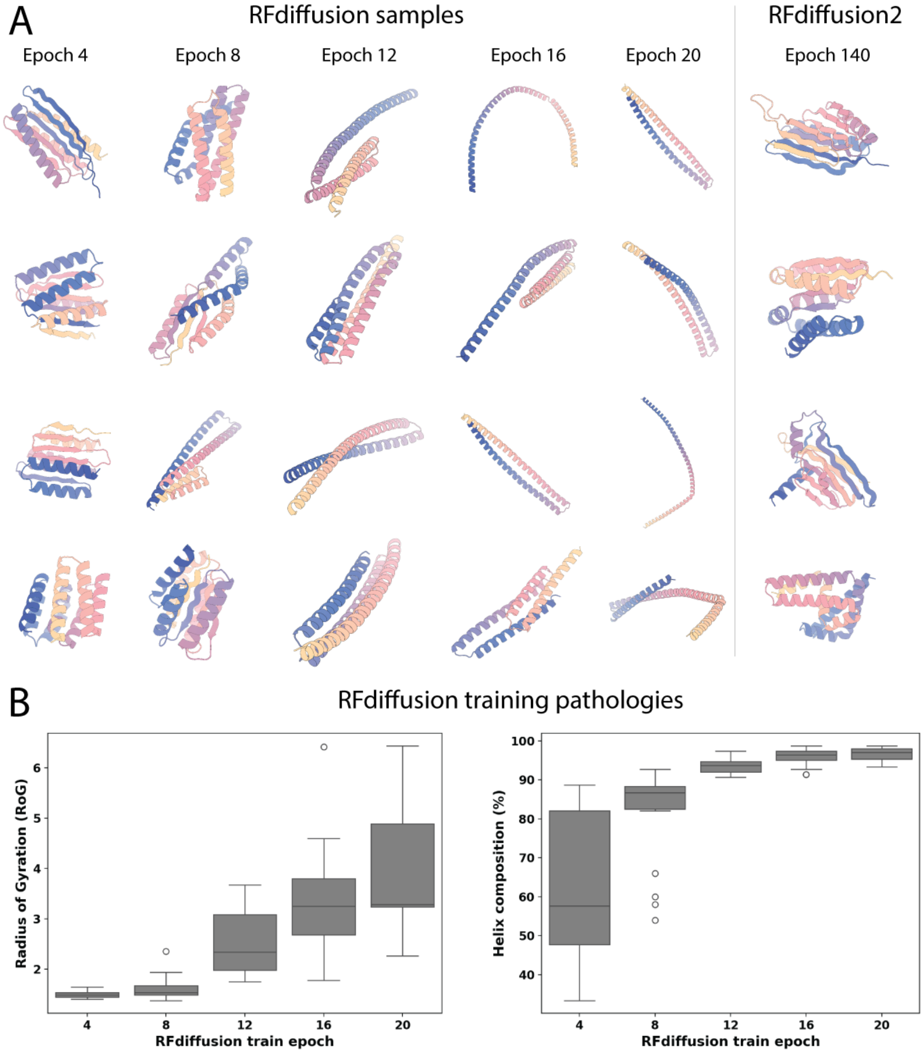

Figure 5 *(previous page)*: RFdiffusion2 does not experience RFdiffusion training pathologies. **A.** Each column shows four random samples after we trained RFdiffusion for 4, 8, 12, 16, and 20 epochs with the (unconditional) diffusion training objective in [12]. As a reference point, the published RFdiffusion model only trained for 5 epochs. We observe samples have diverse secondary structure of mixed beta sheets and alpha helices but become more helical as training progresses. Eventually starting in epoch 16 the samples start to only become long alpha helices. We find the training paradigm of RFdiffusion2 does not suffer from the same pathologies as RFdiffusion. Namely, we do not start from structure prediction weights and use only a principled flow matching objective. Shown are samples at epoch 140 where the samples retain secondary structure diversity with good quality. **B.** We take the five RFdiffusion checkpoint at each of the aforementioned epochs and sample 100 proteins of length 180 with different initial seeds. Using MDTraj [53], we quantify the Radius of Gyration (RoG) and alpha helix composition of the samples at each checkpoint. We find the RoG and helix composition steadily increase to a point where only helices generated. This matches the trend of the qualitative samples in panel A.

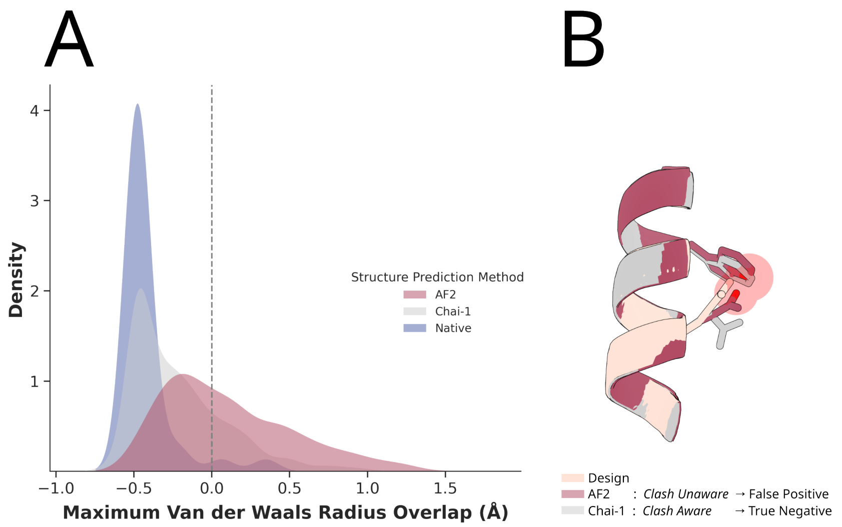

**Figure 6**: Comparison of physical plausibility of sidechain predictions across structure prediction methods. **A.** Kernel Density Estimates of the largest overlap of allowed nonbonded atoms' Van der Waal radii in structure predictions made by Chai-1 and AF2 (1000 structures) designed sequences and natives (36 structures). AF2 predictions frequently contain inter-atomic distances that correspond to nonphysical clashes: cases where the overlap is greater than zero, which are rarely observed in crystal structures. Chai-1 produces fewer clashes. **B.** An example of a false positive with our *in silico* RMSD filter if one uses AF2. Compared to our design (tan), AF2 predicts the side chains (red) that pass the motif atom RMSD filter but only due to packing the motif side chains with physical violations. This leads to false positives that would pass the motif heavy atom RMSD $< 1.5$Å filter but be non-physical. Chai-1, on the other hand, correctly rotates the motif side chains away to avoid a clash and rejects this design with a RMSD above the threshold. As a result, we use Chai-1 in our *in silico* metrics.

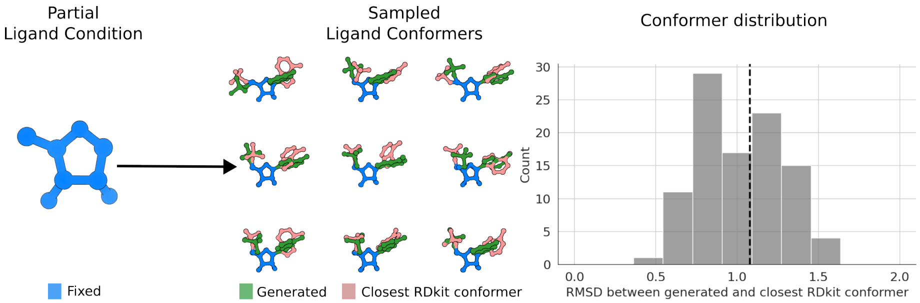

**Figure 7**: Partial ligand conformer analysis. We take case M0904 (PDB 1qgx) from the AME benchmark and generate 100 samples with a partial ligand condition taken from AMP. **Left**: shows the motif of AMP that RFdiffusion2 is given as the partial ligand. **Middle**: using RDkit, we generate 200 conformers of AMP with the partial ligand coordinates fixed. For each generated conformer in green, we find the conformer with the lowest RMSD and show it in orange. **Right**: the histogram shows the RMSDs between each generated conformer and its closest RDkit conformer. As a reference point, we draw a vertical dashed line of the RMSD between the native AMP conformer in the ground truth structure 1qgx. The RMSDs of the RFdiffusion2 conformers are centered on the native ligand RMSD with many conformers having $< 1.0$Å RMSD. This indicates RFdiffusion2 generates physical and plausible conformers that match conformers from RDkit.

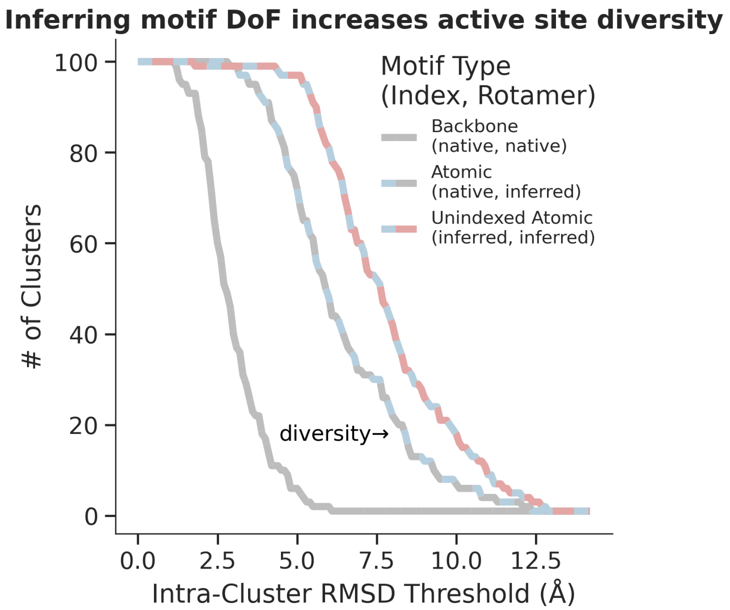

**Figure 8**: Comparison of the active site diversity across different motif-scaffolding modes. 100 designs were generated for a tetradic active site in AME (M0058_1cju) using three different representations of the motif as shown in Figure 2B: Backbone (using the full residue coordinates and sequence indices in the native enzyme [1cju]), Atomic (only providing the motif atoms but still providing the native sequence indices), and Unindexed atomic (only providing the motif atoms, no indices). For each of these three motif representations, the active site similarity is measured between all of the 4950 pairs of designs. The structural similarity local to the active site is assessed by aligning two designs on their motif (which superimpose perfectly), and then measuring the backbone N, C$\alpha$, C RMSD between the 7 residue fragments centered on each motif residue (the fragments pictured in Figure 2B Scaffold Diversity). These pairwise RMSDs are then used to agglomeratively cluster the active sites using the "complete linkage" criterion, under which an active site must have an RMSD below the intra-cluster threshold to all other active sites in the cluster. This clustering is performed for cluster thresholds ranging from 0 to 13.5Å with a step size of 0.1Å. The resulting plot of # of Clusters vs. Intra-Cluster RMSD Threshold will be right shifted for a design method that produces more diverse active sites (more clusters at the same clustering threshold). The diversity ordering indicated by this analysis, Unindexed Atomic > Atomic > Backbone, shows that allowing the model to resolve the DoF corresponding to Index/Rotamer selection results in more diverse active sites as measured by local backbone dissimilarity.

**Table 1**: Rotamer and index selection benchmarking. We display the data in Figure 3E stratified across each residue island category in table format. We see only using naive and native strategies for both index and rotamer selection achieve 0 success for 4 residue islands and beyond unless inferred is used for the rotamer. Using the inferred strategy for both index and rotamer achieves the best results overall.

| **Index** | **Rotamer** | **Residue Islands** | | | |
|---|---|---|---|---|---|
| | | 3 | 4 | 5 | 6 |
| Naive | Inferred | 12% | 0% | 4% | 0% |
| Naive | Native | 3% | 0% | 0% | 0% |
| Naive | Naive | 2% | 0% | 0% | 0% |
| Native | Inferred | 25% | 6% | 1% | 11% |
| Native | Native | 9% | 0% | 0% | 9% |
| Native | Naive | 5% | 0% | 0% | 0% |
| Inferred | Inferred | 45% | 8% | 14% | 6% |
| Inferred | Native | 7% | 0% | 1% | 0% |
| Inferred | Naive | 4% | 0% | 0% | 0% |

### B RFdiffusion2 details

#### B.1 Representations

***Amino acid representation.***

RFdiffusion2 introduces a novel biomolecule representation that can capture a wide range of inputs and outputs. Amino acid residues can be represented as follows.

- $X = [x_1, \ldots, x_N]$ is the collection of $N$ *indexed* amino acids residues $x_i$ with subscripts $i \in [1, \ldots, N]$ describing where each residue is along the polypeptide chain via a mapping `idx` (described later).
- $\bar{X} = [\bar{x}_1, \ldots, \bar{x}_{\bar{N}}]$ is the collection of $\bar{N}$ *unindexed* amino acid residues $\bar{x}_i$ with subscripts $i \in [1, \ldots, \bar{N}]$ that *do not* correspond to a location along the polypeptide chain. We use indices $i$ to enumerate through unindexed residues, but they have no connection to the polypeptide chain.

Each residue $x$ (or $\bar{x}$) can be represented as a frame $x \in \mathrm{SE}(3)$ or atomized $x \in \mathbb{R}^{14\times 3}$ following the amino acid 14 atom representation [35]. We acknowledge the abuse of notation since $x_i^{(t)}$ and $\bar{x}_{\bar{i}}^{(t)}$ have different dimensionalities and different spaces. We treat $X$ (and $\bar{X}$) as a "jagged" tensor[1] where each row of a dimension can have a different length to avoid unnecessarily verbosity, i.e. $\mathrm{Shape}(x_i^{(t)}) \neq \mathrm{Shape}(x_j^{(t)})$ for some $i \neq j$. This notation helps simplify the exposition. We will use the over head bar notation to refer to unindexed variables. For example, $\bar{X}$ is the collection of unindexed residues, $\bar{N}$ is the number of unindexed residues. Providing the sequence index as an input feature helps the model form a polypeptide chain by placing $x_i$ in between its neighboring residues in the chain. Unindexed residues are a way of prompting RFdiffusion2 to figure out the sequence index placement for $\bar{x}_i$.

Before we describe how unindexed residues work, we must describe how motifs are represented. There are two types of motif residues, indexed and unindexed, which we denote as

$$X^M = \{x_i^M : i \in \mathcal{M}\}, \qquad \bar{X}^M = \{\bar{x}_i^M : i \in \mathcal{M}\} \tag{1}$$

where $\mathcal{M} \subset [1, \ldots, N]$ is the collection of subscripts for indexed residues in the motif. Typically in motif-scaffolding with generative models, the model is provided $X^M$ and is tasked with generating the scaffold $X^S$ defined as the remaining residues in $X$ not provided for in $X^M$

$$X^S = \{x_i : i \in [1, \ldots, N]/\mathcal{M}\} \tag{2}$$

However, the scaffold cannot be defined for motif residues in $\bar{X}^M$ which do not have known sequence indices. This can be overcome by guessing the sequence indices for each unindexed motif residue, but the trial and error for guessing sequence indices in a combinatorial space of $[1, \ldots, N]$ is infeasible to enumerate. If a generated enzyme is bad, how can we know if it is due to the generative model or due to a poor choice of sequence indices?

[1] Pytorch introduced jagged tensors for handling elements of different lengths. See https://pytorch.org/FBGEMM/fbgemm_gpu-overview/jagged-tensor-ops/JaggedTensorOps.html

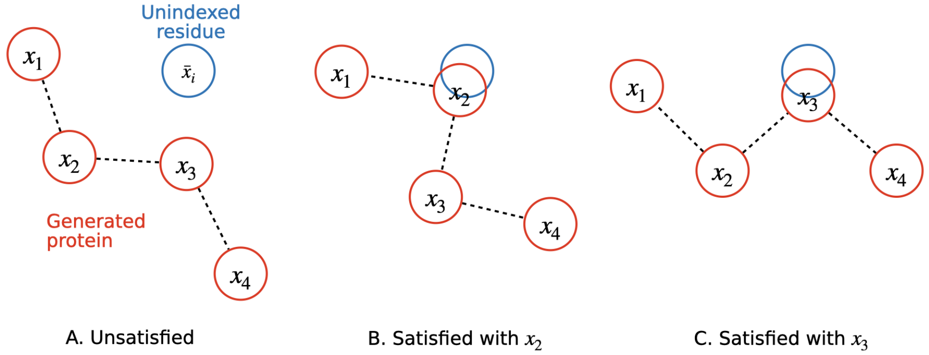

**Figure 9**: Diagram of unindexed residue satisfaction. In red are four indexed residues that make up the generated protein while in blue we show a single unindexed motif residue $\bar{x}$. **A.** The unindexed residue is not satisfied since none of the residues in the generated protein overlap with it. **B.** Here it is satisfied since $x_2$ overlap. The generated protein has placed this particular motif residue at index 2. **C.** Again it is satisfied but this time with $x_3$. Whichever residue in the generated protein used to satisfy $\bar{x}$ does not matter. It is up to the model to select which indexed residue to use.

***Unindexed residue satisfaction.***

Our representation allows for a solution where RFdiffusion2 can handle $\bar{X}$ without manually guessing sequence indices at the start. At a high level, we set up an input representation where the model generates a full protein $X$ but with the constraint that each unindexed residue in $\bar{X}$ has to have an overlapping indexed residue. Let us assume $X^M = \emptyset$ to simplify our notation and that $\bar{x}_i \in \mathrm{SE}(3)$ for each $i \in [1, \ldots, \bar{N}]$. A unindexed residue $\bar{x}_i$ is defined as being *satisfied* if the following holds

$$\left( \min_{j \in [1,\ldots,N]} \|\mathrm{Log}_{\bar{x}_i}(\bar{x}_i) - \mathrm{Log}_{x_j}(x_j)\|_g \right) < \epsilon \tag{3}$$

for some very small epsilon $\epsilon$ and where $\|\cdot\|_g$ is the norm with the Riemannian metric $g$ for SE(3) and $\mathrm{Log}_x$ is the logarithmic map onto the tangent space centered at point $x \in \mathrm{SE}(3)$. If $\bar{x}$ is satisfied then we can extract the sequence index that RFdiffusion2 assigned with the following

$$\arg\min_{j \in [1,\ldots,N]} \|\mathrm{Log}_{\bar{x}_i}(\bar{x}_i) - \mathrm{Log}_{x_j}(x_j)\|_g \tag{4}$$

An illustration of unidexed residue satisfaction is shown in Figure 9. RFdiffusion2's must satisfy each $\bar{x}_i$ with a indexed residue. In other words, it is protein generation while adhering to constraint satisfaction that we specify (or prompt) through our representation.

***Partially defined atomized motif residues.***

We have discussed $\bar{x}_i^M$ as if the residue coordinates are fully specified. However, it is often the case only the side chain functional groups are known for $\bar{x}_i^M$. Here, $\bar{x}_i^M$ is represented as an atomized residue, i.e. in $\bar{x}_i^M \in \mathbb{R}^{14\times 3}$. There are two tasks RFdiffusion2 must perform for partially specified unindexed atomized residues. First, it must satisfy the eq. (1) constraint in finding a indexed residue to satisfy $\bar{x}_i^M$. Second, it must generate the unspecified atoms in $\bar{x}_i^M$. We will denote $\bar{x}_i$ as the atom coordinates in $\bar{x}_i^M$ that are to be generated by RFdiffusion2. $\bar{x}$ takes the place of inverse rotamer sampling that was previously used when motif residue atom coordiantes were not fully specified. We now tasks the model with generating the atom coordinates that become the rotamers in the motif residue. Partially specified atomized residues are not unique to unindexed motif residues. Indeed, $x_i^M$ can also be specified as atomized residues with missing atom coordinates that must be inferred by the model.

***Ligand representation.***

Following RFdiffusionAA, we represent ligands, denoted $L \in \mathbb{R}^{K\times 3}$, as a collection of $K$ 3D atomic coordinates. Similar to partially defined atomized residues, ligands can be partially defined with the unknown atom coordinates left to RFdiffusion2 to infer. We use $L^M$ to denote the ligand motif that holds the unnoised ligand atoms provided to the model.

#### B.2 Generative modeling

RFdiffusion2 utilizes Flow Matching (FM) to generate proteins. We refer to Lipman et al. [54] for an introduction to (Riemannian) FM. Our exposition closely follows FrameFlow [25]. To be self-contained, we first discuss FM background (Appendix B.2.1) followed by the implemented forms of FM depending on the representation: frames or atomized (Appendix B.2.2).

##### B.2.1 Background

FM is a method for training Continuous Normalizing Flows (CNFs) which describe a process in which samples from a prior distribution $x^{(0)} \sim p^{(0)}$ are transformed into a sample from the data distribution $x^{(1)} \sim p^{(1)}$ via a *marginal flow*, $\phi^{(t)}$, defined in terms of a *marginal vector field* $v^{(t)}$ as

$$\frac{\mathrm{d}}{\mathrm{d}t}\phi^{(t)}(x) = v^{(t)}(\phi^{(t)}(x)) \quad \text{where} \quad \phi^{(0)}(x) = x \tag{5}$$

for some time $t \in [0, 1)$. We can derive the exact form of the flow or vector field with integration and differentiation of the Ordinary Differential Equation (ODE) in eq. (5) if the closed form of one is known. Unfortunately, neither the marginal vector field nor flow are available in closed form but the key idea of FM is to *learn* the marginal vector field by regressing the *conditional* vector field $u(x^{(t)}|x^{(1)}) = \frac{\mathrm{d}}{\mathrm{d}t}x^{(t)}$ from which we can integrate to derive the *conditional* flow $x^{(t)} = \phi^{(t)}(x^{(0)}|x^{(1)})$. The closed form of $x^{(t)}$ depends on the manifold which the data is generated on. In general, we can write the conditional flow $x^{(t)}$ as a geodesic path

$$x^{(t)} = \text{Exp}_{x^{(0)}}\left(t \cdot \text{Log}_{x^{(0)}}(x^{(1)})\right) \tag{6}$$

where $\text{Exp}_{x^{(0)}}$ and $\text{Log}_{x^{(0)}}$ are the exponential and logarithmic maps at point $x^{(0)}$. Then the conditional vector field can be written as

$$u(x^{(t)}|x^{(1)}) = \frac{\text{Log}_{x^{(t)}}(x^{(1)})}{1-t}. \tag{7}$$

The FM objective is then

$$\mathbb{E}_{\mathcal{U}(t;0,1),p^{(1)}(x^{(1)}),p^{(0)}(x^{(0)})}\left[\|u(x^{(t)}|x^{(1)}) - \hat{v}(x^{(t)})\|_g^2\right] \tag{8}$$

where $\hat{v}$ is the output of a neural network and $g$ is the Riemannian metric for the Manifold on which $x$ lies. Since we only deal with Euclidean and Lie groups, we can assume the norm takes the form of a L2. Using eq. (7), we can reparameterize the model output in terms of its *denoised* prediction $\hat{x}^{(1)}(x^{(t)})$. One can view this as the model being provided the noisy data $x^{(t)}$ for $t \in [0, 1)$ and predicting the *clean* data. The following objective will train the model to predict the marginal vector field

$$\mathbb{E}_{\mathcal{U}(t;0,1),p^{(1)}(x^{(1)}),p^{(0)}(x^{(0)})}\left[\|u(x^{(t)}|x^{(1)}) - \frac{\text{Log}_{x^{(t)}}(\hat{x}^{(1)}(x^{(t)}))}{1-t}\|_2^2\right]. \tag{9}$$

We will use eq. (9) as the form of the training objective henceforth. If the model is trained to low enough error across all $t$, then the learned vector field approximates the marginal vector field:

$$\underbrace{\hat{v}^{(t)}(x^{(t)}) = \frac{\text{Log}_{x^{(t)}}(\hat{x}^{(1)}(x^{(t)}))}{1-t}}_{\text{Learned}} \approx v^{(t)}. \tag{10}$$

Now with an approximation of the marginal vector field at hand, we can generate data by first drawing a sample from noise $x^{(0)} \sim p^{(0)}$ and simulating the ODE in eq. (5) using the Euler method. First, we select a number of timesteps, e.g. 100. Then set the step size to $\Delta = 1/100$. The following update are performed starting at $t = 0.0$ until $t = 1.0$

$$x^{(t+\Delta)} = \text{Exp}_{x^{(t)}}\left(\Delta \cdot \text{Log}_{x^{(t)}}\left(\hat{v}^{(t)}(x^{(t)})\right)\right) \tag{11}$$

The final step is a sample from the learned data distribution: $x^{(1)} \sim p^{(1)}$.

##### B.2.2 Diffusion with hybrid representations

We now turn to describing the generative model used in RFdiffusion2. Noised residues are denoted with $x_i^{(t)}$ and $\bar{x}_i^{(t)}$ for indexed and unindexed residues respectively. Noised ligand atoms are denoted as $l_i^{(t)}$. Collections of noised elements are denoted as

$$\text{(Noised indexed residues)} \qquad X^{(t)} = \left[x_1^{(t)}, x_2^{(t)}, \ldots, x_N^{(t)}\right], \tag{12}$$

$$\text{(Noised unindexed residues)} \qquad \bar{X}^{(t)} = \left[\bar{x}_{\bar{1}}^{(t)}, \bar{x}_{\bar{2}}^{(t)}, \ldots, \bar{x}_{\bar{N}}^{(t)}\right], \tag{13}$$

$$\text{(Noised ligand atoms)} \qquad L^{(t)} = \left[l_1^{(t)}, l_2^{(t)}, \ldots, l_K^{(t)}\right], \tag{14}$$

$$\text{(Noised biomolecule)} \qquad B^{(t)} = \left[X^{(t)} \middle| \bar{X}^{(t)} \middle| L^{(t)}\right], \tag{15}$$

$$= \left[b_1^{(t)}, b_2^{(t)}, \ldots, b_{N+\bar{N}+K}^{(t)}\right]. \tag{16}$$

For brevity, $B^{(t)}$ is used as shorthand for the noised residues and ligand atoms where $|$ is the concatenation operator leading to a length $N+\bar{N}+K$ tensor where $b_i^{(t)}$ is the $i$th element of $B^{(t)}$. When $t=0$, $B^{(0)}$ refers to the biomolecule with pure noise for each of its elements $b_i^{(0)}$ except for the motif elements. Similarly, $B^{(1)}$ refers to pure data for each of its elements $b_i^{(1)}$. As mentioned in appendix B.1, we use an abuse of notation to concatenate multiple tensors along their leading dimension despite the other dimensions and spaces being different.

RFdiffusion2 is provided with the motif information $X^M$, $\bar{X}^M$, and $L^M$ each time it predicts a vector field. We refer to

$$\text{(Motif biomolecule)} \qquad B^M = [X^M \mid \bar{X}^M \mid L^M] \tag{17}$$

as shorthand for the motif biomolecule that holds all the motif components.

There are other features RFdiffusion2 is provided with such as the bond geometry, atom types, and amino acid types of the ligand and residues. Each atom in the ligand can be paired with a RASA condition to have further control over how the generated protein is position relative to the ligand. We support three possible RASA conditions: exposed, buried, and partially exposed. Exposed corresponds to a RASA $> 0.1$; buried corresponds to RASA $< 0.1$; partially exposed corresponds to RASA between 0.1 and 1.0.

For succinctness, we use $C$ as the *condition* variable to hold the motif and auxiliary features used in the model architecture, i.e. bond geometry, atom types, amino amino acid types. Since the set of inputs is too large to enumerate, we use ellipses to indicate all inputs – each of which are described in appendix B.5.

$$\text{(Conditions and features)} \qquad C = \left[B^M, \ldots\right]. \tag{18}$$

**Motif convention.** Our notation assumes the motif $B^M$ is not part of $B^{(t)}$ but is part of $C$ (Equation (18)). This greatly simplifies how we discuss and enumerate through elements of $B^{(t)}$. Since the model inputs are always $(B^{(t)}, C)$, it is trivial for the model to learn to replace the motif residues and atoms in its predictions with the motif positions in $C$.

It is common to assume independence across each residue or ligand atom that is noised (as done in other domains such as images and video for pixels) such that it suffices to consider FM for each possible representation.

***Frame case***

$b_i^{(t)} = (\tau_i^{(t)}, r_i^{(t)}) \in \mathrm{SE}(3)$ where $\tau_i \in \mathbb{R}^3$, $r_i \in \mathrm{SO}(3)$. The conditional flows become

$$\tau_i^{(t)} = (1-t)\tau_i^{(0)} + t\tau_i^{(1)}, \qquad r_i^{(t)} = \mathrm{Exp}_{r_i^{(0)}}\left(t\mathrm{Log}_{r_i^{(0)}}(r_i^{(1)})\right). \tag{19}$$

As priors, we use $\tau_i^{(0)} \in \mathcal{N}(\vec{0}, \mathrm{I}_3)$ where $\mathrm{I}_3$ is the $3 \times 3$ identity matrix, $\vec{0}$ is the 3D vector of 0s, and $\mathcal{N}(\mu, \Sigma)$ is the normal distribution with mean $\mu$, covariance $\Sigma$; $r_i^{(0)} \in \mathcal{U}(\mathrm{SO}(3))$ where $\mathcal{U}(\mathrm{SO}(3))$ is the uniform distribution over SO(3). The end points $r_i^{(1)}$ and $\tau_i^{(1)}$ are the true residue frames from the data distribution. The conditional vector fields, eq. (7), and the model's predicted marginal vector fields are then

$$\text{(Conditional vector fields)} \qquad u^{\mathbb{R}}(\tau_i^{(t)}|\tau_i^{(1)}) = \frac{\tau_i^{(1)} - \tau_i^{(t)}}{1-t} \tag{20}$$

$$u^{\mathrm{SO}(3)}(r_i^{(t)}|r_i^{(1)}) = \frac{\log_{r_i^{(t)}}\left(r_i^{(1)}\right)}{1-t} \tag{21}$$

$$\text{(Predicted vector fields)} \qquad \hat{v}_\theta^{\mathbb{R}}(\tau_i^{(t)}; B^{(t)}, C, \ell) = \frac{\hat{\tau}_i^{(1)}\left(B^{(t)}, C, \ell\right) - \tau_i^{(t)}}{1 - \min(t, \epsilon)} \tag{22}$$

$$\hat{v}_\theta^{\mathrm{SO}(3)}(r_i^{(t)}; B^{(t)}, C, \ell) = \frac{\log_{r_i^{(t)}}\left(\hat{r}_i^{(1)}\left(B^{(t)}, C, \ell\right)\right)}{1 - \min(t, \epsilon)}. \tag{23}$$

We use $\theta$ to denote the neural network weights. We follow RFdiffusionAA in applying the structure loss across the intermediate and final layers of the SE(3)-transformer during training. $\hat{\tau}_i^{(1)}\left(B^{(t)}, C, \ell\right)$ and $\hat{r}_i^{(1)}\left(B^{(t)}, C, \ell\right)$ are the denoised rotation and translation outputs of the $\ell$th layer in the SE(3) transformer of the RFdiffusion2 architecture. At inference time, we will use the output of the final layer. Following Yim et al. [24], we truncate the vector fields by $\epsilon$ during training to avoid a unstable loss. During training, we set $\epsilon = 0.9$ and during sampling we set $\epsilon = 1.0$. Equation (22) and eq. (23) show that the model predictions, $\hat{\tau}_i^{(1)}$ and $\hat{r}_i^{(1)}$, take the noised biomolecule $B^{(t)}$ and condition $C$ as input to make its predictions. The training objective take the following form. For brevity, we omit $\theta$ in the loss arguments.

$$\mathcal{L}_{\mathrm{SE}(3)}\left(x_i^{(t)}, x_i^{(1)}, B^{(t)}, C, \ell\right) = \mathcal{L}_{\mathrm{SO}(3)}\left(r_i^{(t)}, r_i^{(1)}, B^{(t)}, C, \ell\right) + \mathcal{L}_{\mathbb{R}}\left(\tau_i^{(t)}, \tau_i^{(1)}, B^{(t)}, C, \ell\right) \tag{24}$$

$$\mathcal{L}_{\mathbb{R}}\left(\tau_i^{(t)}, \tau_i^{(1)}, B^{(t)}, C, \ell\right) = \|u^{\mathbb{R}}(\tau_i^{(t)}|\tau_i^{(1)}) - \hat{v}_\theta^{\mathbb{R}}(\tau_i^{(t)}; B^{(t)}, C, \ell)\|_2^2 \tag{25}$$

$$\mathcal{L}_{\mathrm{SO}(3)}\left(r_i^{(t)}, r_i^{(1)}, B^{(t)}, C, \ell\right) = \|u^{\mathrm{SO}(3)}(\tau_i^{(t)}|\tau_i^{(1)}) - \hat{v}_\theta^{\mathrm{SO}(3)}(\tau_i^{(t)}; B^{(t)}, C, \ell)\|_2^2 \tag{26}$$

***Euclidean case***

$b_i^{(t)} \in \mathbb{R}^{14\times 3}$ or $b_i^{(t)} \in \mathbb{R}^3$. The form is identical to the translation case above: eq. (20), eq. (22), and eq. (25). In the case of atomized residues, the leading dimension is 14 coordinates in 3D instead one 3D coordinate for the translation. For instance, instead of $\tau_i^{(0)} \in \mathcal{N}(\vec{0}, \mathrm{I}_3)$, we have $\tau_i^{(0)} \in \mathcal{N}(\vec{0}, \mathrm{I}_3)^{14}$.

***Unindexed case***

$b_i^{(t)} = \bar{x}_i^{(t)} \in \bar{X}^{(t)}$. As discussed in appendix B.1, we require a training objective that teaches RFdiffusion2 to satisfy $\bar{x}_i^{(t)}$ by placing it over a indexed residue. We can construct this by randomly pairing each unindexed residue with a indexed residue. Assuming $\bar{x}_i^{(1)}$ is the ground truth for unindexed residue $\bar{x}_i^{(t)}$, we set $\bar{x}_i^{(1)} = x_j^{(1)}$ for some $x_j^{(1)} \in X^{(1)}$ chosen uniformly at random. We sampling a new set of index pairings $(i, j)$ for each $i \in [1, \ldots, \bar{N}]$ at the beginning of each training step. Though we set $\bar{x}_i^{(1)} = x_j^{(1)}$, any sequence index information about $\bar{x}_i^{(1)}$ is not provided to the model. The model must figure out at each time $t$ the true indexed residue that $\bar{x}_i^{(t)}$ was computed from. As $t$ gets closer to 1, the flow trajectory should reveal to which indexed residue The training teaches RFdiffusion2 to place unindexed residues on indexed residues such that at inference time, when generating a protein, all unindexed residues of the motif will be part of the generated protein.

##### B.2.3 Hybrid representation loss

On each training step, we take a biomolecule $B^{(1)}$ from the training dataset then sample a random set of conditions $C$. Using the notation from eq. (16), we present the training loss in terms of each element $b_i^{(t)}$.

$$\mathcal{L}(B^{(1)}, C) = \mathbb{E}_{\substack{\mathcal{U}(t;0,1)\\ p^{(0)}(B^{(0)})}} \left[ \sum_{\ell=1}^{N_{\text{layers}}} \frac{\gamma^{(N_{\text{layers}}-\ell)}}{\sum_{\ell=1}^{N_{\text{layers}}} \gamma^{(N_{\text{layers}}-\ell)}} \sum_{i=1}^{N+\bar{N}+K} \mathcal{L}_{\text{FM}} \left( b_i^{(t)}, b_i^{(1)}, B^{(t)}, C, \ell \right) \right] \tag{27}$$

$$\mathcal{L}_{\text{FM}} \left( b_i^{(t)}, b_i^{(1)}, B^{(t)}, C, \ell \right) = \begin{cases} \mathcal{L}_{\text{SE}(3)} \left( b_i^{(t)}, b_i^{(1)}, B^{(t)}, C, \ell \right) & \text{if } b_i^{(1)} \in \text{SE}(3) \\ 10 \cdot \mathcal{L}_{\mathbb{R}} \left( b_i^{(t)}, b_i^{(1)}, B^{(t)}, C, \ell \right) & \text{if } b_i^{(1)} \in \mathbb{R}^{14\times 3} \text{ or } \mathbb{R}^3 \end{cases}. \tag{28}$$

The loss is stratified based on the representation of $b_i^{(1)}$. In eq. (28), we upweight the loss over the atomized residues and ligand coordinates by 10 compared to the loss over the frames. This is to place emphasis on correctness of bond geometries. The loss in eq. (27) includes a summation over all the layers in the SE(3)-transformer of the architecture. Following RFdiffusionAA, we apply a weight $\gamma^{(N_{\text{layers}}-\ell)}$ for $0 < \gamma < 1.0$ that weights the loss less in the early layers and more in the deeper layers. In this work, we set $\gamma = 0.95$. Lastly, we note that all coordinates in 3D are scaled by 0.25 during training and inference as done in RFdiffusionAA. We scale all coordinates by

4.0 when outputting the final samples. Compared to RFdiffusion, we emphasize that RFdiffusion2 does not use any additional auxiliary losses.

##### B.2.4 Hybrid noising

Following the notation and conventions in appendix B.2.2, we describe the noising process for a biomolecule $B^{(1)}$ with a hybrid representation. The noising process is a function of the ground truth biomolecule $B^{(1)}$ and the noise time $t$. The pseudocode is provided in algorithm 1.

---

**Algorithm 1** NoiseBiomolecule

---

**Require:**

$B^{(1)}$ Ground truth biomolecule

$t$ Noise time

$B^{(t)} \leftarrow []$
**for** $b^{(1)}$ in $B^{(1)}$ **do**
  **if** $b^{(1)}$ is frame **then**
    $(r^{(1)}, \tau^{(1)}) \leftarrow b_i^{(1)}$
    $r^{(0)} \sim \mathcal{U}\,(\mathrm{SO}(3))$
    $\tau^{(0)} \sim \mathcal{N}(\vec{0}, \mathrm{I}_3)$
    $r^{(t)} \leftarrow \mathrm{Exp}_{r^{(1)}}\left(t \cdot \mathrm{Log}_{r^{(1)}}(r^{(0)})\right)$
    $\tau^{(t)} \leftarrow t \cdot \tau^{(1)} + (1 - t) \cdot \tau^{(0)}$
    $b^{(t)} \leftarrow (r^{(t)}, \tau^{(t)})$
  **else if** $b^{(1)}$ atomized or ligand atoms **then**
    $b^{(0)} \sim \mathcal{N}(b_i^{(1)}, \mathrm{I}_3)^{14}$
    $b^{(t)} \leftarrow t \cdot b_i^{(1)} + (1 - t) \cdot b^{(0)}$
  **end if**
  $B^{(t)} \leftarrow \mathrm{Concat}(B^{(t)}, b^{(t)})$
**end for**
**Return** $B^{(t)}$

---

#### B.3 Centering

Careful centering of the inputs is necessary to avoid data leakage. Since RFdiffusion2 primarily uses distances as features, it can learn to memorize the offsets between a motif and the noised structure. In this section, we provide further intuition and the issues that arise with common strategies. We demonstrate a novel centering strategy in RFdiffusion2 that we found greatly improves learning and sampling.

Assume we are in the training phase and have a protein $X$ with a motif $X^M$. For the purposes of explaining centering, we can assume each residue is atomized, no unindexed residues $\bar{X} = \emptyset$, and no ligand $L = \emptyset$. Our findings still hold if unindexed residues and ligands are included in the motif. The Center-of-Mass (CoM) is computed with the following operation

$$\mu(X) = \frac{1}{(\#\text{ atoms in } X)} \sum_{i=1}^{(\#\text{ atoms in } X)} \text{reshape}(X, [-1, 3])_i. \tag{29}$$

The reshape operation in eq. (29) is equivalent to the reshape operation in pytorch. Here we utilize reshape to turn the input of $\mu(\cdot)$ into a shape $[-1, 3]$ tensor where the first dimension is the number of atoms and the second dimension holds the 3D coordinates of each atom. The notation $\text{reshape}(\cdot, [-1, 3])_i$ is indexing into the $i$th atom of the reshaped tensor.

During training, we noise the protein (specifically the scaffold residues) $X^{(t)}$ up to the mean and variance prescribed by $t$ according to the linear interpolation schedule (Equation (19)). The endpoints $X^{(1)}$ and $X^{(0)}$ correspond to the ground truth structure and the pure noise structure sampled from a isotrpoic gaussian, respectively. By linearity, the CoM at any $t \in [0, 1]$ can be computed as

$$\mu\left(X^{(t)}\right) = t \cdot \mu\left(X^{(1)}\right) + (1 - t) \cdot \mu\left(X^{(0)}\right) = t \cdot \mu\left(X^{(1)}\right) \tag{30}$$

where the second equality follows because $\mu\left(X^{(0)}\right) = \vec{0}$. We use $\vec{0}$ to denote a vector of zeros, i.e. $\vec{0} = [0, 0, 0]$. Equation (30) will become important to show the systematic issues with different centering strategies. Deep neural networks are powerful in that they can learn to exploit unintended patterns or artifacts in the data that do not lead to better generalization. The neural network of RFdiffusion2 uses distances and orientations as the primary features to be SE(3) equivariant and has knowledge of $t$ as an input. Therefore, it is possible for RFdiffusion2 to back-calculate various SE(3) equivariant quantities such as distances and orientations.

We start by describing the issues with global and motif centering. Then, we present our solution of stochastic centering.

##### *Global centering*

One approach is to globally center the protein such that $\mu\left(X^{(1)}\right) = \vec{0}$. As a result, $\mu\left(X^{(t)}\right) = \vec{0}\ \forall t \in [0, 1]$. The distance and orientation of $X^M$ and $X^{(t)}$ are provided as features which allows RFdiffusion2 to become aware of the offset between the motif and the noised protein

$$\mu\left(X^M\right) - \mu\left(X^{(t)}\right) = \mu\left(X^M\right) - \vec{0}. \tag{31}$$

The issue is the model then becomes sensitive to the placement of the motif relative to the noised protein. One has to select the placement of the motif carefully at sample time relative to the origin. The model learns to always keep the protein centered and never change the CoM towards or away from the motif. We find that placing the motif too far or too close will result in poor sample quality and diversity.

***Motif centering***

Here we consider the strategy of centering based on the motif such that $\mu\left(X^M\right) = 0$. While this strategy appears to work given the success of prior works [12, 25], we find a SE(3) equivariant model such as RFdiffusion2 can extrapolate the ground truth's CoM using the following formula: starting from eq. (30)

$$\mu\left(X^{(1)}\right) = \frac{\mu\left(X^{(t)}\right)}{t} \tag{32}$$

$$= \mu\left(X^{(t)}\right) - \mu\left(X^{(t)}\right) + \frac{\mu\left(X^{(t)}\right)}{t} + \underbrace{\frac{1-t}{t}\mu\left(X^M\right)}_{=0} \tag{33}$$

$$= \mu\left(X^{(t)}\right) + \frac{1-t}{t}\left(\mu\left(X^{(t)}\right) - \mu\left(X^M\right)\right). \tag{34}$$

One can easily show Equation (34) is an $SE(3)$ equivariant function and so it can be learned by the network since $t, X^M, X^{(t)}$ are provided as features. The network can learn to compute the exact center of mass of the ground truth (left hand side) from its inputs at any $t > 0$ (right hand side). We observed motif centering performing poorly when we saw at inference time that adding noise partway through the generation trajectory often resulted in poor samples that would extrapolate the sample's CoM at a very far distance from the motif. We saw the model was exploiting knowledge of $t$ and the displacement with respect to the motif which was always fixed at the origin. In light of this, we develop a new centering technique called stochastic centering to prevent any reliance on time based displacements.

***Our solution: Stochastic centering***

We will first present how we center in algorithm 2 then discuss its benefits.

**Algorithm 2** StochasticCentering

**Require:** Biomolecule $B$

$A \leftarrow []$
**for** $b$ in $B$ **do**
    **if** $b$ is frame **then**
        $(r, \tau) \leftarrow b$
        $A \leftarrow \text{Concat}(A, [\tau])$ ▷ Concatenate $C_\alpha$ coordinate.
    **else if** $b$ is atomized or ligand atom **then**
        $A \leftarrow \text{Concat}(A, b)$ ▷ Concatenate atomized residue or ligand coordinate.
    **end if**
**end for**
$\epsilon \sim \mathcal{N}(0, \mathrm{I}_3 \cdot \sigma_{\text{perturb}})$ ▷ Sample global translation.
$\tilde{B} \leftarrow (B - \mu(A)) + \epsilon$ ▷ - and + are broadcasted along the atom dim. and applied to the translation of frames.
**Return** $\tilde{B}$

Algorithm 2 first construct the global coordinate tensor $A$ that holds all the atom coordinates from the protein and ligand in $B$ (as a reminder in this section we are assuming $B = X$ for simplicity). In line 11, we subtract the global CoM $\mu(A)$ from both all the coordiantes followed by a global translation $\epsilon$ that is sampled from $\mathcal{N}(0, \mathrm{I}_3 \cdot \sigma^2_{\text{perturb}})$ for some $\sigma_{\text{perturb}} > 0$.

Stochastic centering causes the CoM of the input features to be noisy which forces RFdiffusion2 to not rely on CoMs when training. The issue of motif centering does not arise since $\mu\left(X^M\right) \neq 0$. The issue of motif placement in global centering does not arise either since $\mu(X^{(t)}) = t \cdot \mu(X^{(1)})$ where $\mu(X^{(1)}) \sim \mathcal{N}(\vec{0}, \mathrm{I}_3 \cdot \sigma_{\text{perturb}})$ according to line 11. Recall the issue was RFdiffusion2 would produce poor samples with certain initializations of $\mu(X^M)$ since RFdiffusion2 is trained to always keep $\mu(X^{(t)}) = 0$. With our new centering, RFdiffusion2 develops an inductive bias to move $\mu(X^{(t)})$ towards the location of the data point $X^{(1)}$ containing the motif. Stochastic centering also expands the distribution of $\mu(X^M)$ at training time to allow for increased robustness at inference time. First, we state the following proposition.

**Proposition 1.** *Suppose the following*

$$\nu \sim \mathcal{N}(\vec{0}, \mathrm{I}_3 \cdot \sigma_1^2), \qquad x_i|\nu \sim \mathcal{N}(\nu, \mathrm{I}_3 \cdot \sigma_2^2) \tag{35}$$

*where* $\vec{0} = [0, 0, 0]$. *The CoM (i.e. sample mean) for* $n$ *samples* $Y = [x_1, \dots, x_n]$ *follows the distribution*

$$\mu(Y) \sim \mathcal{N}\left(\vec{0}, \mathrm{I}_3 \cdot (\sigma_1^2 + n^{-1}\sigma_2^2)\right) \tag{36}$$

*Proof.* The proof is a straightforward exercise in multivariate statistics. Starting with the mean,

$$\mathbb{E}\left[\mu(Y)\right] = \mathbb{E}\left[\mathbb{E}\left[\mu(Y)|\nu\right]\right] = \mathbb{E}[\nu] = \vec{0}. \tag{37}$$

Now consider each dimension of $\mu(Y) = [y_1, y_2, y_3]$ where

$$y_j = \frac{1}{n}\sum_{i=1}^{n}(x_i)_j \text{ for } j \in \{1, 2, 3\} \tag{38}$$

we use $(x_i)_j$ as our notation for taking the $j$th index of $x_i$. The variance can be calculated using law of total variance

$$\begin{aligned} \mathrm{Var}(y_j) &= \mathbb{E}[\mathrm{Var}(y_j|\nu)] + \mathrm{Var}(\mathbb{E}[y_j|\nu]) && (39)\\ &= \frac{\mathbb{E}[\sigma_2^2]}{n} + \mathrm{Var}(\nu) && (40)\\ &= \frac{\sigma_2^2}{n} + \sigma_1^2 && (41) \end{aligned}$$

Similarly, covariance can be calculated with law of total covariance

$$\mathrm{Cov}(y_i, y_j) = \mathbb{E}[\mathrm{Cov}(y_i, y_j|\nu)] + \mathrm{Cov}(\mathbb{E}[y_i|\nu], \mathbb{E}[y_j|\nu]) \tag{42}$$

$$= \mathbb{E}[0] + \text{Cov}(\nu_i, \nu_j) \tag{43}$$

$$= 0 \tag{44}$$

where $\nu_i, \nu_j$ are the $i$th and $j$th index of $\nu$ respectively. Equation (41) and eq. (44) allows us to conclude the covariance matrix of $\mu(Y)$ is given by $\text{I}_3 \cdot (\sigma_1^2 + n^{-1}\sigma_2^2)$. □

To apply proposition 1, assume each $x_i$ in $X$ follows a multivariate gaussian distribution after stochastic centering

$$x_i|\mu(X) \sim \mathcal{N}(\mu(X), \text{I}_3 \cdot \sigma^2) \tag{45}$$

then it follows

$$\mu(X^M) \sim \mathcal{N}(0, \text{I}_3 \cdot (\sigma_{\text{perturb}}^2 + m^{-1}\sigma^2)). \tag{46}$$

where $m$ is the number of motif residues. Equation (46) explains that stochastic centering increases the variance of the motif residue's CoM distribution which allows for RFdiffusion2 to see a larger range of motif placements at training time depending on $\sigma_{\text{perturb}}$. The increased variance of the motif allows for increased robustness to different motif initializations at inference time. We allow users to specify an Origin token that we call `ORI` (discussed in appendix B.7) which specifies the origin at which noise is initialized. We found enzyme designers will use the `ORI` token to run sampling with different motif CoM initializations and discover diverse samples. RFdiffusion2 is capable of handling `ORI` tokens distributed approximately $\mathcal{N}(0, \sigma_{\text{perturb}}^2)$ since RFdiffusion2 is trained to move $\mu\left(X^{(t)}\right)$ towards the motif CoM.

Our gaussianity assumption is a heuristic necessary for our analysis but is still insightful and can be approximately accurate in high dimensions (which large biomolecules arguably fall under). Throughout this section, we have claimed the neural network architecture of RFdiffusion2 is able to learn different displacement vectors between CoMs. We have not provided proof for this claim since it requires delving into formal arguments with equivariant neural networks and universal neural network approximation theory that is out of the scope for our work. We empirically observed RFdiffusion2 is able to exploit *any features that give any clue* about the location of the ground truth. Stochastic centering is designed to train RFdiffusion2 in such a way that it does not learn to exploit SE(3) equivariant features that give privileged information present when training but not necessarily when sampling.

#### B.4 Features

The features used in RFdiffusion2 closely follows that of RFAA [19]. Each training example is taken from one of the biomolecule categories described in the table below. First column describes the data category. The second and third column show the number of clusters based on sequence similarity and number of total examples in the dataset respectively. The last column is the probability of the dataset being sampled for each training example.

| Biomolecule Category | Sequence Clusters | Examples | Probability |
|---|---|---|---|
| Protein Monomer | 13,872 | 203,101 | 40% |
| Protein Heteromer | 14,307 | 193,257 | 20% |
| Protein Small Molecule | 5,475 | 114,782 | 15% |
| Protein Metal Complex | 5,324 | 112,453 | 1% |
| Protein Multi-Residue Ligand | 532 | 4,324 | 5% |
| Protein Small Molecule Assembly | 2,455 | 41,313 | 5% |
| Covalent Modification | 1,029 | 11,912 | 5% |

Once a training example is selected, we process and construct a set of features extracted from a training example. In the following sub sections, we provide high-level descriptions of each feature (Appendices B.4.1 to B.4.3) and refer to the code for how each feature is computed. We will describe how different structure prediction inputs from RFAA were adapted for RFdiffusion2. As noted in appendix B.2.2, the features described in this section are captured by the variable $C$ which we will reference in pseudo code and in other sections. Once all the features have been described, appendix B.4.4 will give the variable names used in the code that hold all the features as pytorch tensors used as inputs into RFdiffusion2.

##### B.4.1 Sequence Features

In structure prediction, the sequence features are provided in the form of multiple sequence alignments (MSA) which use genetic databases to augment the primary predicted sequence with related sequences. When doing protein design, we do not know what the target sequence is (in entirety). The RFAA architecture allows for a variable number of aligned sequences, so for RFdiffusion2 we provide a single sequence in all cases.

| Feature | RFAA Description | RFdiffusion2 Description |
|---|---|---|
| Clustered MSA | Used to provide a clustered version of the MSA to the model | Provide the sequence of the motif residues and MASK tokens for the other positions |
| Clustered MSA Profile | Used to provide the frequency of the each position in the overall MSA | Provide an one-hot encoding of the motif sequences and MASK tokens for the other positions |
| Insertion Statistics | Two features on whether there are insertions at that position and how many there are | Always initialized to zeros |

| | | |
|---|---|---|
| Termini Annotations | Two binary features indicating whether a residue is a N terminus and whether a residue is a C terminus. This distinguishes the task of predicting a full protein at inference time and predicting a cropped protein in training | Same as structure prediction |

##### B.4.2 Bond and Positional Encodings

In the RFAA architecture, there is an explicit feature of bond order (single, double, triple and aromatic). This is used to create a positional embedding that is permutation invariant for small molecules and breaks permutation symmetry for proteins. The positional embedding also handles the case where a residue in the polypeptide chain is *atomized* where it's location in the sequence is represented to the model. This feature remains unchanged from RFAA but it used in RFdiffusion2 when residues are atomized and the rotamer is inferred.

| **Feature** | **RFAA Description** | **RFdiffusion2 Descriptions** |
|---|---|---|
| Residue Index | Used to break permutation symmetry for polymers. Feature is provided to network for non-polymers but does not change the operations of the network | Same as structure prediction. |
| Bond Order Matrix | Used to provide the bond order (single, double, triple, aromatic) or protein-atom bonds to the network | Same as structure prediction, with the addition of an extra bond token between unindexed residues and indexed residues |
| Bond Distance Matrix | Feature indicating the number of bonds that would need to be traversed between two atoms in a bonded system. Used to break symmetry for different atoms within an atomized residue. | Same as structure prediction |
| Chiral Center Definitions | All orderings of 4 atoms around a chiral center (first four dimensions) and the ideal pseudo-dihedral angle formed by that ordering of atoms (fifth dimension). | Same as structure prediction |

| | | |
|---|---|---|
| Atom Frames | Indices that form frames for each atom node in the input. These are provided as $l1$ feature biases to the structure module | Same as structure prediction |

##### B.4.3 Diffusion and Structure Features

In structure prediction, template structures of homologous sequences are provided to the network to improve structure prediction in cases where a homologous structure has been solved. In RFdiffusion2, we reuse the architectural components that are used to provide the template in structure prediction to provide the noisy diffusion structure. n these inputs, we diverge from structure prediction and provide features that are not present in the structure prediction training.

| **Feature** | **RFAA Description** | **RFdiffusion2 Descriptions** |
|---|---|---|
| Template Sequence | Sequence of the homologous template structure. | Motif sequence and unknown tokens for all diffused regions. |
| Template Confidence | Alignment confidence of the template sequence. | Not used. |
| Atom-wise RASA | Not used. | A feature that specifies each atom's RASA. |
| Timestep $t$ Sinusoidal Embedding | Not used. | Sin and cosine embedding of the timestep concatenated as in [9] |
| Noised Biomolecule $B^{(t)}$ | Not used. | Noisy coordinates and frames of the biomolecule. |
| Motif Biomolecule $B^M$ | Not used. | Motif residues and ligand atoms used as the fixed compoments of the structure. |

##### B.4.4 Tensor inputs

The previous sections discussed the features at a high-level, but we must convert these features into machine readable inputs to run RFdiffusion2. We list the pytorch features that the model sees in table 6. The first column describes the input name and shape and the second column describes the input features. Each input name matches the input in the code. We use $N_{\text{tot}}$ to refer to the total number of residues and atoms both indexed and unindexed.

| **Input (shape)** | **Description** |
|---|---|

| | |
|---|---|
| *msa_masked*<br>(1, $N_{\text{tot}}$, 164) | Sequence Input into model. Channels 0-79: Sequence (replacing MSA input). Channels 80-159: Sequence (replacing profile input). Channel 160-161: Zeros (replacing insertion statistics). Channel 162-163: N terminus and C terminus features. |
| *msa_full*<br>(1, $N_{\text{tot}}$, 83) | Channel 0-79: Sequence (replacing full MSA input). Channel 80-81: Zeros (replacing insertion statistics). Channel 82-83: N terminus and C terminus features. |
| *seq*<br>($N_{\text{tot}}$) | The protein sequence and any atom tokens, including mask tokens. |
| *idx*<br>(L) | Residue index of each residue in the input. Nil for atom nodes. |
| *bond_feats*<br>($N_{\text{tot}}$, $N_{\text{tot}}$, 7) | Pairwise bond adjacency matrix. Pairs of residues are either single, double, triple, aromatic, residue-residue, residue-atom, other, or UNINDEXED_BOND, denoting an edge between an unindexed residue and an indexed residue. |
| *dist_matrix*<br>($N_{\text{tot}}$, $N_{\text{tot}}$) | Minimum amount of bonds to traverse between two nodes. This is 0 between all protein nodes. |
| *chirals*<br>($N_{\text{num\_chiral\_centers}}$, 5) | All orderings of 4 atoms around a chiral center (first four dimensions) and the ideal pseudo-dihedral angle formed by that ordering of atoms (fifth dimension). |
| *atom_frames*<br>($N_{\text{num\_atoms}}$, 3, 2) | Indices that form frames for each atom node in the input. The second dimension represents that there are three atoms in each frame. The third dimension represents an offset in the node dimension because atom frames go across nodes and the absolute index in the atom dimension. These are provided as biases in the SE(3) transformer layers as in [19] |
| *t1d*<br>(2, $N_{\text{tot}}$, 114) | 1D template feature. First, 79 represent the "sequence" (residue/atom types) of the motif structure. Channel 80 contains the diffusion timestep (as a float between 0 and 1). Channel 81-90 contains the Radius of Gyration conditioning (10 bins; binned evenly between 0,100 with a bin for over 100), Channel 91-94: RASA features (3 bins; binned evenly between 0 and 0.2), Channel 95-114: Sinusoidal Timestep Embedding (embedding dim:20, max_positions=10000). |
| *t2d*<br>(2, $N_{\text{tot}}$, $N_{\text{tot}}$, 68) | The first "template" contains distances and angles between residues and atoms in $B^{(t)}$ using the features used in [19]. The second "template" is always zeros and was left from legacy experiments using self conditioning. |
| *alpha_t*<br>(2, $N_{\text{tot}}$, 60) | Sidechain torsion angles from $B^{(t)}$ (20 angles x sin, cos and whether the angle exists in the structure for each residue). Usually 0s, unless there is a backbone motif with sidechain information provided as torsion angles. |
| *msa_prev*<br>($N_{\text{num\_clusters}}$, L, $C_m$ | Always set to None. In structure prediction, recycled MSA features. $C_m$=256 (number of 1D channels) |

| | |
|---|---|
| *pair_prev* ($N_{\text{tot}}$, $N_{\text{tot}}$, $C_p$) | Always set to None, In structure prediction, recycled pair features. $C_p$=192 (number of 2D channels) |
| *state_prev* ($N_{\text{tot}}$, $C_s$) | Always set to None, In structure prediction, recycled state features. $C_s$=32 (number of 3D $\ell$0 channels) |
| *xyz_prev* ($N_{\text{tot}}$, 36, 3) | $B^{(t)}$ structure is provided as coordinates. |
| *sc_torsions_prev* (L, 30) | Always set to 0s as there is no recycling. |

#### B.5 Architecture

The neural network architecture used in RFdiffusion2 closely resembles RosettaFold All-Atom (RFAA) and RFdiffusion All-Atom (RFdiffusionAA) [19]. We refer to those works for more details while providing a brief description here and the changes we made.

RFdiffusion2 follows the three tracks introduced in RosettaFold (RF) [55] where the neural network can accept 1D sequence information, 2D distance and bond information and 3D coordinate information. While we train the network from scratch, we choose to maintain the input tensor shapes of RFAA because the network architecture was optimized for the structure prediction inputs. Briefly, RFAA introduced the ability to model arbitrary small molecules with the following innovations:

- Adding tokens for all atoms found in small molecules in the PDB.
- Adding explicit bond feature inputs
- Adding chirality inputs which break 3D chiral symmetries
- Generalizing the positional encoding to break permutation symmetry for polymers but maintain the symmetry for small molecules
- Scaling the trunk attention heads and channels to improve performance
- Generalizing the structure module to operate on frames for polymer residues and global translations for ligand atoms

***Architectural Improvements***

We largely followed the RFAA architecture [19]. In training RFAA for structure prediction, we found there was training instability associated with the rotation prediction which has also been noted in AF2 [35]. We used the same heuristic solution that was used in AF2 where the gradients were stopped between rotation predictions in the network. This was exacerbated since in RFAA we were predicting a structure at every block in the network instead of the structure module at the end of the network as in AF2. In RFdiffusion2, we found that allowing the gradients to flow through rotations of the SE(3) Transformer blocks in the trunk of the network did not cause instability. We still stop the rotation gradients in the refinement blocks at the end of the network.

Following RFdiffusionAA, we zero out the rotation and translation updates for motif residues and atoms. In this case, since the motif is provided within the noisy coordinates, the motif coordinates will never move during the trajectory.

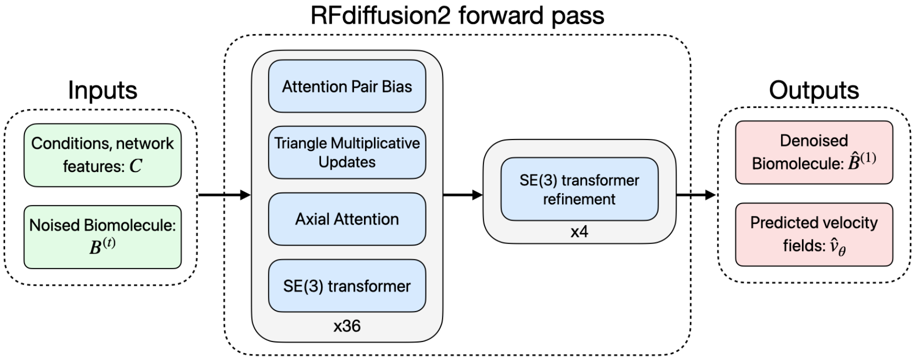

**Figure 10**: Diagram showing the neural network architecture of RFdiffusion2. The inputs are denoted with the noised biomolecule $B^{(t)}$ and $C$ which contains the conditions and network features that are discussed in appendix B.5. The middle of the diagram labeled RFdiffusion2 forward pass shows the neural network operations including Attention Pair Bias, Tri. Mult. Update, Axial Attention, SE(3) transformer, and SE(3) transformer refinement (see Krishna et al. [19] for details on each operation). We use plate notation to show the number of times each operation is performed. The outputs include the denoised biomolecule that is converted into the vector fields used in the flow matching loss eq. (28).

RFAA performs recycling of latent features as introduced in [35]. In RFdiffusion2, we do not use the recycling features and they are always initialized to zeros. This speeds up running the model.

#### B.6 Training

In this section, we discuss the training of RFdiffusion2. We first discuss the data processing pipeline (Appendix B.6.1) then the training procedure (Appendix B.6.2).

##### B.6.1 Training data pipeline

The training data uses the same set of structures as in RFdiffusionAA [19]. The steps to create a single training example are enumerated below and shown in fig. 11.

1. Sample a biomolecule category with probability from appendix B.4.
2. Sample a cluster from the biomolecule category by first computing the cluster probabilities as follows. First, compute the size of each cluster in the dataset. Then, clip the cluster sizes to be between 256 and 512 and divide by 512 to get the cluster weights.

$$\texttt{cluster\_weights} = \frac{1}{512}\text{Clip}(\texttt{clusters\_sizes}, \min = 256, \max = 512)$$

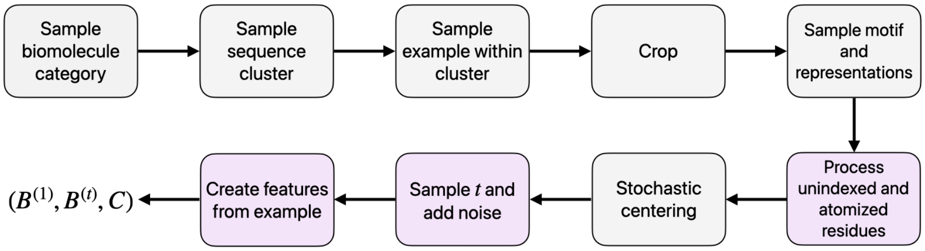

**Figure 11**: Flow chart of the training data pipeline. The final output $(B^{(1)}, B^{(t)}, C)$ is used as the training example for RFdiffusion2. Boxes in lilac are steps shared with the inference data pipeline.

Finally, sample a cluster using the cluster weights as the unnormalized probabilities. We reweight the cluster weights to avoid under or under sampling unbalanced clusters.

3. Sample a biomolecule from the cluster uniformly at random.
4. Crop each feature to a fixed size.
5. Sample which substructures are motifs and the representations of all residues. The procedure for sampling the motif and representations is discussed later in this section.
6. Process the unindexed and atomized residues. Using the masks in the previous step, we create a new biomolecule with the unindexed residues and atomized residues. Each residue is represented as a frame or atomized residue with the correct number of atoms. The unindexed residues are duplicated and given the correct labels they were duplicated from.
7. Perform stochastic centering (Algorithm 2).
8. Sample a noise time $t$ from a uniform distribution between 0 and 1 then noise the ground truth biomolecule up to $t$ following algorithm 1.
9. Compute the features described in appendix B.4 from the training example.
10. Return the ground truth biomolecule $B^{(1)}$, the noised biomolecule $B^{(t)}$, and $C$ which contains conditions and features.

The remainder of this section describes the sampling of the motif and representation in step 5 of the training data pipeline.

***Motif and Representation Sampling***

Our goal is to train RFdiffusion2 to handle any combination of motifs and representations that may be present in the input at inference time. Our strategy will be to randomly sample the motif and representation of the biomolecule during training such that a single biomolecule can be used multiple times by sampling different motifs and representations each time it is shown to RFdiffusion2. In the following table, we describe different functions that correspond to different motif and representation sampling functions. The name of each function corresponds to its name in the code. For each training example, we randomly select one function from the table and apply it to the biomolecule.

After selection of a motif, each residue of the motif is annotated as unindexed with probablity `P_IS_UNINDEXED` while atom RASA labels for the ligand are provided 50% of the time. The probability of each function will be given after the table.

| **Function** | **Description** |
|---|---|
| `get_unconditional_diffusion_mask_free_ligand` | No motif is used. All residues and atoms are generated. Residues are represented as frames unless the residue is a covalently modified residue. |
| `get_tip_mask_whole_ligand` | Uniformly at random choose 1 to 8 residues for which all heavy atoms are resolved. These residues will be atomized. For each, select an atom to be the seed atom for the motif: 80% of the time select the atom farthest from the backbone oxygen in the bond graph; 20% of the time select the seed atom uniformly at random. The seed atom selection is then expanded $n \sim \text{Geom}(p = 0.5)$ bonds outward to form a connected subgraph of atoms where Geom is the geometric distribution with success probability $p$. All ligand atoms are included in the motif. |
| `get_tip_mask_partial_ligand` | Perform `get_tip_mask_whole_ligand`. For each ligand select a fraction of the ligand that will be shown $p \sim \mathcal{U}(0, 1)$ and a seed atom uniformly at random. From the seed atom, perform a breadth-first traversal of the bond graph, adding each visited atom to the motif until the number of motif atoms $n$ in the $N$-atom ligand satisfies $\frac{n}{N} \geq p$. |
| `get_tip_mask_unconditional_free_ligand` | Perform `get_tip_mask_whole_ligand`. Keep the residue atomization selection, but include no residue atoms or ligand atoms in the motif. |
| `get_tip_mask_free_ligand` | Perform `get_tip_mask_whole_ligand`. Provide no ligand atoms as motif. |
| `get_tip_mask_bb_only_whole_ligand` | Uniformly at random choose 1 to 8 residues for which all heavy atoms are resolved. Provide each as a backbone motif. All ligand atoms are included in the motif. |

| | |
|---|---|
| get_tip_mask_bb_only_partial_ligand | Perform get_tip_mask_bb_only_whole_ligand. For each ligand select a fraction of the ligand that will be shown $p \sim \mathcal{U}(0, 1)$ and a seed atom uniformly at random. From the seed atom, perform a breadth-first traversal of the bond graph, adding each visited atom to the motif until the number of motif atoms $n$ in the $N$-atom ligand satisfies $\frac{n}{N} \geq p$. |
| get_tip_mask_bb_only_free_ligand | Perform get_tip_mask_bb_only_whole_ligand. Provide no ligand atoms as motif. |

We train two RFdiffusion2 models: a base model and an ablation model. The base model represents our best performing model trained on the most common motif and representation scenarios. The ablation model is trained with all the functions in order for us to study the different rotamer and index selection strategies (Figure 3E). P_IS_UNINDEXED is 80% for the Base model and 50% for the Ablation model.

| | **Sample probability** | |
|---|---|---|
| **Function** | **Base** | **Ablation** |
| get_unconditional_diffusion_mask_free_ | 10% | 10% |
| get_tip_mask_whole_ligand | 70% | 4.5% |
| get_tip_mask_partial_ligand | 0% | 28.35% |
| get_tip_mask_unconditional_free_ligand | 20% | 8.1% |
| get_tip_mask_free_ligand | 0% | 4.05% |
| get_tip_mask_bb_only_whole_ligand | 0% | 31.5% |
| get_tip_mask_bb_only_partial_ligand | 0% | 9% |
| get_tip_mask_bb_only_free_ligand | 0% | 4.5% |

The steps for sampling the motif and representation are as follows:

1. Sample a function to call from the table above based on whether the model is "Base" or "Ablation".
2. Call the function to sample the motif and representations.
3. Each residue is annotated as unindexed with probability P_IS_UNINDEXED, the value of which is based on whether the model is "Base" or "Ablation".
4. Sample $x \sim \text{Bern}(p = 0.5)$ where Bern is the Bernoulli distribution with probability $p$. If $x = 1$ then all ligand atoms are annotated with atom RASA labels. If $x = 0$ then no annoations are provided.
5. The above specifications are then passed to the next step in the pipeline where the biomolecule is processed according the motif and representation annotations including unindexed and RASA.

##### B.6.2 Training procedure

In this section, we describe the training procedure for RFdiffusion2. Training examples are generated using the training data pipeline described in appendix B.6.1. We represent the training data with $\mathcal{D}$ which is an iterator over tuples $(B^{(1)}, B^{(t)}, C)$. On each epoch, we set the random seed for the dataset to ensure that the order of the training examples is randomized.

For our optimization hyperparameters, we use the Adam optimizer [56] with learning rate 0.001, a scheduler that linearly ramps up the learning rate over 500 warmup steps and then decays by a factor of 0.95 every 5000 steps, l2 regularization coefficient of 0.01, 140 epochs, 25600 steps per epoch. The model was trained over several months, and when more GPUs became available we would resume training with higher batch sizes and make small adjustments based on observations of the model's performance. It is likely that a hyperparameter search would yield an improved model, but for completeness we detail the changes made here. Training proceeded in 3 stages: in stage 1 [epochs 0-63]: `batch_size` = 24, `P_UNINDEXED` = 0.5 (fraction of motifs provided as unindexed), `atom_loss_weight` = 1. In stage 2 [epochs 64-108], we increased the fraction of examples in which the motif was treated as unindexed, as the model sees disproportionately many indexed residues, followed AF3 [34] in upweighting loss on atoms by a factor of 10, and increased the batch size: `P_UNINDEXED` = 0.8, `atom_loss_weight` = 10, `batch_size` = 72. In stage 3 [epochs 109-140] we doubled the batch size: `batch_size` = 144.

Following Krishna et al. [19], we use gradient accumulation to maximize the size of the model we can fit in memory while training with a large batch size. This is done by accumulating gradients over $\xi$ steps before performing a single optimization step.

**Algorithm 3** Train

**Require:**

$N_{\text{epoch}}$ Number of epochs
$N_{\text{steps}}$ Steps per epochs
$N_{\ell}$ Number of SE(3) transformer layers
$\theta$ Model weights
$\gamma$ Layer decay
$\xi$ Batch size

$\theta \leftarrow \text{Init}(\theta)$ ▷ Randomly initialize model weights
$\theta_g \leftarrow 0$ ▷ Gradient accumulator
**for** $e$ in $[1, \dots, N_{\text{epoch}}]$ **do**
  SetRandomSeed($e$) ▷ Set random seed.
  $\mathcal{D}_e \leftarrow \text{Sample}(\mathcal{D}, N_{\text{steps}})$ ▷ Sample $N_{\text{steps}}$ training examples.
  **for** $s$, $B^{(1)}, B^{(t)}, C$ in $\mathcal{D}_e$ **do**
    $N_{\text{tot}} \leftarrow \text{Len}(B^{(t)})$ ▷ Number of elements in $B^{(t)}$
    **for** $i$ in $[1, \dots, N_{\text{tot}}]$ **do**
      **for** $l$ in $[1, ..., N_{\ell}]$ **do**
        $l \leftarrow \mathcal{L}_{\text{FM}}\left(b_i^{(t)}, b_i^{(1)}, B^{(t)}, C; \gamma, \ell\right)$
        $\theta_g \leftarrow \theta_g + \nabla_\theta l$ ▷ Gradient accumulation.
      **end for**
    **end for**
    **if** $\text{mod}(s, \xi) = 0$ **then**
      $\theta \leftarrow \text{TrainStep}(\theta_g)$
      $\theta_g \leftarrow 0$
    **end if**
  **end for**
**end for**
**Return** $\theta$

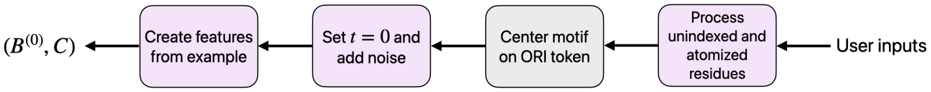

**Figure 12**: Flow chart of the inference data pipeline. The final output $(B^{(1)}, B^{(t)}, C)$ is used as the training example for RFdiffusion2. Boxes in lilac are steps shared with the training data pipeline.

##### B.7 Sampling

To generate new proteins with RFdiffusion2, the user provides a PDB file with the motif to scaffold; length of the protein to generation; number of discretization steps; and specification of the motifs and representations. Observe that during training, the motifs and representations were chosen and sampled at random from a training example (Appendix B.6.1) where at inference time the specification is set by the user. Our code provides utilities and functions to aid in specifying the motifs and representations corresponding to residue and atom identifiers in the PDB file.

The data processing pipeline for inference is shown in fig. 12 where we see the steps are similar to the training data pipeline except for a few key differences. Instead of stochastic centering, the user specifies a coordinate `ORI` in the PDB file to specify the CoM and orientation of the generated protein relative to the motif. The motif biomolecule is given by $B^M$. We perform the following to position the motif biomolecule with respect to `ORI`.

$$B^M \leftarrow \text{ApplyORI}(B^M, \texttt{ORI}). \tag{47}$$

Next, instead of randomly sampling $t$, we set $t = 0$ to initialize a biomolecule with pure noise $B^{(0)}$, except for the motif. This can be achieved by creating a placeholder biomolecule $B$ with an arbitrary initialization. We perform the noising step (Algorithm 1) on the biomolecule to create a noise biomolecule $B^{(0)}$.

$$B^{(0)} \leftarrow \text{NoiseBiomolecule}(B, 0). \tag{48}$$

As previously discussed in appendices B.2.2 and B.4, $C$ is used to hold all conditioning information and features used by the model other than $B^{(t)}$. This includes the motif $B^M$, ligand bond geometry, amino acid types, atom types, and RASA conditions. Inference is finally ran using algorithm 5

$$B^{(1)} \leftarrow \text{Inference}(B^{(0)}, C, \texttt{num\_steps}). \tag{49}$$

**Algorithm 4** ApplyORI

**Require:**

$B^M$ Motif biomolecule

`ORI` 3D coordinate to act as new center of mass

$B^M \leftarrow B^M - \mu\left(B^M\right)$ ▷ Center the motif where $\mu(\cdot)$ computes the CoM.
**for** $b_i$ in $B^M$ **do**
  **if** $b_i$ is atomized or ligand atom **then**
    $\tilde{b}_i \leftarrow b_i - \texttt{ORI}$ ▷ Broadcast `ORI` across atom dimension.
  **else if** $b_i$ is frame **then**
    $(r_i, \tau_i) \leftarrow b_i$ ▷ Separate rotation and translation component.
    $\tilde{\tau}_i \leftarrow \tau_i - \texttt{ORI}$ ▷ Only center translation.
    $\tilde{b}_i \leftarrow (r_i, \tilde{\tau}_i)$
  **end if**
**end for**
$\tilde{B}^M \leftarrow [\tilde{b}_1, \tilde{b}_2, \dots]$ ▷ ORI-centered motif.
**Return** $\tilde{B}^M$

**Algorithm 5** Inference

**Require:**

$B^{(0)}$ Noise initialization

$C$ Conditions and features

`num_steps` Number of discretization steps

$t \leftarrow 0$
$\Delta \leftarrow 1/\texttt{num\_steps}$
$\ell \leftarrow -1$ ▷ Use output of last SE(3)-transformer layer.
**while** $t < 1.0$ **do**
  **for** $b_i^{(t)}$ in $B^{(t)}$ **do**
    **if** $b_i^{(t)}$ is atomized or ligand atom **then**
      $b_i^{(t+\Delta)} \leftarrow b_i^{(t)} + \Delta \cdot \hat{v}_\theta^{\mathbb{R}}(b_i^{(t)}; B^{(t)}, C, \ell)$
    **else if** $b_i^{(t)}$ is frame **then**
      $(r_i^{(t)}, \tau_i^{(t)}) \leftarrow b_i^{(t)}$ ▷ Separate rotation and translation component.
      $r_i^{(t+\Delta)} \leftarrow \mathrm{Exp}_{r_i^{(t)}}\left(10 \cdot \Delta \cdot \mathrm{Log}_{r_i^{(t)}}\left(\hat{v}_\theta^{\mathrm{SO(3)}}(r_i^{(t)}; B^{(t)}, C, \ell)\right)\right)$
      $\tau_i^{(t+\Delta)} \leftarrow \tau_i^{(t)} + \Delta \cdot \hat{v}_\theta^{\mathbb{R}}(\tau_i^{(t)}; B^{(t)}, C, \ell)$
      $b_i^{(t+\Delta)} \leftarrow (\tau_i^{(t+\Delta)}, r_i^{(t+\Delta)})$
    **end if**
  **end for**
  $B^{(t+\Delta)} \leftarrow [b_1^{(t+\Delta)}, b_2^{(t+\Delta)}, \dots]$
  $t \leftarrow t + \Delta$
**end while**
**Return** $B^{(1)}$

### C Atomic Motif Enzyme (AME) details

#### C.1 Curation

We first discuss the curation of the AME benchmark. We downloaded the M-CSA's curated data flat file, containing 963 enzyme entries with accompanying PDB structures and sequences.

With 952 enzymes, we performed the filters listed below where each filter is sequentially applied. The left column indicates the order in which the filters were performed. The middle column gives the number of enzymes before and after the filter. The right column gives a brief description of the filter applied.

| **Order** | **Num. enzymes** | **Filter** |
|---|---|---|
| 1 | 963 → 952 | Parse errors, e.g. chain-misnumbering, missing cofactors |
| 1 | 952 → 915 | Remove enzymes with reactions at post translation modification (PTMs) or occurring on C- or N- terminus. |
| 2 | 915 → 357 | Enzyme must have a PARITY entry. |
| 3 | 357 → 90 | All reactants are present in corresponding PDB file. |
| 4 | 90 → 74 | Active site has at most 7 residues in the active site. |
| 5 | 74 → 60 | All active site residues must be <20Å from each other. |
| 6 | 60 → 45 | At least one ligand be <3Å from a active site residue. |
| 7 | 45 → 41 | All the ligands be <6Å from a active site residue. |

In filter (1), we choose to remove enzymes with active sites involving PTMs to simplify the benchmark and metrics to be based on canonical amino acids. Additionally reactions that require N- or C-terminal catalytic residues would put prior art at an unfair disadvantage, so in the interest of fairness we remove these as well. The most stringent filter is (2) which removes enzymes in which a close analog of each reactant and cofactor is not present in the crystal structure. In filter (4), we limit the number of motif residues to at most seven residues, as scaffolding even just 4 residue active sites was beyond the capabilities of prior state of the art. The purpose of filters (5) through (7) are to choose active sites that are compact by ensuring all the residues, reactants, and co-factors are within reasonable proximity of each other. These filters all serve to ensure that the theozymes extracted are not missing necessary catalytic components in order to capture a representative sample of realistic theozyme scaffolding problems.

The 41 enzymes are then processed to construct atomic motifs from their respective PDB files. We extract residue, reactant and co-factor identities from the M-CSA curated data flat-file. We then find the structure the active site by extracting the residues and ligands from the corresponding PDB file. This dataset does not describe the catalytically active atoms within each residue, so we must select them ourselves. Our aim with this benchmark is not to create minimal necessary and sufficient theozymes, but rather to capture the difficulty and functional form of the enzyme design task from theozymes. We do not believe that our selection of catalytic atoms would be more difficult to scaffold than the true set of catalytic atoms. For each motif residue, we find the atom in the side chain functional group that is closest in proximity to any of the ligands. Next, we take a subgraph of the side chain based on the graph distance to

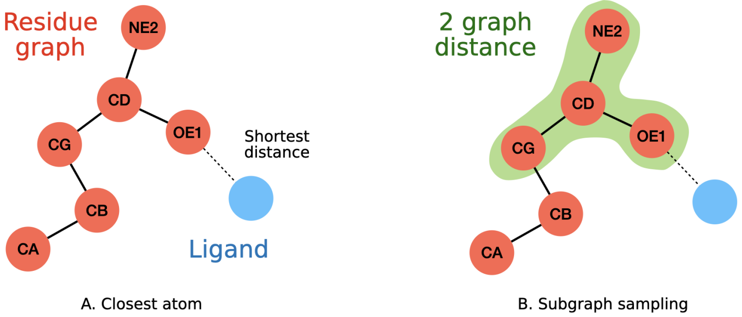

**Figure 13**: Depiction of motif subgraph sampling. We use Glutamine as an example for the residue subgraph drawn in red while an arbitrary ligand is drawn in blue. The atom names in the residue graph are the associated atom names as they would appear in PDB files. **A.** We find the atom in the residue graph that is closest to the ligand. In this case, it is OE1. **B.** We randomly sample a graph distance between one to three. In this case it is two and construct the subgraph around OE1. The subgraph is highlighted in green and becomes the atomic motif of this residue.

this closest atom where the covalent bonds are treated as the edges. We select between one to three uniformly at random as the geodesic distance. This subgraph becomes the atomic motif of the motif residue. A depiction of the motif sub graph extraction is shown in fig. 13. Every residue undergoes this procedure. In our benchmark, we provide the ligand as whole without any partial ligands. However, RFdiffusion2 is capable of partial ligands and so it is possible to extract and benchmark partial ligand performance.

A description of each enzyme case in AME is provided below. The first column contains the M-CSA ID that can be used to identify the enzyme in the M-CSA database[2]. The second column is a description of the enzyme found in its M-CSA entry. The third column is the corresponding PDB ID. The fourth column is the atom identities of the residues that participate in the atomic motif. For example, in the first row, A64 stands for residue number 64 on chain A of the structure in PDB 1nzy. [O, C] stands for the the O and C atoms in residue A64. The atom names correspond to the atom identities of the residue's amino acid identity. The fifth column is the list of ligands made of small molecules and co-factors. The sixth column is the number of residue islands which correspond to the number of residue chain segments. For example, if residues A32, A33, A50, A51, A52 are listed in the fourth column, then there are two residue islands. One can also think of residue islands as the number of contiguous residue segments in the motif. We find that residue islands indicate the difficult of the task with more islands being a harder, more complicated motif to scaffold.

[2]See urlhttps://www.ebi.ac.uk/thornton-srv/m-csa/browse/

| M-CSA ID | Enzyme name | PDB ID | Selected motif residue atoms | Ligands | Residue islands |
|---|---|---|---|---|---|
| M0024 | 4-chlorobenzoyl-CoA dehalogenase | 1nzy | A64: [O, C]; A86: [CB, CA, N, C]; A90: [CE1, ND1, NE2, CG, CD2]; A114: [N, CA]; A137: [NE1, CD1, CE2, CG, CD2, CZ2]; A145: [OD2, CG, CB, OD1]; | BCA | 6 |
| M0040 | phosphoglycerate kinase | 13pk | A39: [NH1, CZ, NE, NH2]; A219: [NZ, CE, CD]; A376: [N, CA, C]; A399: [N, CA]; | ADP, MG, 3PG | 4 |
| M0050 | orotidine-5'-phosphate decarboxylase | 1dbt | A33: [NZ, CE]; A60: [OD2, CG, CB, OD1]; A62: [NZ, CE]; A123: [OG1, CB, CA, CG2]; A215: [N, CA, C, CB]; B65: [OD2, CG]; | U5P | 6 |
| M0054 | 3-dehydroquinate dehydratase (type I) | 1qfe | A86: [O, C, CA]; A143: [NE2, CD2, CE1, CG, ND1]; A170: [NZ, CE, CD]; | DHS | 3 |
| M0058 | adenylate cyclase | 1cju | A396: [OD2, CG]; A397: [O, C, CA]; A440: [OD2, CG]; B1029: [NH1, CZ, NE, NH2]; B1065: [NZ, CE, CD]; | DAD, MG | 4 |
| M0078 | citrate (Si)-synthase | 1al6 | A244: [OG, CB]; A274: [ND1, CG, CE1, CB, CD2, NE2]; A320: [NE2, CD2, CE1]; A329: [NH2, CZ, NE, NH1]; A375: [OD2, CG, CB, OD1]; | OAA, HAX | 5 |

| | | | | | |
|---|---|---|---|---|---|
| M0092 | UDP-glucose 6-dehydrogenase | 1dli | A118: [N, CA, C, CB]; A145: [N, CA]; A204: [NZ, CE, CD]; A208: [OD1, CG, CB, ND2]; A260: [SG, CB, CA]; A264: [OD1, CG, CB, OD2]; | UDX, NAD | 6 |
| M0093 | hydroxymethylglutaryl-CoA reductase (NADPH) | 1dqa | A691: [NZ, CE, CD]; A767: [OD2, CG, CB, OD1]; B559: [OE2, CD]; B866: [CD2, CG, NE2, CB, ND1, CE1]; | NAP, COA | 4 |
| M0096 | creatinase | 1chm | A232: [NE2, CD2, CE1]; A262: [OE2, CD]; A358: [OE1, CD]; | CMS | 3 |
| M0097 | cytidine deaminase | 1ctt | A102: [ND1, CG, CE1]; A104: [OE1, CD]; A129: [SG, CB]; A132: [SG, CB, CA]; | DHZ, ZN | 4 |
| M0110 | D-amino-acid oxidase | 1c0p | A1054: [N, CA]; A1335: [O, C, CA]; A1339: [N, CA]; | DAL, PER, FAD | 3 |
| M0129 | taurine dioxygenase | 1os7 | A99: [NE2, CD2, CE1]; A101: [OD1, CG, CB, OD2]; A255: [NE2, CD2, CE1, CG, ND1]; A270: [NH1, CZ]; | AKG, FE2, TAU | 4 |
| M0151 | 2-amino-4-hydroxy-6-hydroxymethyldihydropteridine diphosphokinase | 1q0n | A82: [NE, CD, CZ]; A92: [NH2, CZ, NE, NH1]; A95: [OD1, CG]; A97: [OD2, CG]; | APC, PH2, MG | 4 |
| M0157 | hydroxyacylglutathione hydrolase | 1qh5 | A54: [NE2, CD2, CE1]; A56: [ND1, CG, CE1, CB, CD2, NE2]; A58: [OD2, CG]; A59: [NE2, CD2, CE1]; A110: [NE2, CD2, CE1]; A134: [OD2, CG]; A173: [NE2, CD2, CE1]; | GSH, ZN | 6 |

| | | | | | |
|---|---|---|---|---|---|
| M0179 | chaperonin ATPase | 1q3s | A64: [OD1, CG, CB, OD2]; A97: [N, CA, C, CB]; A98: [N, CA, C, CB]; A393: [OD2, CG, CB, OD1]; | ADP, MG | 3 |
| M0188 | UDP-glucose 4-epimerase | 1xel | A124: [OG, CB, CA]; A149: [OH, CZ, CE1, CE2]; A153: [NZ, CE]; | UPG, NAD | 3 |
| M0209 | adenosine kinase | 1lij | A136: [NE, CD, CZ, CG, NH1, NH2]; A318: [OD2, CG]; | RPP, ACP, MG | 2 |
| M0255 | alcohol dehydrogenase (SDR type) | 1mg5 | A108: [ND2, CG]; A139: [OG, CB, CA]; A152: [OH, CZ]; A156: [NZ, CE, CD]; | NAI, ACT | 4 |
| M0315 | enoyl-CoA hydratase | 1ey3 | A98: [N, CA]; A141: [N, CA]; A144: [OE1, CD, CG, OE2]; A164: [OE2, CD]; | DAK | 4 |
| M0349 | delta 5-3-ketosteroid isomerase | 1e3v | A16: [OH, CZ, CE1, CE2]; A40: [OD2, CG]; A100: [N, CA, C, CB]; A103: [OD2, CG]; | DXC | 4 |
| M0365 | Phosphofructokinase I | 1pfk | A11: [N, CA]; A72: [NH2, CZ, NE, NH1]; A125: [OG1, CB, CA, CG2]; A127: [OD2, CG]; A129: [OD2, CG]; A171: [NH1, CZ, NE, NH2]; | ADP, FBP, MG | 6 |
| M0375 | Purine nucleoside phosphorylase DeoD-type | 4ts9 | A20: [N, CA]; A24: [CD, CG, NE, CB, CZ]; A87: [NH2, CZ]; A90: [OG, CB]; A204: [OD1, CG]; A217: [NH2, CZ, NE, NH1]; C43: [NH1, CZ]; | PO4, FMC | 7 |

| | | | | | |
|---|---|---|---|---|---|
| M0500 | alcohol dehydrogenase (class II) | 1e3i | A46: [SG, CB]; A47: [CD, N, CG]; A48: [OG1, CB]; A67: [NE2, CD2, CE1]; A178: [SG, CB, CA]; A182: [OG, CB]; | CXF, NAI, ZN | 4 |
| M0552 | aconitate hydratase | 1fgh | A100: [OD2, CG]; A101: [NE2, CD2, CE1]; A165: [OD1, CG, CB, OD2]; A447: [NH1, CZ]; A642: [OG, CB, CA]; A644: [NE, CD, CZ]; | ATH | 5 |
| M0555 | L-amino-acid oxidase | 1f8r | A223: [NE2, CD2, CE1, CG, ND1]; A326: [CE, CD, NZ]; | CIT, FAD | 2 |
| M0584 | L-lactate dehydrogenase | 1ldm | A106: [NE, CD, CZ]; A166: [OD1, CG]; A169: [NH2, CZ]; A193: [NE2, CD2, CE1]; | NAD, OXM | 4 |
| M0630 | dihydroorotase | 1j79 | A250: [OD1, CG]; | ORO, ZN | 1 |
| M0636 | cytosine deaminase (yeast) | 1uaq | A64: [OE2, CD, CG, OE1]; A89: [O, C]; A91: [SG, CB]; | DUC, ZN | 3 |
| M0663 | ribokinase | 1rk2 | A252: [N, CA]; A253: [CB, CA]; A254: [N, CA]; A255: [OD2, CG, CB, OD1]; | ADP, RIB | 1 |
| M0664 | dihydroneopterin aldolase | 2dhn | A22: [OE2, CD]; A100: [NZ, CE, CD]; | PH2 | 2 |
| M0674 | N-carbamoyl-D-amino-acid hydrolase | 1uf7 | A46: [OE1, CD, CG, OE2]; A126: [NZ, CE, CD]; A171: [CB, CA, N, C]; | CDV | 3 |
| M0710 | cytosine deaminase (bacterial) | 1ra0 | A156: [OE1, CD]; A217: [OE2, CD]; | FE, FPY | 2 |
| M0711 | glyceraldehyde-3-phosphate dehydrogenase (NADP+) | 2esd | A154: [ND2, CG]; A250: [O, C, CA]; A284: [SG, CB, CA]; | G3H, NAP | 3 |

| | | | | | |
|---|---|---|---|---|---|
| M0717 | ornithine cyclodeaminase | 1x7d | A56: [OE1, CD, CG, OE2]; A228: [OD2, CG, CB, OD1]; | ORN, NAD | 2 |
| M0731 | fatty acid amide hydrolase | 1mt5 | A142: [NZ, CE]; A217: [OG, CB, CA]; A218: [N, CA]; A238: [N, CA, C, CB]; A239: [N, CA, C]; A240: [N, CA]; A241: [OG, CB]; | MAY | 3 |
| M0732 | dCTP deaminase | 1xs1 | A124: [O, C]; A126: [O, C]; A138: [OE1, CD, CG, OE2]; C111: [N, CA, C, CB]; C115: [NH2, CZ, NE, NH1]; | DUT | 5 |
| M0738 | phosphoglycerate mutase (2,3-bisphosphoglycerate-independent) | 1o98 | A62: [OG, CB, CA]; A154: [OD1, CG]; A261: [NH1, CZ]; | 2PG, MN | 3 |
| M0739 | L-aspartate oxidase | 1knp | A290: [NH2, CZ, NE, NH1]; A351: [CE1, ND1, NE2]; A386: [CD2, CG, CB, CD1]; | SIN, FAD | 3 |
| M0870 | acetylglutamate kinase | 1oh9 | A8: [NZ, CE, CD]; A11: [N, CA, C]; A45: [N, CA, C]; A162: [OD2, CG, CB, OD1]; A217: [NZ, CE]; | ADP, MG, NLG | 5 |
| M0904 | 3'(2'),5'-bisphosphate nucleotidase | 1qgx | A49: [OD2, CG, CB, OD1]; A72: [OE1, CD, CG, OE2]; A142: [OD1, CG, CB, OD2]; A144: [O, C, CA]; A145: [CA, N, C, CB, O, CG]; A147: [OG1, CB]; A294: [OD2, CG]; | MG, AMP | 6 |
| M0907 | ribulose-bisphosphate carboxylase (type I) | 1rbl | A175: [NZ, CE, CD]; A201: [NZ, CE]; A203: [OD1, CG, CB, OD2]; A204: [OE1, CD, CG, OE2]; A294: [NE2, CD2, CE1, CG, ND1]; A334: [NZ, CE, CD]; | MG, FMT, CAP | 5 |

#### C.2 Evaluation

To evaluate a motif-scaffolding method, we first process each case of AME to work with the input specification as discussed later in this section and perform inference with 100 Neural Function Evaluations (NFEs) also known as the number of steps used during inference. We sample 100 scaffolds for each motif then use LigandMPNN to sample 8 sequences for each scaffold and motif structure. When running LigandMPNN, we fix the motif amino acid identities to their amino acid found in their respective PDB file. We use a temperature of 0.1 in LigandMPNN.

For each sequence, we use Chai-1 with weights downloaded from their github repository[3], specifically version 0.0.1, to predict the all-atom biomolecule of the ligand and protein. We run Chai-1 single sequence mode, i.e. without MSAs, structural templates, or constraints. Five diffusion samples are outputted for each sequence and the highest confidence structure is chosen. For each design, there are 8 Chai-1 predicted structures which we further restrict to structures without steric clashes in the atomic motif. We then compute the all-atom RMSD of the atomic motif between a design and each predicted structure. We take the minimum all-atom RMSD and consider a success if the all-atom motif RMSD is $<1.5$Å and if there are no steric clashes in predicted atomic motif.

Our metrics closely resemble those used in the previous RFdiffusion benchmark where instead the RMSD is taken over the motif backbone atoms; ProteinMPNN is used for sequence generation; AlphaFold2 is used as the structure prediction method. We have already argued in section 1 and section 3 the limitations of the RFdiffusion benchmark as being too coarse by using backbone atoms as the motif which don't necessarily participate in the enzyme reaction. We have already justified the choice of Chai-1, a re-implementation of AlphaFold3, over using the traditional choice of AlphaFold2. In total, a total of (41 cases)$\times$(10 designs)$\times$(8 sequences) designs are generated for each method.

##### C.2.1 RFdiffusion protocol

In order to evaluate RFdiffusion on AME, we first must transform the unindexed tip atom motifs into the indexed, full-residue motif expected by RFdiffusion. To generate full-residue motifs from side chain functional groups, we sample inverse rotamers for each motif residue using the `invrotzyme` tool (https://github.com/ikalvet/invrotzyme), following a similar approach to previous works that utilize RFdiffusion for *de-novo* enzyme design [1]. The workflow utilizes an enzyme constraint file that specifies geometric relationships in the form of distances, angles, torsions and their associated tolerances and sampling levels between pairs of motif residues or between the ligand and motif residues, defined using triplets of atoms for each component (described in more detail in https://docs.rosettacommons.org/docs/latest/rosetta_basics/file_types/match-cstfile-format). When the atomic motif contains only one or two atoms for a given residue, we specify additional atoms by selecting the nearest neighboring atom(s) in the associated residue to complete the constraints required by the protocol. We set the tolerance values for these constraints to 0 and sampling

[3] https://github.com/chaidiscovery/chai-lab

levels to 1, ensuring that the sampled residues maintain the exact side chain functional group positions specified in the atomic motif. For the ligand, we fix its position from the input file and select 3 atoms closest to the first residue being sampled to make the first constraint block. The `invrotzyme` script selects all rotamers from the Dunbrack library whose cumulative probability equals or exceeds a specified threshold (selected to be 0.9) and then samples the selected rotamers with equal probability during inverse-rotamer generation. In addition, the tool also samples backbone positions in helical and strand conformations using Rosetta to accommodate these rotamers. For each benchmarking case, we establish a random order for processing the motif residues at the beginning of the protocol and build each residue sequentially while constraining its position using the geometric constraints defined in the constraint files. The outputs are filtered to eliminate structures with steric clashes between the ligand and motif residues or between different motif residues. We explore all combinations of secondary structure assignments (helix or strand) for each motif residue, resulting in $2^n$ total combinations where $n$ is the number of residues. We then draw samples from each of these combinations with equal probability until we obtain a total of 100 candidates, which become our final set of input motifs to RFdiffusion.

To assign sequence indices to the motif islands, we sample indices uniformly at random across the length of the protein for each design. Whereas RFdiffusion2 does not receive the privileged information that 2 motif residues are part of an island (i.e. contiguous in the native enzyme), in the interest of a fair comparison we give this information to RFdiffusion, as it is in some cases visually obvious from the catalytic residue motif atoms that the two residues would have to be abutting in primary sequence. We then perform inference with 100 timesteps, using the same settings used in the RFdiffusion enzyme-motif scaffolding evaluations [12] which employs an attractive-repulsive ligand potential.

##### C.2.2 RFdiffusion2 protocol

RFdiffusion2 accepts the unindexed atomic motifs present in AME in their native form, using `ORI` tokens placed at the center of mass of the motif atoms.

### D *In vitro* experimental method

#### D.1 Retroaldolase

##### D.1.1 Design

We extract a minimal theozyme from the evolved retroaldolase: RA95.5-8F (PDB: 5AN7). The constellation of catalytic atoms in the active site consists of the (CZ, OH) of Y1051; (NZ,CE) of K1083; (OD1, CG, ND2) of N1180; and (CZ, OH) of Y1180. For the reactant we use the suicide inhibitor (CCD:LLK) present in the PDB, as it has sufficient homology to our desired substrate methadol. Together the reactant and these 9 active site atoms form the entirety of the motif. The ORI token was placed at the CoM of the active site atoms. We generated 20,000 structures for each of 6 conditions corresponding to all combinations of (protein length $\in [120, 150, 180]$) and (ligand RASA $\in [0, \emptyset]$). For each of the resulting designs we fit 8 sequences with LigandMPNN, configured to exclude lysine from the set of possible amino acids to ensure the motif lysine is the the only lysine int he structure, for a total of 960,000 sequences.

First we filter on the self-consistency of the active site (motif) by retaining only sequences that possess motif all-atom RMSD $<$ 1.2A between the design structure and the structure predicted from sequence alone made by AF2. The resulting 9826 designs were the filtered on SASA of the catalytic Lysine to ensure its burial, and in turn, the probability of it existing in its deprotonated state. We then ensure that the reactant has a sufficient degree of burial in the active site by filtering on the SASA of the reactant. Additionally, we filter on the global self consistency of the backbone. After applying these three filters (LysineSASA $<$ 5 AND reactantSASA $<$ 120 AND global backbone RMSD $<$ 2A) we are left with 2724 sequences.

Finally, the set of 2724 designs was subjected to analysis with PLACER [39]. For each design we generated six structural ensembles of 50 models each for apo structure of the design, holo structure of non-covalent enzyme-substrate complex and structures of four possible stereoisomers of carbinolamine intermediate fig. 14. For each of these six ensembles we computed the following metrics: average side chain heavy atoms RMSD from the design for each theozyme residue (`design_rmsd_res`{`1,2,3,4`}), intra-ensemble average side chain RMSF (`intra_rmsf_res`{`1,2,3,4`}), fraction of the models in which substrate carbonyl and hydroxyl oxygens (Y1051 and Y1180) are positioned to form hydrogen bonds to respective active site residues(`hbond_fraction_res`{`2,3`}). These 10 metrics were then averaged across all six ensembles (with the exception the apo structure for (`hbond_fraction_res`{`2,3`}) as it is not relevant in that context. In these filters res1, res2, res3, res4 correspond to catalytic residues K1083, Y1051, Y1180, and N1110 respectively. We applied the following filter values appendix D.1.1 to select 90 designs for experimental characterization.

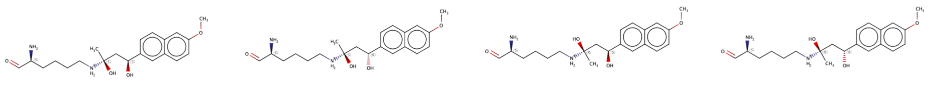

(S,S) (S,R) (R,S) (R,R)

**Figure 14: Stereoisomers of carbinolamine intermediate of retroaldolase reaction**

| Filter | Threshold |
|---|---|
| `design_rmsd_res1` | < 0.95Å |
| `design_rmsd_res2` | < 1.26Å |
| `design_rmsd_res3` | < 1.25Å |
| `design_rmsd_res4` | < 1.02Å |
| `intra_rmsf_res1` | < 0.34Å |
| `intra_rmsf_res2` | < 0.92Å |
| `intra_rmsf_res3` | < 0.93Å |
| `intra_rmsf_res4` | < 0.28Å |
| `hbond_fraction_res2` | ≥ 38% |
| `hbond_fraction_res3` | ≥ 18% |

##### D.1.2 Experimental setup

We used a protocol for medium throughput semi-quantitative retro-aldolase activity using racemic methodol for testing of the designs using in vitro transcription translation system (IVTT). We obtained linear duplex DNA, encoding protein of interest, flanked by appropriately spaced T7 promoter and T7 terminator sequences as eBlocks™ gene fragments from IDT, and by mixing 1μL of 4 ng/$\mu$L eblock solution directly with 4 $\mu$L components of PurExpress IVTT system produced enough protein to screen for active variants. After incubating for 4 hours at 37°C, 10 $\mu$L of phosphate buffered saline (PBS) was added and 10 $\mu$L of the mixture was used to test for retro-aldolase activity by mixing with 10 $\mu$L of 800 $\mu$M racemic methodol in PBS with 2% residual acetonitrile. It should be noted that in this format of the experiment, concentration of the soluble protein remains unknown and therefore activity level should be treated as apparent activity defined as a product of the amount of soluble protein and intrinsic activity of each variant. It is possible to make assay more quantitative by fusing protein of interest to the fluorescent protein reporter at the expense of decreased amount of retro-aldolase catalyst synthesized. Control proteins with known catalytic efficiencies: (RAbB-16.2 [$\frac{k_{\text{cat}}}{K_M} = 500\ \text{M}^{-1}\text{s}^{-1}$][57], RA110.4-6 [$\frac{k_{\text{cat}}}{K_M} = 55\ \text{M}^{-1}\text{s}^{-1}$][58]) were included to assist in ranking activity of the designs.

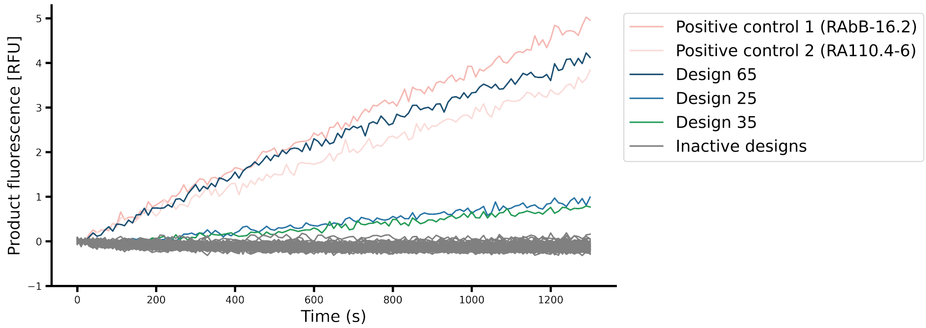

**Figure 15**: Retroaldolase IVTT sem-quantitative activity characterization. The RFU versus time of the tested 90 designs, 3 of which show activity. In order to compare the rate of product formation, the RFU values for each well have the background RFU (as measured by the the mean RFU for the first 3 timepoints) subtracted.

#### D.2 Cysteine Hydrolase

Cysteine hydrolases catalyze hydrolysis through a double-displacement mechanism utilizing a thiol nucleophile. This multi-step mechanism begins with nucleophilic attack by the activated cysteine thiolate on the substrate's carbonyl carbon, forming a tetrahedral intermediate that collapses to an acyl-enzyme intermediate with release of the alcohol product. This acyl-enzyme intermediate is subsequently attacked by a water molecule activated by the histidine, forming a second tetrahedral intermediate that resolves to release the carboxylic acid product and regenerate the free enzyme. In native cysteine hydrolases such as *Papain*, this reaction is facilitated by the Cys-His-Asn catalytic triad, where histidine acts as a general acid/base and the tetrahedral intermediates are stabilized by an oxyanion hole composed of the cysteine backbone nitrogen and a glutamine side chain. The spatial arrangement of these functional groups is critical for catalysis, likely requiring subangstrom level precision in their relative positions and orientations in the designed enzymes.

##### D.2.1 Design

We developed a computational pipeline that involved scaffold generation using RFdiffusion2 followed by two stages of sequence design for the *de novo* design of cysteine hydrolases. We utilized the crystal structure of native Papaya cysteine hydrolase (PDB ID: `1PPN`) to construct a precise catalytic motif by modeling the first tetrahedral intermediate with 4MU-Butyrate substrate. Using RDKit, we generated a tetrahedral conformer at the carbonyl carbon and oriented it according to established geometric parameters for optimal nucleophilic attack ($\alpha_{\text{cat}} \sim 76°$) and proton transfer ($\chi_{\text{oxy}} \sim 30°$), as described in Buller and Townsend [44], in addition to ensuring optimal distance and orientation to Cys25's backbone amide nitrogen ($N$) and Gln19's side chain nitrogen ($N\varepsilon 2$) to enable hydrogen-bond stabilization of the oxyanion. The motif incorporated fixed atomic coordinates from His159 (C$\beta$ atom and all atoms of the imidazole ring); Asn175 (C$\beta$, C$\delta$, $N\delta 1$, and $O\delta 2$ atoms), Gln19 ($N_{\varepsilon 2}$, $O\varepsilon 1$, C$\delta$, and C$\gamma$ atoms) atoms; all heavy atoms of Cys25; and the complete substrate. Additionally, we fixed backbone $N$, $C\alpha$, and $C$ atoms of Ser24 and the $N$, $C\alpha$, $C$, $O$ atoms of Trp26 to position the nucleophile at the N-terminus of a helix, enabling oxyanion stabilization via the helix dipole. The motif is shown in Figure 16E.

For scaffold generation, we provided the motif as unindexed atomic motif conditioning to RFdiffusion2 to generate scaffolds for protein lengths drawn uniformly from the range of 95–185 amino acids. An earlier version of the model—fine-tuned from structure prediction weights and trained with a motif-centering approach (appendix B.3)—was employed for this campaign. In the current version of the model, one can specify the approximate motif:protein center of mass offset appendix B.3), but this was not possible in the motif-centered version, so as a workaround we recentered $B^{(t)}$ to the origin for the first $t'$ timesteps to ensure burial of the motif. For this design campaign, we used $t'$ values of 1, 3, 5, and 10 over a total of 200 denoising steps and generated 10,000 scaffolds for each $t'$. Designability of the scaffolds was evaluated by generating eight sequences per scaffold using LigandMPNN, predicting their structures with AlphaFold2, and identifying scaffolds where at least one sequence achieved global $C\alpha$ RMSD $< 1.5$Å and motif residue heavy-atom RMSD $< 1.5$Å. This assessment yielded approximately 1,200 designable scaffolds.

In stage-1 sequence design, we designed 100 sequences per designable backbone using three iterative cycles of LigandMPNN [32] and Rosetta FastRelax [59] with geometric constraints on the catalytic residues and the orientation of the ligand relative to the nucleophile. This approach was used to allow for broader exploration of sequence space under minor backbone refinements and substrate conformer changes, while maintaining the catalytic geometry defined in the initial motif (this is illustrated in fig. 16E for the design *Eta1*). The constraints defined precise geometric parameters and tolerances (0.1Å for distances, 5° for angles and dihedrals) for all hydrogen bonding networks in the active site (Cys–His, His–Substrate, the backbone and side-chain oxyanion interactions, as well as the His–Asn interaction). Subsequent filtering with AlphaFold2 included stringent structural criteria (global $C\alpha$ RMSD $< 1.5$Å, catalytic residue $C_\alpha$ RMSD $< 1.0$Å upon global $C\alpha$ alignment, and motif residue heavy-atom RMSD $< 1$Å), yielding 1,321 designs derived from 331 distinct scaffolds.

In stage-2 sequence design, we employed PLACER (Protein-Ligand Atomistic Conformational Ensemble Resolver) [39] to quantitatively assess active site preorganization as described by Lauko et al. [1]. PLACER is a graph neural network that was trained to recapitulate corrupted atomic coordinates of side chains and small molecules within a spherical region of up to 600 heavy atoms. For each design, we generated 50 structural ensembles for both the enzyme–substrate complex and acyl–enzyme intermediate states using the AlphaFold2 predictions as input. For the apo prediction, we placed the ligand from the design model obtained in the constrained sequence design step by superimposing on the $C\alpha$ atoms of the AlphaFold2 prediction. The acyl–enzyme intermediate was modeled as a modified Cysteine residue (CCD code: `CY4`). Our filtering criteria incorporated PLACER's predicted uncertainty in atomic position and distance measurements across these ensembles. Specifically, median Cys-$S\gamma$:His-$N\delta 1$ distance ($< 3.8$Å) and accompanying median uncertainties for both atoms ($< 1.5$Å), and median $N\varepsilon 2$–$O\delta 1$ distance ($< 3.3$Å) filters were used. For the acyl–enzyme intermediate predictions, we also filtered on median distances between the carbonyl oxygen and both the Cys backbone $N$ ($< 3.3$Å) and Gln-$N\varepsilon 2$ ($< 3.5$Å). To enrich our candidate pool, we used the 57 parent scaffolds from the sequences that passed the first two PLACER filters to further sample 1000 sequences for each scaffold using the constrained sequence design method and AlphaFold2 structural criteria described above. From this unified candidate pool, we selected 48 designs that passed all the PLACER filters while maintaining sequence diversity ($< 70\%$ sequence identity between designs from the same scaffold). This resulted in 48 designs representing 29 distinct scaffolds for experimental characterization, with 2 sequences originating from the stage-1 sequence design and the remainder from stage-2.

##### D.2.2 Experimental Methods

***Protein Expression and Purification***

Genes encoding the designed proteins were synthesized by Integrated DNA Technologies (IDT) as eBlocks and subsequently cloned into the expression vector LM627 (addgene) containing the C-terminal SNAC tag and a hexahistidine-tag via the

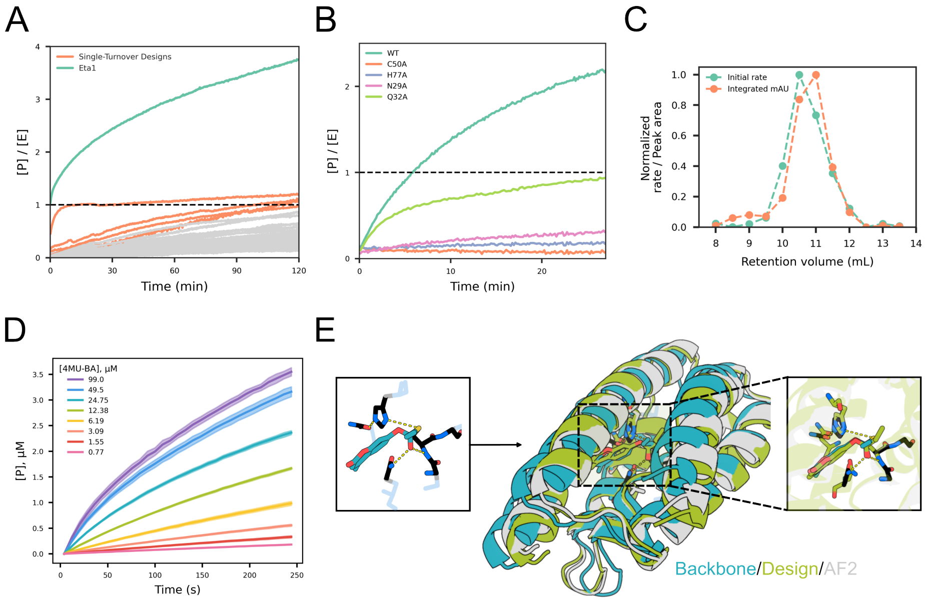

**Figure 16**: **Cysteine hydrolase design and characterization**: (A) Reaction progress curves from the initial activity screen showing the normalized activity (background subtracted) of the designed enzymes. Dashed line indicates the single-turnover end-point where [P] = [E]. In gray are designs that fail to reach this end-point over the course of the two hour experiment. (B) Reaction progress curves showing the normalized activity (background-subtracted) of the wild-type (green) and knockout variants of *Eta1* where each active-site residue is individually mutated to Alanine. Dashed line indicates the single-turnover end-point. (C) Overlay of the activity profile (initial rate, in green) and the corresponding absorbance peak area (orange) of each SEC fraction obtained from re-purifying the *Eta1* sample, normalized by the maximum value. (D) Reaction progress curves (background-subtracted) of 3 $\mu$M of *Eta1* incubated with different substrate concentrations in to determine Michaelis Menten parameters. Shaded areas represent standard deviation of three technical replicates. (D) Overlay of the RFdiffusion2 generated backbone (teal), the design model obtained after LigandMPNN and Rosetta Fast Relax (green), and the AlphaFold2 prediction (gray) for *Eta1*. The input motif is shown in the box to the left and is superimposed on the active-site from the design model in the box to the right, with the dashed yellow lines indicating the hydrogen-bonding network in the active site. Rosetta Fast Relax makes minor backbone refinement and substrate conformer changes while constraining the active-site geometry defined in the motif. AlphaFold2 prediction for *Eta1* shows excellent agreement with the design model (global $C\alpha$ RMSD = 0.57Å).

Golden Gate assembly method, following established protocols outlined in Wicky et al. [60]. The resulting recombinant plasmids were transformed into *Escherichia coli* BL21(DE3) competent cells using heat-shock, with cells recovered in 100 $\mu$L of SOC medium at 37 °C for 1 hour. After recovery, the transformed cells were aseptically transferred to individual wells of a 2mL deep-well plate, each containing 900 $\mu$L of LB medium supplemented with 50 $\mu$g/mL kanamycin. The plate was sealed with a breathable film and incubated overnight at 37 °C with shaking at 1200 rpm. For each design that exhibited growth—as indicated by turbidity— glycerol stocks were made and 500 $\mu$L of the overnight culture was transferred into a 250mL flask containing 50 mL of autoinduction medium [61]. The cultures were incubated at 37 °C with shaking for 4–6 hours to promote cell growth, after which the temperature was reduced to 18 °C for overnight protein expression.

After expression, cultures were harvested by centrifugation at 4000 $g$ for 15 minutes at 4 °C, and the supernatant was discarded. The cell pellets were resuspended in 25 mL of ice-cold wash buffer (40 mM imidazole, 500 mM NaCl, 50 mM sodium phosphate, pH 7.4, and 1 mM TCEP), supplemented with 1 mg/mL lysozyme and 0.1 mg/mL DNase I to facilitate cell lysis and reduce viscosity. The suspensions were sonicated on ice for 2.5 minutes at 80% amplitude using 10-second on/off cycles. The lysates were then clarified by centrifugation at 14 000 $g$ for 30 minutes at 4 °C, and the supernatants containing soluble proteins were collected. Protein purification was performed via Ni-NTA affinity chromatography. Each supernatant was applied to 1 mL of Ni-NTA resin pre-equilibrated with the wash buffer, washed three times with 15 mL of the same buffer to remove contaminants, and subjected to a pre-elution with 400 $\mu$L of elution buffer (400 mM imidazole, 500 mM NaCl, 50 mM sodium phosphate, 1 mM TCEP, pH 7.4) prior to elution with 1.3 mL of the same buffer. The eluates were further purified by size-exclusion chromatography on a Superdex 75 Increase 10/300 GL column (GE Healthcare) using a running buffer composed of 20 mM HEPES, 50 mM NaCl, and 1 mM TCEP (pH 7.4). The purified protein fractions were collected and stored at 4 °C until further use.

###### ***Kinetic Analysis and Hit Validation***

To evaluate the enzymatic activity of the designed proteins, kinetic assays were performed using the fluorogenic substrate 4-methylumbelliferyl butyrate (4MU-Bu). Upon hydrolysis, 4MU-Bu releases 4-methylumbelliferone (4MU), a fluorescent compound that permits real-time monitoring of the reaction progress. In the initial activity screen, reactions were set up in a 96-well half-area microtiter plate with a total volume of 40 $\mu$L per well and a final substrate concentration of 100 $\mu$M. Both the purified protein samples, obtained from the size-exclusion chromatography (SEC) fractions, and the substrate solutions were freshly prepared in an assay buffer comprising 20 mM HEPES, 50 mM NaCl (pH 7.4) and 5% dimethyl sulfoxide (DMSO) to enhance substrate solubility. For each reaction, 4 $\mu$L of a 10-fold diluted enzyme sample was dispensed into the well, and the reaction was initiated by the addition of 36 $\mu$L of the substrate solution. The formation of 4MU was continuously monitored at 30 °C using a Neo2 plate reader (Biotek) with excitation and emission wavelengths set to 365 nm and 445 nm, respectively. A negative control containing only the reaction buffer was also run concurrently, and its fluorescence signal was subtracted from the enzyme reactions to correct for background fluorescence. Standard curves were generated by measuring the fluorescence of known 4MU concentrations, enabling conversion of fluorescence units to product concentration. Enzyme concentrations in the reaction mixtures ranged between 1–7 $\mu$M. The resulting normalized activity profiles ($[P]/[E]$ versus time) are presented in fig. 16A. Several designs exhibited detectable fluorescence, indicating substrate cleavage. Of the 48 designs, four reached the single-turnover endpoint, as indicated by the final product concentration approximating the enzyme concentration. The most active design, named *Eta1*, demonstrated multiple turnover activity. *Eta1* displayed a burst-phase kinetic profile—indicative of a mechanism where the nucleophilic attack by water is rate-limiting [45]. Based on these results, *Eta1* was selected for further characterization to confirm its activity and determine its Michaelis–Menten parameters, while the other designs were not further characterized.

First, the molecular weight of *Eta1* was confirmed by liquid chromatography-mass spectrometry (LC/MS) (results shown in Table X). To verify that the observed catalytic activity was inherent to our designed enzyme and not due to an active contaminant, we re-purified the sample by size-exclusion chromatography (SEC). Individual SEC fractions were collected and assayed for activity using the same fluorometric protocol described above. As shown in fig. 16C, the normalized activity profile—overlaid with the normalized $A_{280}$ absorbance—matched the enzyme's elution peak, indicating that the activity is attributable to the intended protein. To further substantiate that the catalytic function originates from the designed active site residues, we tested knockout variants in which each active site residue was individually replaced by alanine. As shown in fig. 16B, all knockout variants exhibited markedly reduced catalytic activity compared to the wild-type design, with the glutamine (Gln19) knockout showing reduced, single-turnover activity, likely due to impaired oxyanion stabilization. Note that the activity of the wild-type *Eta1* was lower compared to what was observed during the initial activity screen, likely due to partial oxidation of Cysteines during extended storage of this sample at 4 °C.

To determine the kinetic parameters of *Eta1*, a fresh batch of samples was expressed and purified from the glycerol stocks using the same protocol outlined above. To preserve stability, the collected SEC fractions were aliquoted, immediately snap-frozen in liquid nitrogen and stored at −80 °C until further use. Initial reaction velocities were measured at a fixed enzyme concentration of 3 $\mu$M in a 100 $\mu$L reaction volume using an assay buffer comprising 20 mM HEPES and 50 mM NaCl (pH 8.0). Reaction rates were obtained across a series of substrate concentrations, with each condition performed in triplicate. In parallel, a calibration curve was prepared under these reaction conditions to convert fluorescence signals into product concentrations. These data were then used to calculate the second-order rate constant ($k_2/K_M$) for the acylation step, providing a measure of catalytic efficiency (reaction progress curves are shown in fig. 16D and the Michaelis-Menten plot in fig. 4B). Notably, *Eta1* exhibited a catalytic efficiency of 250 $M^{-1}\,s^{-1}$, which is significantly higher than the best values reported in prior work on *de-novo* designed Cysteine Hydrolases [45] for the same Coumarin leaving group (excluding evolved variants), highlighting the significance of RFdiffusion2 in scaffolding complex active sites required for multi-step catalysis.

#### D.3 Zinc Hydrolase

##### D.3.1 Design

Zinc metallohydrolases were designed for the fluorogenic esters 4-methylumbelliferyl phenylacetate (4MU-PA) and 4-methylumbelliferyl butyrate (4MU-B) as model reactions that would allow for kinetic characterization of active enzymes and benchmarking with previously designed zinc esterases. The design pipeline and results for 4MU-PA are described extensively in a separate publication [50]. A similar design strategy was pursued for 4MU-B. Note that a second design campaign was performed for the 4MU-PA substrate in [50], described further in that manuscript, with the underlying pipeline remaining the same.

Briefly, rational theozymes for the rate-limiting hydroxide attack transition state in the proposed hydrolysis reaction were constructed and optimized using density functional theory. Each theozyme contained three histidines coordinated to the zinc ion and a carboxylic acid to represent a protonated general base that deprotonated the attacking water molecule. The oxyanion in each theozyme was stabilized either by the zinc ion or two methanol molecules representing general hydrogen bonds coming from a protein sidechain or backbone (see [50] for more details). Approximately $4500 \sim 5000$ enzyme scaffolds were generated in total around the designed theozymes for each substrate using RFdiffusion2 with the catalytic histidine imidazoles fixed in place during generation. The general base and oxyanion stabilizing hydrogen bonds were left to be found during later sequence generation to explore additional backbone and sequence space alike due to the flexibility of residues that can accomplish these tasks. About 1000 competent scaffolds were resampled with RFjoint2 [62] inpainting to modify loop structures and lengths outside of the active site to generate extra diversity for each scaffold. ProteinMPNN was used to generate 20 global sequences for each scaffold while keeping the catalytic histidines fixed. All sequences were then passed through AlphaFold2. Sequences whose AlphaFold2 predicted structures matched the designed structure were kept. Successfully folded designs then underwent iterative active site sequence redesign and constrained fast relaxing (i.e., energy minimization) with LigandMPNN and Rosetta [63], respectively (see [50] for more details). The catalytic histidines remained fixed throughout design. General bases that were found within 3Å of the theozyme hydroxide position and residues identified to Hbond with the substrate were kept. This step produced several hundred thousand resulting design sequences across hundreds of unique scaffolds, but only the designs containing a general base, and hydrogen bonds to the oxyanion hole for the theozymes modeled with them, were kept. These updated design sequences were then passed through AlphaFold2 to examine the predicted structure confidence and agreement with the designed structure. Structurally verified designs were then predicted and scored by ChemNet [39] for substrate and catalytic side chain preorganization and agreement with the designed active sites. Rosetta was used to further score the designs on additional metrics. Designs that had the best balance of favorable ChemNet and Rosetta metrics within a scaffold family were selected for a 96-well order. That is, the top-scored sequence was selected within 96 different scaffolds.

***Theozyme construction***

The initial theozymes modeling the rate-limiting nucleophilic zinc-hydroxide attack of the 4MU-PA or 4MU-B ester were constructed manually in a 3D chemical modeling software. Each theozyme consisted of (1) a single Zn(II) ion, (2) three imidazole rings representing histidine sidechains coordinating to the zinc ion, (3) the 4MU-PA or 4MU-B ester substrate, (4) a hydroxide ion coordinated to the zinc ion representing the deprotonated nucleophilic water molecule, and (5) an acetic acid group interacting with the hydroxyl group, representing a protonated catalytic general base (e.g., aspartate or glutamate). Furthermore, we considered two modes for oxyanion stabilization: (1) Zn(II)-O interaction, and (2) external hydrogen bond oxyanion hole. The Zn(II)-O oxyanion stabilization uses the zinc ion to stabilize the forming oxyanion in addition to stabilizing the nucleophilic hydroxide, requiring no extra elements to be modeled. On the contrary, the external hydrogen bond oxyanion stabilization has the oxyanion facing away from the zinc and uses two additional H-bonded methanol molecules to mimic a potential oxyanion hole. These two methanol molecules represent a generic form of an oxyanion hole and help locate the transition state geometry more akin to what it would look like in the presence of an oxyanion hole by polar groups in an enzyme active site. Each of these two classes of theozymes had the nucleophilic attack modeled from each side of the planar 4MU-PA/4MU-B ester bond before forming the chiral tetrahedral intermediate (i.e., R or S attack). Thus, four unique theozymes were modeled in total as transition states for each substrate (i.e., 8 total DFT theozymes).

All transition state optimization calculations were performed using Gaussian 16 software [64]. Structural optimizations and frequency calculations were performed with the B3LYP-D3 method along with the 6-31G(d) basis set and the SDD ECP on the Zn(II) atom. D3 dispersion correction was applied using the Becke-Johnson damping function [65]. Solvent effects of water were included using the CPCM solvation model during geometry optimization. Frequency calculations were performed to confirm whether the structure is a minimum or a transition state. Intrinsic reaction coordinate (IRC) analysis was used to confirm that the obtained transition states connect the correct minima.

Conformational flexibility of the transition states was sampled based on rotatable bonds in the 4MU-PA and 4MU-B substrates. The conformers were sampled as frozen coordinates with no geometry optimization, in order not to affect the transition state geometries. Energies of the conformers were evaluated using the GFN2-XTB semiempirical QM method [66]. This procedure was performed using a Python script frozen-conf-xtb, available on GitHub[4]. We generated 10 conformers for each transition state, all within 5 kcal/mol of each other, and with a minimum difference between conformers being at least 20 degrees for at least one rotatable bond. These conformers do not necessarily represent local minima in the conformational energy surface but rather capture structural diversity among low energy conformers. Depending on the specific protein environment each of these conformers may be more or less preferred, and furthermore, Rosetta minimization is capable of adjusting these rotatable bonds within a small range.

[4] https://github.com/ikalvet/frozen-conf-xtb

We generated unique Rosetta ligand params files from each of the transition state geometries. Each params file defines the ligand, containing the 4MU-PA or 4MU-B substrate atoms, the Zn(II) atom, and the hydroxyl group. Rosetta ligand params file was created from the produced conformers by first converting the conformer multiXYZ files to mol2 files using Open Babel, and thereafter converting the mol2 file to params using the `$ROSETTA_MAIN\source\scripts\python\public\molfile_to_params.py` script provided in Rosetta.

##### D.3.2 Materials

###### *General solvents and buffers*

Milli-Q water was obtained in-house using a Q-POD® Ultrapure Water Remote Dispenser from Millipore Sigma. Nuclease-free water was commercially bought from Invitrogen™.

Dimethylsulfoxide (DMSO) was obtained from Sigma Aldrich. All general buffer components (i.e., HEPES, NaCl, Tris-HCl, EDTA) were bought from Millipore Sigma. All buffers were prepared in Milli-Q water and filtered through a 0.2 $\mu$m pore size Nalgene™ Rapid-Flow™ Sterile Disposable Filter Unit with a PES membrane. Further mentions of 'bottle filtration' will refer to this 0.2 $\mu$m pore size filter.

###### *Materials for biology*

Synthetic linear DNA fragments encoding the genes for each reverse-translated protein design with overhangs for golden gate assembly were ordered in a 96-well eblock® from IDT, shipped dry, and stored at -20˚C. Custom plasmid vectors were ordered from GenScript and stored at -20˚C. Golden gate assembly of the linear gene fragments into the custom plasmid vectors was done using the NEBridge® Golden Gate Assembly Kit (BsaI-HF® v2) from New England Biolabs. Competent BL21(DE3) E. coli cells for protein expression and competent NEB 5-alpha E. coli cells for plasmid production were commercially bought from New England Biolabs and stored at -80˚C. Super optimal broth with catabolite repression (SOC) media was purchased from New England Biolabs for recovery after heat-shock transformation.

Lysogeny broth (LB) media for E. coli inoculation was made from tryptone (Sigma Aldrich; 10 g/L), NaCl (Millipore Sigma; 10 g/L), and Bacto® Yeast Extract (ThermoFisher Scientific; 5 g/L) dissolved in Milli-Q water. Terrific broth II (TB-II) media for protein production was bought as a solid powder from MP Biomedicals and dissolved at a concentration of 50 g/L in Milli-Q water. All media were prepared under flame, sterilized through bottle filtration, and immediately used or stored at 4˚C before use. TB-II and LB were either autoclaved or sterilized by bottle filtration before use. Kanamycin sulfate from Streptomyces kanamyceticus was bought as a powdered bioreagent from Millipore Sigma. Isopropyl $\beta$-D-1-thiogalactopyranoside (IPTG, dioxane-free) to induce protein expression was bought from Fisher Scientific. Agar plates with kanamycin were made in-house for colony isolation. Glycerol ($\geq$99.0% grade) for glycerol stocks was bought from Sigma Aldrich.

Axygen® 96-well clear round bottom 2 mL polypropylene deep well plates for cultures are bought from Corning (P-DW-20-C-S). Breathe-Easier® breathable films for culture plates were bought from Andwin Scientific. Axygen® aluminum sealing films for general sealing purposes were bought from Corning (PCR-AS-600).

***Materials for protein purification***

Lysozyme, DNase I, and Polymyxin B sulfate used in E. coli cell lysis were bought from Sigma Aldrich (L6876, DN255G, P0972), respectively. Pierce® protease inhibitor tablets were bought from Thermo Scientific™. Filter polypropylene microplates with 96 wells (800 $\mu$L/well), long drip spouts, and 25 $\mu$m polyethylene frit for small-scale affinity chromatography were purchased from Agilent. Econo-Pac® 25 mL gravity flow-through chromatography columns for low-throughput affinity chromatography were purchased from BioRad. Strep-Tactin® resin for affinity chromatography was purchased from IBA Lifesciences. Desthiobiotin used in protein elution was purchased from IBA Lifesciences. Strep-Tactin® regeneration buffer containing HABA (10x concentration) was also purchased from IBA Lifesciences. Amicon® 3 kDa and 10 kDa MWCO centrifugal concentrators for protein spin-concentration were purchased from Millipore Sigma. Size exclusion chromatography was performed on an ÄKTA pure™ chromatography system from Cytiva using a Superdex 75 Increase 10/300 GL column also purchased from Cytiva.

***Materials for enzyme characterization***

4-Methylumbelliferyl phenylacetate (4MU-PA, commercially known as phenyl-acetic acid 4-methyl-2-oxo-2H-chromen-7-yl ester) for the first round of activity screening was acquired from Millipore Sigma. 4MU-PA for the second round of activity screening and all kinetics characterization experiments was custom synthesized by WuXi. 4-Methylumbelliferyl butyrate (4MU-B) for all experiments was purchased from Millipore Sigma. 4-Methylumbelliferone (4MU) for making product concentration standards was purchased as a solid from Millipore Sigma. Zinc sulfate was purchased from Sigma Aldrich. Full area 96-well flat bottom assay plates with a non-binding surface made from black polystyrene, used in screening measurements, were purchased from Corning. Half area 96-well flat bottom assay plates with a non-binding surface made from black polystyrene, used in kinetic measurements, were purchased from Corning. Fluorescence measurements were performed with The BioTek Synergy Neo2 Hybrid Multimode Reader RUO.

##### D.3.3 Experimental Methods

***Cloning and transformation***

Genes encoding the designed proteins with overhangs for Golden Gate assembly were ordered from IDT as dry eBlocks™. The DNA sequences were codon-optimized for increased E. coli expression, and ordered as follows:

$$(5')\texttt{ATACTACGGTCTCAAGGA} - \texttt{design} > -\texttt{GGTTCCCGAGACCGTAATGC}(3')$$

The DNA fragments were cloned into a custom vector (pDTstrep1; 5684 base pairs) encoding a kanamycin-resistance gene for positive selection, a T7 promoter, and a

C-terminal Strep-tag®, following a published protocol[60]. The final expressed proteins were obtained as MSG-design-GSA-WSHPQFEK, where WSHPQFEK is the C-terminal Strep-tag® sequence for Strep-Tactin® affinity chromatography purification.

Before golden gate assembly, 20 $\mu$L of nuclease-free water was added to each well of the dry eBlock before the plate was covered with an aluminum seal, vortexed, and centrifuged (1000g x 30s) to suspend the linear DNA fragments at a concentration of 10 ng/$\mu$L (approximately 23 nM for the largest fragments of 700 bp). Golden gate assembly was performed using the NEBridge® Golden Gate Assembly Kit (BsaI-HF® v2) containing (1) a solution of BsaI type IIS restriction enzyme and T4 DNA ligase, and (2) a T4 ligase buffer with 1 mM ATP. The reactions were set up at a 5 $\mu$L scale such that DNA fragment concentration was at least 8.8 nM for the largest DNA fragments (700 bp) and the cloning vector concentration was 4.4 nM. First, a master mix of the cloning reaction components (GG assembly kit, T4 buffer, cloning vector, and ddH2O) was prepared. For each 5 $\mu$L reaction, the master mix consisted of 1.8 $\mu$L ddH2O, 0.5 $\mu$L T4 buffer, 0.25 $\mu$L GG assembly kit, and 0.54 $\mu$L cloning vector (from 41 nM stock), mixed in the given order. The master mix was prepared for 96 reactions in a cooled microcentrifuge tube and then spread out in 3.1 $\mu$L aliquots to a cooled PCR plate. Lastly, 1.9 $\mu$L solution of the DNA fragment ($\sim$23 nM stock in case of 700 bp fragment) was added to each well and carefully mixed with the pipette tip. The plate was sealed, and incubated in a thermocycler (37°C) for 20 minutes, followed by a 5 minute incubation at 60°C to heat-inactivate the enzymes. The resulting plasmid products of Golden Gate assembly were then used for transformation; excess reaction mixture was stored at -20°C for future use.

Bacterial transformation was performed over ice using competent BL21(DE3) E. coli cells. A new PCR plate was set into a cooled metal block on ice and 2.0 $\mu$L of the Golden Gate plasmid products from the previous steps were transferred into their respective wells with a multichannel pipette. This PCR plate was briefly sealed and centrifuged (1000g x 30s) to get the plasmids at the bottom of each well. Next, tubes of competent E. coli stored at -80°C were allowed to thaw and swiftly added in 7 $\mu$L aliquots to each well with a repeater pipette. The PCR plate was rapidly centrifuged (100g x 5-10s) to ensure sufficient mixing of the cells and plasmids at the bottom of each well and the plate was returned to the metal cooling block on ice for 30 minutes. Thereafter, the plate was transferred to a 42°C metal block for 17 $\sim$ 20s to heat shock the E. coli and returned to ice for 2 minutes. 100 $\mu$L of pre-warmed SOC recovery media was added into each well of the PCR plate and a breathable film was used to cover the plate. The plate was incubated in a microplate shaker (37°C, 1200 rpm) for 1 hour. 900 $\mu$L aliquots of sterile LB media with kanamycin (50 $\mu$g/mL) were added to each well in a 2 mL deep round-well incubation plate under flame, followed by 100 $\mu$L of the transformed E. coli in SOC media. The deep round-well plate was covered with a breathable film and incubated in a microplate shaker (37°C, 1200 rpm) for 16-18 hours.

Successfully cloned designs had opaque wells while unsuccessfully cloned designs had transparent wells the following day. Glycerol stocks were made for every clone by mixing 125 $\mu$L of 50% v/v glycerol in Milli-Q water with 125 $\mu$L of the overnight culture in a new PCR plate. The PCR plate was covered with an aluminum seal and stored at -80°C. Note that the obtained clones are not sequence-verified at this stage and could be polyclonal or even wrong due to mistakes during DNA synthesis. After preparing the glycerol stocks, the remaining cultures were used for expression, as described below.

##### *Small-scale protein expression with IPTG*

Proteins were expressed at a 4x1 mL culture scale in sterile terrific broth II (TB-II) media supplemented with 50 μg/mL of kanamycin. Four 2 mL deep round-well plates were loaded with 1 mL of the TB-II+kanamycin media under flame. 150 μL of the overnight cultures in LB media were transferred into each respective well for each of the four plates. These plates were covered with breathable film and incubated in a microplate shaker (37°C, 1200 rpm) for 90 minutes to promote sufficient cell growth. Thereafter, 11.5 μL of 100 mM IPTG (isopropyl $\beta$-D-1-thiogalactopyranoside) was added into every well of the four plates (1 mM final [IPTG]) under flame to induce expression. The plates were resealed with the breathable films and incubated in the microplate shaker (37°C, 1200 rpm) for 2 hours before being transferred to a room temperature microplate shaker (25°C, 1200 rpm) to incubate for an additional 20-24 hours. The following day, the plates were centrifuged (4,198 g, 8 min, 25°C) to collect the cell pellets. The resulting plates containing the cell pellets were covered with an aluminum seal and stored at -80°C until purification.

##### *Protein purification for screening*

The frozen cell pellets were thawed and lysed, and the expressed protein was purified by affinity chromatography with Strep-Tactin® resin following an adapted published procedure [67]. First, three buffers were prepared with Milli-Q water as follows: **W1 buffer** (100 mM Tris-HCl, 150 mM NaCl, 1 mM EDTA, pH 8.0), **W2 buffer** (100 mM Tris-HCl, 150 mM NaCl, pH 8.0), and **E buffer** (40 mM HEPES, 50 mM NaCl, 2.5 mM desthiobiotin, pH 8.0). Additionally, a **lysis buffer** was made from the W2 buffer where it was supplemented with 1 mg/mL lysozyme, 0.5 mg/mL polymyxin B sulfate, 10 μg/mL of DNAse I, and 1 Pierce® protease inhibitor tablet per 50 mL of W2 buffer. All buffers were stirred until the reagents were completely dissolved, and sterile-filtered through a 0.2 μm pore size bottle-filter.

Four plates of cell pellets (i.e., 4 mL of culture for each design) were simultaneously thawed and 600 μL of **lysis buffer** was added to each well in one plate and transferred to the other three plates to combine all of the same-design pelleted cultures in the fourth plate (e.g., A1 plate 1 → A1 plate 2 → A1 plate 3 → A1 plate 4). Cells were suspended by repeatedly pipetting the suspension in and out of the tip. The final plate containing the combined cultures in lysis buffer was covered with an aluminum seal and placed in a shaking microplate incubator at 37°C, 1200 rpm for 60 minutes and then transferred to a 60°C metal bead bath for 20 minutes. The plate was then centrifuged (4,198 g, x 30 minutes; 14°C) to pellet the insoluble lysate aggregates before column purification.

Affinity chromatography was performed by adding 200 μL of a 50% Strep-Tactin® resin suspension to each well in a 96-well 25 μm fritted plate such that 1 CV = 100 μL of resin. The resin was equilibrated with 500 μL of W1 buffer 3 times (3 x 5 CV of W1 buffer) while using a vacuum manifold to pull the liquid through the plate. The bottom spouts of the fritted plate were sealed with parafilm, and 600 μL of the soluble lysate supernatant was gently added to each well in the fritted plate. The fritted plate was covered with an aluminum seal and inverted several times to resuspend the resin. The resuspended resin was allowed to settle at room temperature for 15 minutes before the parafilm and aluminum seal were removed. The protein-loaded resin was washed 2 times with 500 μL of W1 buffer (2 x 5 CV of W1 buffer) and 1 time with 500 μL of EDTA-free W2 buffer (1 x 5 CV of W2 buffer) while using a vacuum manifold to pull the liquid through. Finally, the bound protein in each column was eluted into a 2 mL deep 96 well plate by adding 200-300 μL of E buffer (1 x 2-3 CV of E buffer) and eluting by centrifugation (1,500 g, 2 min). The eluted protein plate was covered with an aluminum seal and stored at 4°C until further use.

The used Strep-Tactin® resin was recovered by first washing each well in the plate with 100-200 μL of E buffer (1 x 1-2 CV of E buffer), 500 μL of W1 buffer (1 x 5 CV of W1 buffer), and 300 μL of W2 buffer (1 x 3 CV of W2 buffer). Finally, the resin was regenerated by adding 3x500 μL of regeneration buffer (purchased as 10x concentrated and diluted in Milli-Q water) (3 x 15 CV of regeneration buffer) until a dark red color was homogeneous through the resin. Note that the resin was resuspended several times via pipetting during the cleaning and regeneration steps to remove aggregates and ensure homogeneity. The regenerated resin wells in the fritted plate were stored under 40-60% v/v ethanol and water to prevent bacterial growth; the plate was covered with an aluminum seal on the top and parafilm on the bottom spouts before being stored at 4°C until further use.

***Screening fluorescence assay for 4MU-B***

The activity screening assay for the 4MU-B designs was performed in a 96-well (full area) fluorescence plate to screen for the hydrolysis of the fluorogenic ester 4MU-PA (Sigma), monitored by a plate-reader using an excitation/emission wavelength of 365 nm and 445 nm, respectively. Briefly, each reaction was run at 30°C using 6 μL of eluted enzyme containing 200 μM of supplemented zinc sulfate and 54 μL of a substrate solution composed of 100 μM of 4MU-B in reaction buffer (40 mM HEPES, 50 mM NaCl, pH 8.0) with 5.0% v/v DMSO. Thus, the final reaction in each well contained the eluted enzyme (1/10th the eluted concentration), 20 μM of zinc sulfate, 90 μM of 4MU-B, and approximately 4.5% v/v DMSO in the reaction buffer. The reaction plate was placed into the plate reader, shaken for 3s at 282 cpm, and fluorescent progress curves monitored for one hour. Note that the enzyme concentration was not standardized in the screening assay, as this was only used to identify hits to characterize in more detail.

***Screening fluorescence assay for 4MU-PA (1st round of 96 designs)***

The activity screening assay for the first round of designs was performed in a 96-well (full area) fluorescence plate to screen for the hydrolysis of the fluorogenic ester 4MU-PA (Sigma), monitored by a plate-reader using an excitation/emission wavelength of 365 nm and 445 nm, respectively. Briefly, each reaction was run at 30°C using 6μL of eluted enzyme containing 200 μM of supplemented zinc sulfate and 54 μL of a substrate solution composed of 10 $\mu$M of 4MU-PA in reaction buffer (40 mM HEPES, 50 mM NaCl, pH 8.0) with 5.0% v/v DMSO. Thus, the final reaction in each well contained the eluted enzyme (1/10th the eluted concentration), 20 $\mu$M of zinc sulfate, 9 $\mu$M of 4MU-PA, and approximately 4.5% v/v DMSO in the reaction buffer. The reaction plate was placed into the plate reader, shaken for 3s at 282 cpm, and fluorescent progress curves monitored for one hour. Note that the enzyme concentration was not standardized in the screening assay, as this was only used to identify hits to characterize in more detail.

***Screening fluorescence assay for 4MU-PA (2nd round of 96 designs)***

The activity screening assay for the second round of designs was performed was performed in a 96 half-well fluorescence plate to screen for the hydrolysis of 4MU-PA (WuXi), monitored by a plate-reader using an excitation/emission wavelength of 365 nm and 445 nm, respectively. Briefly, each reaction was run at 25°C using 10 $\mu$L of eluted enzyme containing 100 µM of supplemented zinc sulfate and 90 µL of a substrate solution composed of 111 $\mu$M of 4MU-PA in reaction buffer (40 mM HEPES, 50 mM NaCl, pH 8.0) with 5.55% v/v DMSO. Thus, the final reaction in each well contained the eluted enzyme (1/10th the eluted concentration), 20 $\mu$M of zinc sulfate, 100 $\mu$M of 4MU-PA, and approximately 5% v/v DMSO in the reaction buffer. The reaction plate was placed into the plate reader, shaken for 3s at 282 cpm, and fluorescent progress curves monitored for one hour. Note that the enzyme concentration was not standardized in the screening assay, as this was only used to identify hits to characterize in more detail.

***Isolation and sequence-verification of colonies***

Identified screening hits and designs of interest were characterized in greater detail and verified through DNA sequencing and electrospray ionization mass spectrometry (ESI-MS; mass spectrometry methods below). Single clones of identified hits were obtained by the following procedure. A stab from the corresponding polyclonal glycerol stock was diluted into 200 $\mu$L of sterile water, and 2 $\mu$L of this was further diluted into another 200 $\mu$L of water. 20 $\mu$L of the diluted cell suspension was spread on an LB-agar plate supplemented with kanamycin, and grown at 37°C overnight. The following day, 1-3 of the colonies were picked by careful swabbing with a pipette tip and transferred to individual PCR tubes containing 100 $\mu$L Milli-Q water. Then, 94 $\mu$L of each isolated colony in water was used to inoculate a 1 mL LB culture containing 50 $\mu$g/mL of kanamycin for overnight growth and the rest of the isolated colony in water was used for DNA sequencing (described below). The overnight LB+kanamycin cultures were incubated at 37°C and 225 rpm until the following day when they were visibly opaque (~16-20 hours of growth). Glycerol stocks were made for each of the isolated clones that had verified DNA sequences containing the gene of interest. In the case of a single design having 2+ isolated clones that were DNA-sequence verified, one was chosen at random for glycerol stocks and future experiments. Note that all of these steps were performed using sterile technique under flame.

The DNA sequencing of isolated colonies was performed using Sanger sequencing. ColonyPCR with T7 and T7-term primers was performed to amplify the expressing region of the DNA contained in the isolated cells. The PCR reaction mixture comprised

2 $\mu$L of the single colony suspended in nuclease-free water, 1 $\mu$L of 10 $\mu$M T7 and T7 terminal primers designed to flank the target sequence, 12.5 $\mu$L of GoTaq Green 2X Master Mix (which includes Taq DNA polymerase and the accompanying buffer), and 8.5 $\mu$L of additional nuclease-free water. The thermal cycling conditions were as follows: an initial lysis and denaturation step at 95°C for 3 minutes, followed by 31 cycles of denaturation (95°C, 30 s), annealing (58°C, 30 s), and extension (72°C, 1 min), culminating in a final extension step at 72°C for 5 minutes. The success of the PCR reactions was verified by DNA gel electrophoresis using 1.2% agar/TAE gel with SybrSafe dye. Subsequently, 10$\mu$L of unpurified PCR products were sent for sequencing at Genewiz/Azenta, and the results were compared against the design sequences.

##### *Large-scale protein expression with IPTG*

DNA sequence-verified, single colony glycerol stocks were used to inoculate 5 mL overnight cultures in LB media with 50 $\mu$g/mL of kanamycin that were incubated at 37°C and 225 rpm. After 16-20 hours, the 5 mL cultures were added to 50 mL of TBII+kanamycin media in 250 mL baffled Erlenmeyer culture flasks, and incubated for 90 minutes at 37°C and 225 rpm. After the time elapsed, 55 $\mu$L of 1 M IPTG solution was added to each of the 55 mL cultures to induce expression. The flasks were incubated at 37°C and 225 rpm for an additional 2 hours before the temperature was lowered to 25°C where the flasks were incubated for 20-24 hours to promote sufficient expression. The following day, the cultures were transferred into 50 mL falcon tubes, and cell pellets were collected by centrifugation (4,198g, 8 min, 25°C). The cell pellets were stored at -80°C until purification. Note that all steps involving inoculation, media transfers, and IPTG additions were performed with sterile technique under flame.

##### *Protein purification for characterization*

The frozen cell pellets were thawed, lysed, and purified by (1) affinity chromatography with Strep-Tactin® resin and (2) size-exclusion chromatography. First, three buffers were prepared with Milli-Q water as follows: **W1 buffer** (100 mM Tris-HCl, 150 mM NaCl, 1 mM EDTA, pH 8.0), **W2 buffer** (100 mM Tris-HCl, 150 mM NaCl, pH 8.0), and E buffer (40 mM HEPES, 50 mM NaCl, 2.5 mM desthiobiotin, pH 8.0). Additionally, a **lysis buffer** was made from the W2 buffer where it was supplemented with 100 $\mu$g/mL lysozyme, 10 $\mu$g/mL of DNAse I, and 1 Pierce protease inhibitor tablet per 50 mL of W2 buffer.

The thawed cell pellets were resuspended in 20 mL of lysis buffer and mechanically lysed by ultrasonication (IBA Lifesciences) at 80% amplitude with 10s on/off pulse cycles for 5 minutes of total “on” time. Note that this was all performed in an ice bath to prevent significant heating of the lysate. The lysates were then centrifuged (4,198 g, 30 min, 4°C) to collect the supernatant.

Plastic 25 mL gravity flow-through chromatography columns were prepared with 6-10 mL of new Strep-Tactin® resin in a 50% suspension such that 1 CV = 3-5 mL of resin. The columns were equilibrated 2 times with 2.5 CV of W1 buffer followed by 1 addition of 1 CV of W2 buffer. Note that this step and the following steps for affinity chromatography were carried out in a 4°C room using only gravity flow-through. Once the columns were equilibrated, the lysate supernatant was gently added over the columns and allowed to completely flow through. The columns were then washed 3 times with 1 CV of W1 buffer, 2 times with 1 CV of W2 buffer, and 1 time with 0.5 CV of E buffer. Next, the bound protein in each column was eluted into a falcon tube with 5 additions of 0.5 CV of E buffer. The collected elutants were set aside for further purification by size-exclusion chromatography and the columns were washed 1 time with 2 CV of E buffer before being regenerated by 3 additions of 5 CV of regeneration buffer (obtained as 10x concentrated and diluted in Milli-Q water). Note that the resin was resuspended several times during regeneration so that the columns had a homogenous bright red color after regeneration; the regenerated columns were overlaid with 40-60% v/v ethanol and water to prevent bacterial growth, capped at both ends, and stored at 4˚C for future use.

The collected protein eluants were spin-concentrated to a final volume of 1.0∼1.1 mL using a 3 kDa MWCO centrifugal concentrator. The concentrated elutants were then purified by size-exclusion chromatography on an ÄKTA pure™chromatography system using a Superdex 75 Increase 10/300 GL column with a running buffer of 40 mM HEPES, 50 mM NaCl, pH 8.0. The resulting monomeric fractions of the purified samples were collected and immediately used in downstream experiments. Protein molecular weight was confirmed with electrospray ionization mass spectrometry.

***Kinetic analysis (general setup)***

The Michaelis-Menten kinetics of each purified design of interest were determined by incubating the protein samples with zinc sulfate and the fluorogenic substrate 4MU-PA (WuXi) or 4MU-B (Sigma) while monitoring the fluorescent signal of the product formation at 25˚C with an excitation/emission wavelength 365 nm and 445 nm, respectively. Before the kinetics, the purified protein samples were diluted to 10x of their desired concentration in the reaction using **reaction buffer** (40 mM HEPES, 50 mM NaCl, pH 8.0). Additionally, zinc sulfate dissolved in Milli-Q water was added to each of these 10x concentrated purified protein stocks to reach a desired [zinc]:[enzyme] ratio. Note that the numeric details for the concentrations of protein and zinc are described in a separate section for each reaction. The samples containing protein and zinc sulfate at 10x the reaction concentration were incubated at room temperature for several minutes. Additionally, stock solutions for the 4MU-PA and 4MU-B substrates and 4MU fluorescent product (Sigma) were made by dissolving each of the solids in DMSO.

Two sets of serial dilutions of the 4MU product (1.4 - 45 µM and 0.9 - 30 µM) were made in 95% reaction buffer and 5% DMSO to construct a fluorescence calibration curve, where each point was measured at 25˚C in a 96-well half area plate using 100 $\mu$L. Every point was measured in triplicate and averaged over 4 minutes of measurements. The calibration curve was fitted with a linear regression equation to convert fluorescence into the product concentration of 4MU. New calibration curves were made before every series of experiments.

After constructing a standard curve, serial dilutions of the substrate (specific concentrations given in each substrate section) were made in 94.5% v/v reaction buffer and 5.5% v/v DMSO at concentrations 1.11x the desired concentration in the reaction to account for a 0.9x dilution with the 10x concentrated protein-zinc sample. Kinetic reactions were performed by mixing 10 $\mu$L of the 10x concentrated protein-zinc sample with 90 $\mu$L of the prepared substrate solution in a 96-well half area plate. The reactions were allowed to shake for 3s at 282 cpm immediately after mixing the enzyme and substrate. Each reaction was monitored for 4 minutes at 25°C and all substrate concentration points were measured in triplicate. Note that the DMSO was 5.0% v/v in the final reaction mixture. The resulting curves were analyzed by first subtracting the fluorescence of the background reaction at each substrate concentration point for the matching zinc concentration in the reaction buffer. The resulting fluorescence curve was manually set to start at y=0 fluorescence units such that a linear regression ($y = mx + b$) could be calculated for the linear initial velocity with $b = 0$ (i.e., only the slope, m, needed to be determined). The calculated slopes for the initial velocities at each substrate concentration point were transformed to be expressed in terms of [4MU product]/[enzyme] using the calibration curve and known enzyme concentration. The steady-state parameters kcat and KM were determined by nonlinear regression fitting all data points to the Michaelis-Menten equation: $v_0/[E]_0 = k_{\text{cat}}[S]/(K_M + [S])$, where $v_0$ denotes the initial velocity, $[E]_0$ the enzyme concentration, and $[S]$ the substrate concentration. Standard deviations were calculated from the standard error of the fit considering each triplicate value as an individual data point.

##### *Kinetic analysis (4MU-PA)*

As mentioned in the kinetic analysis (general setup) section, serial dilutions of the 4MU-PA substrate (1.1 - 144 $\mu$M reaction concentration) were made in 94.5% v/v reaction buffer and 5.5% v/v DMSO at concentrations 1.11x the desired concentration in the reaction to account for a 0.9x dilution with the 10x concentrated protein-zinc sample. Kinetic experiments were then carried out as described in the kinetic analysis (general setup) section.

Regarding the hits from the 1st design campaign, note that the kinetics for A1 (ZETA_1) were carried out at $[E]_0$ = 100 nM, 40:1 [Zn(II)]:$[E]_0$ (4 µ$\mu$M zinc sulfate in reaction), and 1.1 - 72 $\mu$M 4MU-PA substrate. The kinetics for C4 were carried out at $[E]_0$ = 3 $\mu$M, 10:3 [Zn(II)]:$[E]_0$ (10 $\mu$M zinc sulfate in reaction) due to precipitation of this design above this zinc concentration, and 1.1 - 144 $\mu$M 4MU-PA substrate. The kinetics for all of the other characterized hits (A8, B9, F7) were carried out at $[E]_0$ = 2$\mu$M, 10:1 [Zn(II)]:$[E]_0$ (20 $\mu$M zinc sulfate in reaction), and 1.1 - 144 $\mu$M 4MU-PA substrate. Experiments were performed in biological duplicates, with the results being in close agreement. We herein report representative results for each design/variant.

Regarding the best design from each of the 3 unique, highly active scaffold hits in the 2nd design campaign, note that the kinetics for ZETA_2 (well position H10) and ZETA_3 (well position H6) were carried out at $[E]_0$ = 100 nM, 10:1 [Zn(II)]:$[E]_0$ (1 $\mu$M zinc sulfate in reaction), and 1.1 - 72 $\mu$M 4MU-PA substrate. The kinetics for ZETA_4 (well position C5) were carried out at $[E]_0$ = 500 nM, 10:1 [Zn(II)]:$[E]_0$ (5 $\mu$M zinc sulfate in reaction), and 1.1 - 72 $\mu$M 4MU-PA substrate. Experiments were performed in biological duplicates, with the results being in close agreement. We herein report representative results for each design/variant.

##### *Kinetic analysis (4MU-B)*

As mentioned in the kinetic analysis (general setup) section, serial dilutions of the 4MU-B substrate (0.77 - 99 µM reaction concentration) were made in 94.5% v/v reaction buffer and 5.5% v/v DMSO at concentrations 1.11x the desired concentration in the reaction to account for a 0.9x dilution with the 10x concentrated protein-zinc sample. Kinetic experiments were then carried out as described in the kinetic analysis (general setup) section.

Note that the kinetics for all 4MU-B hits (C4, B11, D11) were carried out at $[E]_0 = 3\mu$M, 20:3 [Zn(II)]:$[E]_0$ (20 $\mu$M zinc sulfate in reaction), and 0.7 - 99 $\mu$M 4MU-B substrate. Experiments were performed in biological duplicates, with the results being in close agreement. We herein report representative results for each design/variant.

Note that for the design D11, the Michaelis-Menten plot could not resolve $k_{\text{cat}}$ or KM because the substrate conditions were unable to saturate the enzyme, indicating a high KM value; we could not further increase the substrate concentration without the 4MU-B beginning to precipitate (the mixture became increasingly turbid >100 $\mu$M). Thus, we only report the estimated $k_{\text{cat}}/K_M$ value, which was obtained from the Michaelis-Menten equation assuming sub-saturating conditions (i.e., $[S] \ll K_M$): $v_0/[E]_0 = k_{\text{cat}}/K_M$ for a single given substrate concentration. Thus, we measured the initial velocity for each substrate concentration and used them, divided by the known enzyme concentration, each as a single point estimate for $k_{\text{cat}}/K_M$ and averaged them to obtain our reported estimate for the $k_{\text{cat}}/K_M$ of D11.

##### *Electrospray ionization (ESI) mass spectrometry*

MS data for the identified hits and A1 mutants were acquired on an Agilent 1200series LC G6230B TOF LC-MS with an AdvanceBio RP-Desalting column (A: H2O with 0.1% Formic Acid, B: Acetonitrile with 0.1% Formic Acid). Final protein concentrations were adjusted to 1-2 mg/mL in 40 mM HEPES, 50 mM NaCl, pH 8.0. Subsequent data deconvolution was performed in Bioconfirm using a total entropy algorithm. All electrospray ionization mass spectrometry results for the characterized hits are presented below with an error of approximately ±1 Da. In nearly every case, the observed mass corresponded to the loss of N-terminal methionine (-131 Da).

| **Design campaign** | **Design** | **E**: Expected Mass (Da) | **O**: Observed Mass (Da) | Δ(O - E) |
|---|---|---|---|---|
| 4MU-PA (1st Design Campaign) | ZETA_1 (A1) | 18528 | 18397 | -131 |
| | A8 | 25301 | 25170 | -131 |
| | B9 | 20202 | 20071 | -131 |
| | C4 | 18724 | 18593 | -131 |
| | F7 | 21023 | 20892 | -131 |
| 4MU-PA (2nd Design Campaign; Top Design from 3 Unique Scaffold Hits) | ZETA_2 (H10) | 15748 | 15616 | -132 |
| | ZETA_3 (H6) | 22046 | 21915 | -131 |
| | ZETA_4 (C5) | 22585 | 22454 | -131 |
| 4MU-B | C4 | 29398 | 29398 | 0 |
| | B11 | 21832 | 21701 | -131 |
| | D11 | 27332 | 27201 | -131 |
| Cysteine Hydrolase | Eta1 | 18465 | 18334 | -131 |

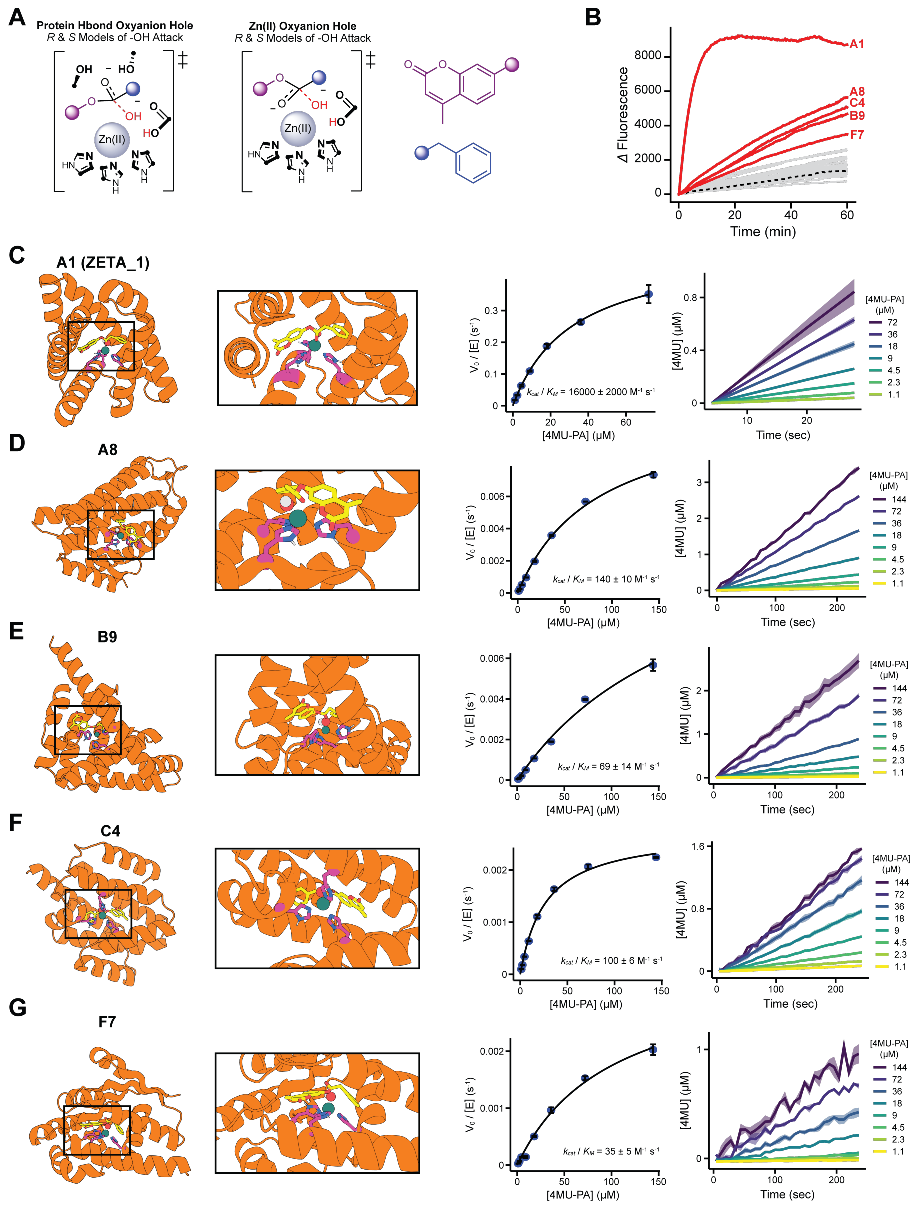

**Figure 17**: Results of the 1st metallohydrolase design campaign for 4MU-PA. More details for these results are available at [50]. **(A)** 2D schematic of the 4 classes of theozymes considered in this design campaign. The classes are first divided by the nature of the oxyanion stabilization (i.e., oxyanion hole), and then further divided into the R- or S- enantiomer of the hydroxide attack on the substrate. Black dots represent points of protein connectivity to the side chains (or options of protein connectivity for the His imidazole rings). **(B)** Reaction progress curves from the screening of the 96 ordered designs from 96 unique backbones. The 5 identified and characterized hits are labeled in red. **(C-G)** Design models with magnified active site panels (left). Corresponding Michaelis-Menten plots, $k_{\text{cat}}/K_M$ values, and activity traces (right).

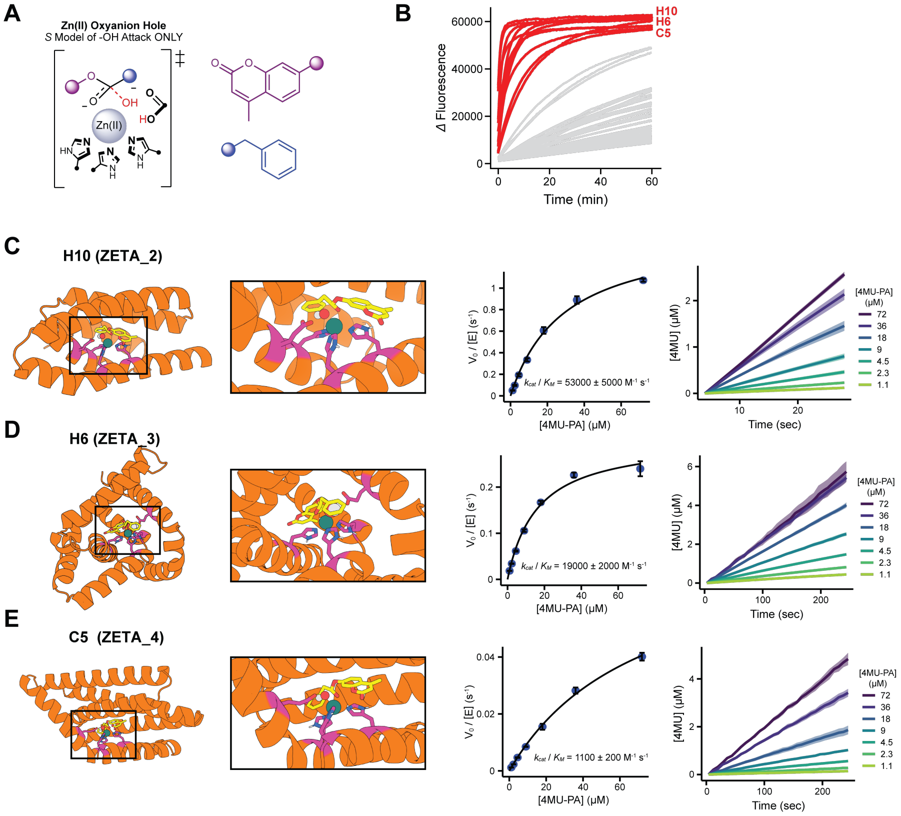

**Figure 18**: Results of the 2nd metallohydrolase design campaign for 4MU-PA. More details for these results are available at [50]. **(A)** 2D schematic of the 1 class of theozymes considered in this design campaign. Note that all of the His imidazole rings are forced to coordinate Zn(II) with their N-epsilon atom (as opposed to N-delta) in this design campaign, which was not a requirement in the first design campaign. Additionally, only the S- enantiomer of the hydroxide attack was considered this time. **(B)** Reaction progress curves from the screening of the 96 ordered designs from 37 unique backbones. The 11 identified and characterized hits are labeled in red, coming from 3 unique scaffold families. **(C-E)** Design models of the best enzyme (named ZETA_2-4) from each of the 3 unique scaffolds with magnified active site panels (left). Corresponding Michaelis-Menten plots, $k_{\text{cat}}/K_M$ values, and activity traces (right). Note that all 11 hits from these 3 unique scaffolds have measured $k_{\text{cat}}/K_M$ values $> 103$ $M^{-1}$ $s^{-1}$, shown in [50].

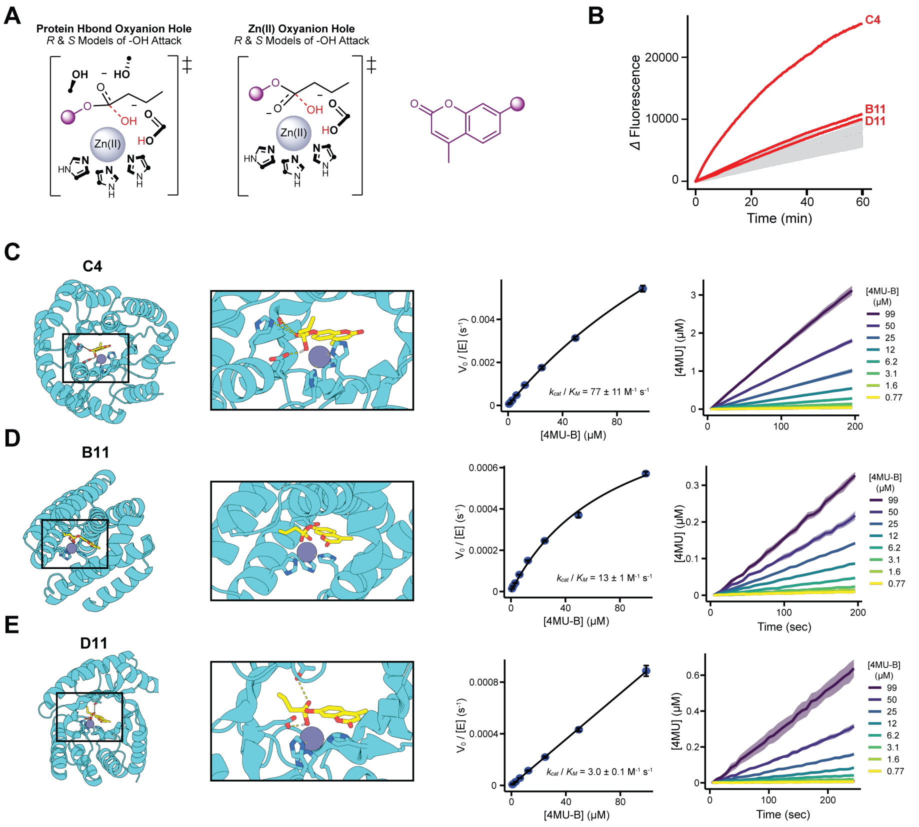

**Figure 19**: Results of the metallohydrolase design campaign for 4MU-B. **(A)** 2D schematic of the 4 classes of theozymes considered in this design campaign. The classes are first divided by the nature of the oxyanion stabilization (i.e., oxyanion hole), and then further divided into the R- or S- enantiomer of the hydroxide attack on the substrate. Black dots represent points of protein connectivity to the sidechains (or options of protein connectivity for the His imidazole rings). **(B)** Reaction progress curves from the screening of the 96 ordered designs from 68 unique backbones. The 3 identified and characterized hits are labeled in red. Each hit came from a unique scaffold. **(C-G)** Design models with magnified active site panels (left). Corresponding Michaelis-Menten plots, $k_{\text{cat}}/K_M$ values, and activity traces (right).

#### D.4 Structural drift during sequence fitting

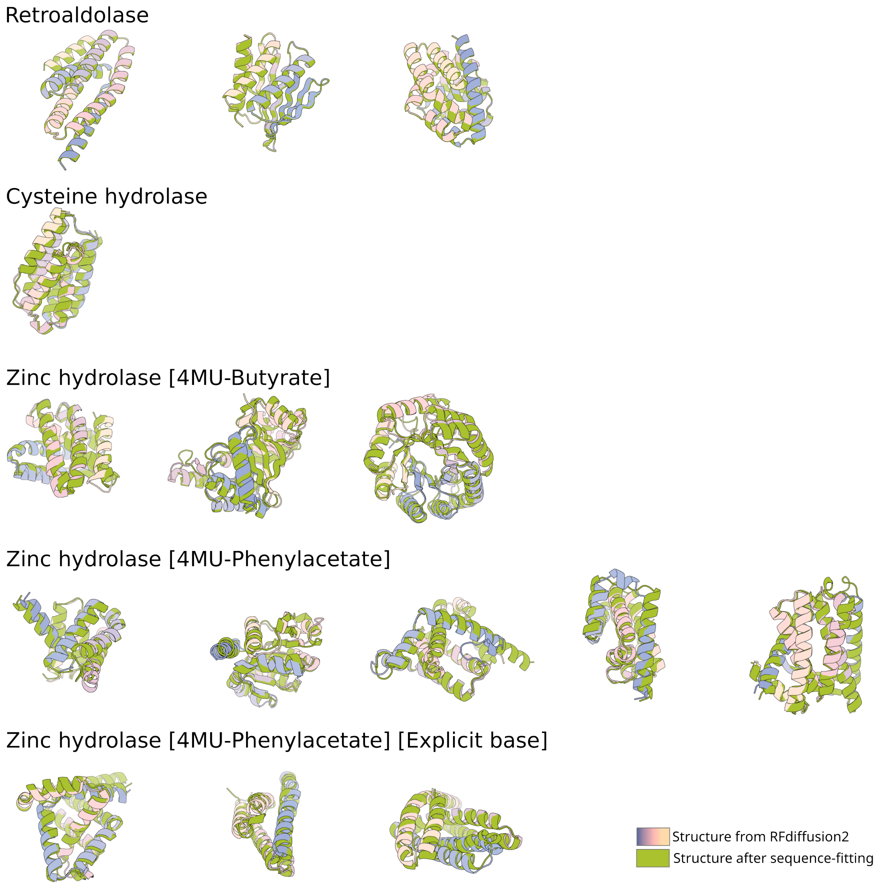

**Figure 20**: **Scale of structural differences induced by sequence fitting** The process of fitting sequence to the structures generated by RFdiffusion2 produces small structural differences from the original design. We show the TM alignment of each unique RFdiffusion2-generated structure that resulted in experimental success to its corresponding structure after from sequence fitting to demonstrate the scale of these relaxations.
